# A novel quinone biosynthetic pathway illuminates the evolution of aerobic metabolism

**DOI:** 10.1101/2024.07.27.605408

**Authors:** Felix J. Elling, Fabien Pierrel, Sophie-Carole Chobert, Sophie S. Abby, Thomas W. Evans, Arthur Reveillard, Ludovic Pelosi, Juliette Schnoebelen, Jordon D. Hemingway, Ahcène Boumendjel, Kevin W. Becker, Pieter Blom, Julia Cordes, Vinitra Nathan, Frauke Baymann, Sebastian Lücker, Eva Spieck, Jared R. Leadbetter, Kai-Uwe Hinrichs, Roger E. Summons, Ann Pearson

**Affiliations:** Department of Earth and Planetary Sciences, Harvard University, Cambridge, MA 02138, USA; Leibniz-Laboratory for Radiometric Dating and Isotope Research, Christian-Albrecht University of Kiel, 24118 Kiel, Germany; Univ. Grenoble Alpes, CNRS, UMR 5525, VetAgro Sup, Grenoble INP, TIMC, 38000 Grenoble, France; Department of Earth, Atmospheric, and Planetary Sciences, Massachusetts Institute of Technology, Cambridge, MA 02139, USA; MARUM – Center for Marine Environmental Sciences and Department of Geosciences, University of Bremen, 28359 Bremen, Germany; Geological Institute, Department of Earth and Planetary Sciences, ETH Zurich, 8092 Zurich, Switzerland; Univ. Grenoble Alpes, INSERM, LRB, Grenoble, 38000, France; GEOMAR Helmholtz Centre for Ocean Research Kiel, 24148 Kiel, Germany; Department of Microbiology, Radboud Institute for Biological and Environmental Sciences, Radboud University, 6525 AJ Nijmegen, The Netherlands; Laboratoire de Bioénergétique et Ingénierie des Protéines UMR 7281 CNRS/AMU, FR3479, F-13402 Marseille Cedex 20, France; Department of Microbiology and Biotechnology, University of Hamburg, 22609 Hamburg, Germany; Division of Geological and Planetary Sciences, California Institute of Technology, Pasadena, CA 91125, USA; Division of Engineering and Applied Science, California Institute of Technology, Pasadena, CA 91125, USA

## Abstract

The dominant organisms in modern oxic ecosystems rely on respiratory quinones with high redox potential (HPQs) for electron transport in aerobic respiration and photosynthesis. The diversification of quinones, from low redox potential in anaerobes to HPQs in aerobes, is assumed to have followed Earth’s surface oxygenation ∼2.3 billion years ago. However, the evolutionary origins of HPQs remain unresolved. Here, we characterize the structure and biosynthetic pathway of a novel ancestral HPQ, methyl-plastoquinone, that is unique to bacteria of the phylum *Nitrospirota*. Methyl-plastoquinone is structurally related to the two previously known HPQs, plastoquinone from *Cyanobacteriota*/chloroplasts and ubiquinone from *Pseudomonadota*/mitochondria, respectively. We demonstrate a common origin of the three HPQ biosynthetic pathways that predates the emergence of *Nitrospirota, Cyanobacteriota*, and *Pseudomonadota*. An ancestral HPQ biosynthetic pathway evolved ≥ 3.4 billion years ago in an extinct lineage and was laterally transferred to these three phyla ∼2.5-3.2 billion years ago. We show that *Cyanobacteriota* and *Pseudomonadota* were ancestrally aerobic and thus propose that aerobic metabolism using HPQs significantly predates Earth’s surface oxygenation. Two of the three HPQ pathways were later obtained by eukaryotes through endosymbiosis forming chloroplasts and mitochondria, enabling their rise to dominance in modern oxic ecosystems.

**Significance statement:** Oxygenic photosynthesis and aerobic respiration by bacteria and eukaryotes rely on respiratory quinones with high redox potential that facilitate membrane-bound electron transport. These quinones are integral to aerobic metabolism and therefore the evolution of aerobic metabolism and quinone biosynthesis must be intertwined. Only two types of high redox potential quinones have been described in bacteria and eukaryotes. Here, we describe the structure and biosynthetic pathway of a third type, methyl-plastoquinone, that is exclusive to bacteria of the phylum *Nitrospirota*. We then use phylogenetic analysis to show that the three high redox potential quinones have a single evolutionary origin and are much older than previously considered, predating the Great Oxygenation Event, when significant amounts of O_2_ first accumulated in the atmosphere.

## Introduction

The oxygenation of Earth’s surface environments following the emergence of oxygenic photosynthesis in ancestors of *Cyanobacteriota* enabled the metabolic and genetic diversification of life (1–4). The use of oxygen as a terminal electron acceptor, i.e., aerobic respiration, enabled a higher energy yield compared to anaerobic metabolisms and was a prerequisite for the emergence of eukaryotes (5, 6). However, it remains poorly resolved how and when the electron transport chain (ETC) used for aerobic respiration evolved. While geochemical evidence indicates iron oxidation by acidophilic bacteria must have evolved by the time oxygen accumulated in the atmosphere during the great oxygenation event (GOE; ∼2.4-2.3 Ga) (3, 7–9), there is now considerable evidence for an ancient origin of dioxygen-utilizing and detoxifying enzymes as early as 3.1 Ga (10–13). Though these enzymes may not have participated in aerobic respiration (11, 14, 15), their widespread occurrence in bacteria suggests the availability of oxygen in physiologically significant quantities, at least in some niches, before the GOE. Studying the evolution of ETC components, such as oxygen reductases that use electrons derived from the ETC (16–21), can help elucidate the origins of aerobic metabolisms. However, the interpretation of oxygen reductase evolution has remained contentious (16–20, 22), and alternative roles of ancestral oxygen reductases in oxygen detoxification and nitric oxide reduction rather than aerobic respiration have been proposed (17). Exploring the evolution of other ETC components, such as respiratory quinones, may yield new insights into the evolution of ETCs and aerobic respiration.

Strict anaerobes use ETCs and quinones with low redox potential (LPQs), while aerobes and facultative aerobes generally use high-potential quinones (HPQs) (23–25). HPQs require all parts of the ETC to operate at high redox potential (25–27) and confer no known benefit over LPQs under anaerobic conditions. However, under aerobic conditions HPQs are advantageous due to their decreased electron leakage to oxygen, thus reducing oxidative stress and minimizing free energy losses (26, 28). The occurrence of HPQs may represent a marker for high-potential ETCs and their evolution may be tied to the history of oxygenic photosynthesis and aerobic respiration. Within bacteria, HPQs have been found only in two phyla, oxygenic *Cyanobacteriota* (here used *sensu stricto*, including only *Cyanophyceae*) and *Pseudomonadota* (formerly Proteobacteria, now comprising the classes *Alpha*-, *Beta*-, *Gammaproteobacteria, Acidithiobacillia*, and *Hydrogenophilia*) (29). The *Cyanobacteriota* and *Pseudomonadota* produce two distinct types of HPQs, plastoquinone (PQ) and ubiquinone (UQ), respectively (23, 24), which became the quinones of plastids (PQ) and mitochondria (UQ) through endosymbiosis during the early evolution of eukaryotes (24, 30). Yet, despite the dominance of HPQ-utilizing organisms in Earth’s oxic environments today (31–36), the co-evolution of HPQs and Earth surface oxygenation remains largely unresolved (37).

Recent progress in metagenomic coverage of uncultivated bacteria and isolation of novel lineages may help elucidate HPQ evolution through the discovery of new quinone structures and biosynthetic pathways in unstudied lineages of aerobic bacteria. Here, we describe the discovery of a third, novel type of HPQ, methyl-plastoquinone (mPQ). mPQ occurs only in aerobic members of the phylum *Nitrospirota* (formerly Nitrospirae), a metabolically diverse group of bacteria that perform essential transformations in the biogeochemical cycles of iron, nitrogen, and manganese. We characterize the biosynthetic pathway of mPQ using bioinformatic, genetic, and biochemical techniques and use these data to infer the evolutionary history of HPQs. Our study sheds new light on the evolutionary history of ETCs by revealing a single origin of the three HPQ biosynthetic pathways prior to the radiation of crown-group *Cyanobacteriota, Nitrospirota*, and *Pseudomonadota*, which evidently preceded the GOE.

## Results & Discussion

### Novel respiratory quinones in Nitrospirota

Despite their widespread distribution and the important roles of *Nitrospirota* in biogeochemical cycles of iron, manganese, and nitrogen (38–42), many aspects of their chemotaxonomy and bioenergetics remain understudied. Genome-based bioenergetic models implicate the presence of ETCs in aerobic and anaerobic *Nitrospirota* (42–46), yet their corresponding respiratory quinones have not been studied. During screening of *Nitrospirota* genomes for lipid biosynthetic pathways (47), we observed that the genomes of aerobic *Nitrospirota* did not contain any of the characterized quinone biosynthesis pathways (24, 37). In contrast, genomes of anaerobic sulfur-reducing *Nitrospirota*, i.e., *Thermodesulfovibrio* species and some *Nitrospirota* metagenome-assembled genomes from anoxic environments, contained the futalosine pathway (MK_mqn_, composed of *mqn* genes) for biosynthesis of the LPQ menaquinone (MK; Supplementary Datafile S1).

To evaluate the presence of respiratory quinones, we analyzed lipid extracts of one anaerobic and eight aerobic species of *Nitrospirota*, covering all formally described genera (*Thermodesulfovibrio, Leptospirillum, Nitrospira*, and *Candidatus* Manganitrophus), using high-performance liquid chromatography coupled to high-resolution tandem mass spectrometry. MKs were detected only in the anaerobic species, *Thermodesulfovibrio islandicus* (Fig. 1a), and we did not find any of the previously known respiratory quinone types in the aerobic *Nitrospirota*. Instead, all eight studied aerobic *Nitrospirota* contained a novel type of quinone, identified as methyl-plastoquinone (mPQ). The polyprenyl chain of mPQ varied in length and saturation depending on species (Fig. 1, Fig. S1, SI results). Mass spectrometric characterization of mPQ revealed fragmentation spectra analogous to PQ but with a dominant ion at *m*/*z* 165 instead of 151, indicative of a distinctive trimethyl-benzoquinone headgroup connected to the isoprenoid tail (Fig. 1, Fig. S1, Table S1). Stable isotope labeling experiments and nuclear magnetic resonance spectroscopy of mPQ confirmed the structural assignment of the headgroup (SI Results & Discussion; Fig. S2-4). Specifically, ^1^H-NMR spectra showed the absence of any proton linked to the C2 of the quinone moiety and ^1^H-NMR and ^13^C-NMR confirmed the presence of a third methyl group (Fig. S4; see SI Results & Discussion). mPQ is thus structurally related to both UQ (methylated at C2 of the benzoquinone) and PQ (methylated at C5 and C6).

**Fig. 1.**
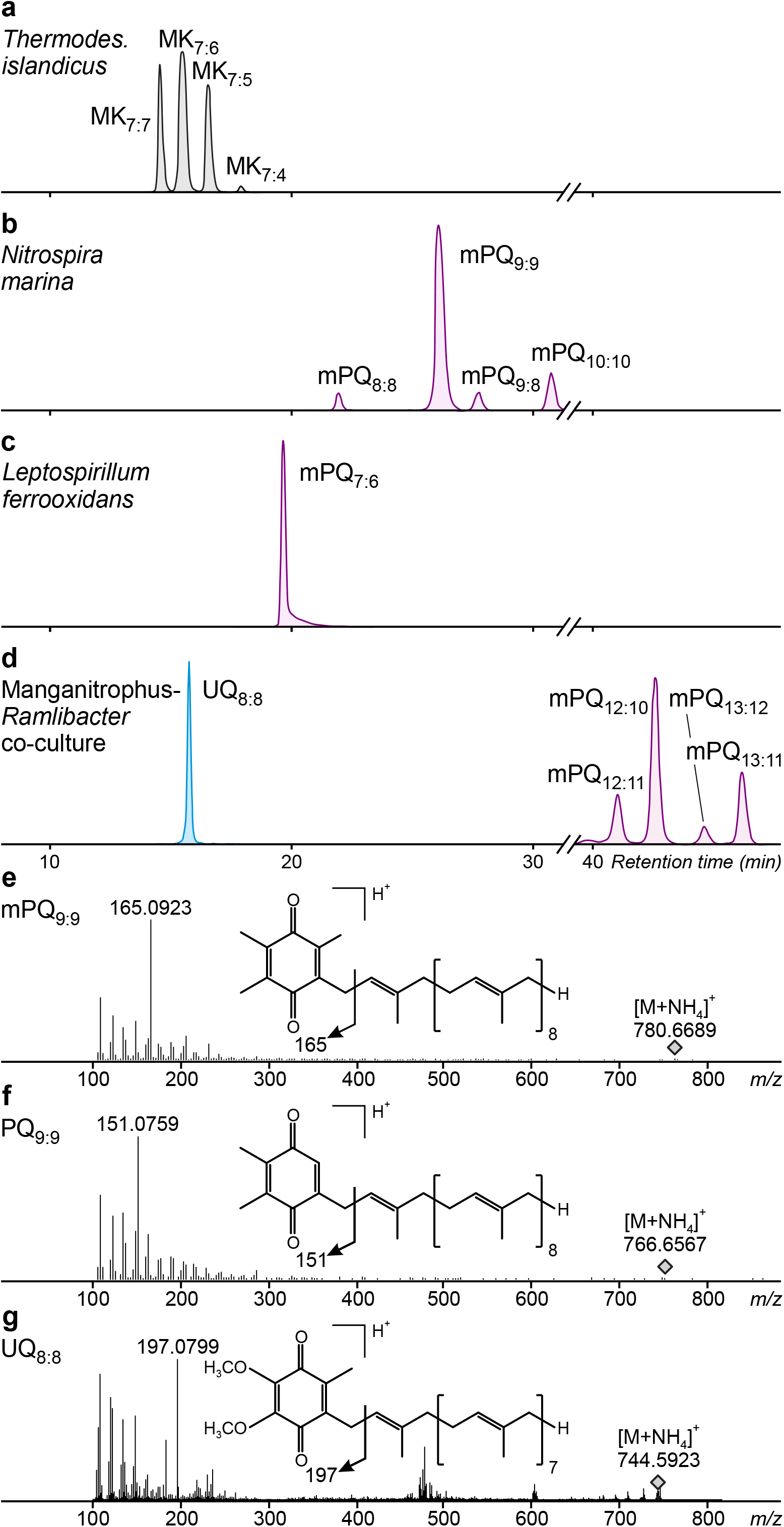
Novel quinones detected in aerobic *Nitrospirota*. **a-d**, Chromatograms showing presence of a distinct quinone type (methyl-plastoquinone, mPQ) in aerobic *Nitrospirota* (*Nitrospira marina, Leptospirillum ferrooxidans, Ca*. Manganitrophus noduliformans) and canonical menaquinones (MK) in the anaerobic *Nitrospirota* species *Thermodesulfovibrio islandicus*. Ubiquinone (UQ_8:8_) in the *Ca*. Manganitrophus*-Ramlibacter* co-culture derives from *Ramlibacter* (see Fig. S1). **e-g**, High resolution mass spectrometric characterization of mPQ_9:9_ and PQ_9:9_ showing similar fragmentation patterns but suggesting the presence of a trimethyl-benzoquinone moiety in mPQ_9:9_ (see Fig. S1); structure and fragmentation pattern of UQ_8:8_ from *Ramlibacter* shown in **g** for reference.

### Characterization of the biosynthetic pathway of mPQ

Based on the structure of mPQ, we hypothesized that its biosynthesis pathway might share characteristics with the UQ and PQ biosynthesis pathways. The PQ biosynthesis pathway of *Cyanobaceriota* has been partially resolved (48, 49) and contains several enzymes that are homologous to enzymes involved in the well-characterized bacterial UQ pathway (37, 50). In both pathways, the conversion of chorismate into 4-hydroxybenzoate (4-HBA) is mediated by a UbiC homolog and the subsequent prenylation, decarboxylation, and hydroxylation of 4-HBA involve the UbiA, UbiD/X, and UbiH homologs, respectively (Fig. 2a) (37). Specific to the UQ pathway, methylation at C2 is mediated by UbiE (51). In the cyanobacteriotal PQ pathway, methylation at C5 and C6 has been proposed to be mediated by Sll0418 (PlqQ) (50, 52).

**Fig. 2.**
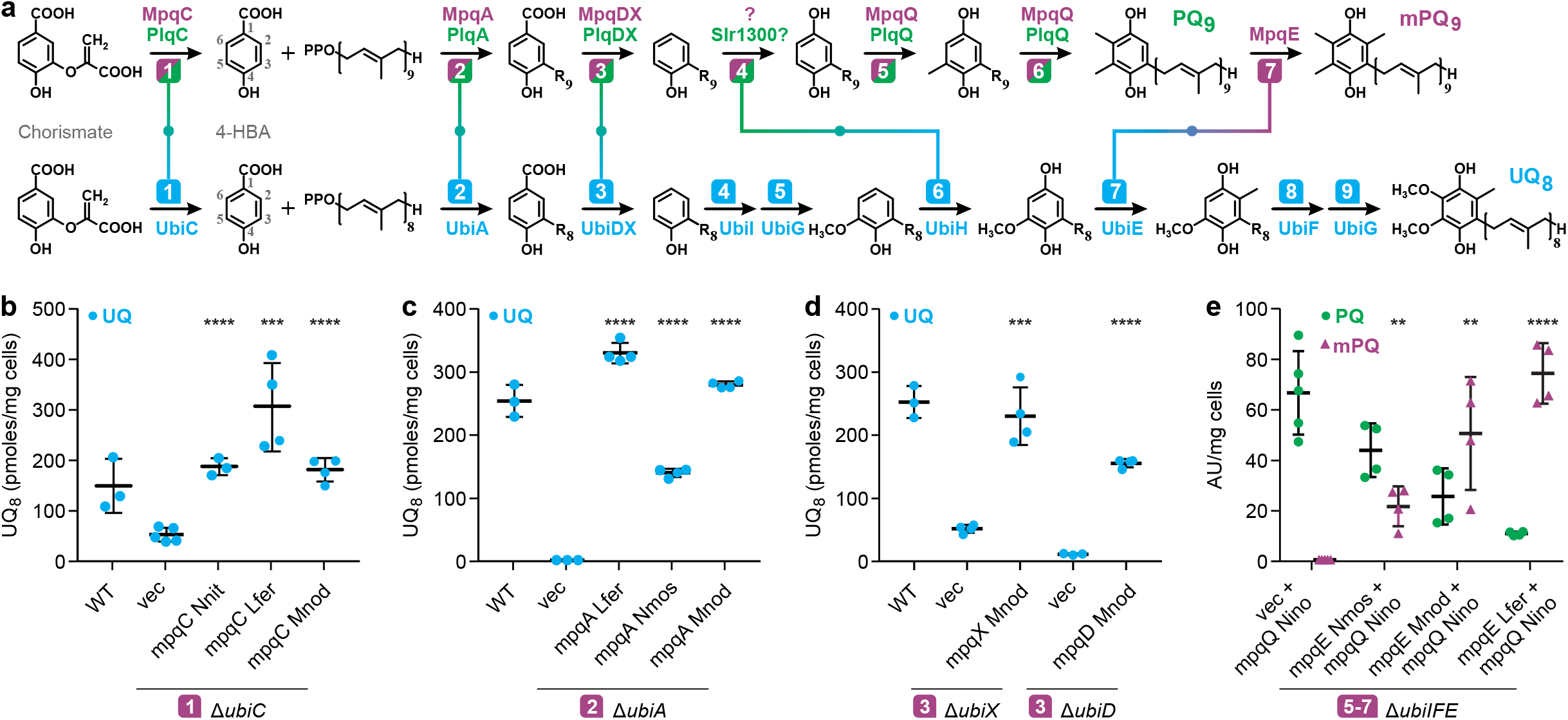
Characterization of the mPQ biosynthetic pathway. **a**, Biosynthetic pathways of quinones showing homology of pathways for mPQ_9_ in *Nitrospirota* (purple), PQ_9_ in the cyanobacterium *Synechocystis sp*. PCC6803 (green) and UQ_8_ in the gammaproteobacterium *Escherichia coli* (blue). Biosynthetic steps are numbered, and homologous steps are connected by colored lines. **b-d**, Heterologous complementation experiments using mPQ biosynthesis gene candidates to restore UQ_8_ production in *E. coli* mutants lacking key genes for ubiquinone biosynthesis (Δ*ubiC*+*mpqC*, Δ*ubiA*+*mpqA*, Δ*ubiX*+*mpqX*, Δ*ubiD*+*mpqD*). **e**, PQ production in *E. coli* Δ*ubiIFE* mutants complemented with *mpqQ* from *N. inopinata* as well as PQ and mPQ in *E. coli* Δ*ubiIFE* mutants complemented with *mpqQ* from *N. inopinata* and *mpqE* from other *Nitrospirota*. WT=wild type; vec=empty vector; thick bars represent means and error bars represent standard deviations of the means, *n*=3-5; AU=arbitrary units. Abbreviations: *Ca*. N. nitrificans (Nnit), *N. moscoviensis* (Nmos), *N. inopinata* (Nino), *L. ferrooxidans* (Lfer), *Ca*. M. noduliformans (Mnod). The numbering of the carbon atoms on the 4-HBA precursor (panel **a**, light grey) defines the nomenclature for all intermediates described in the text. The octaprenyl and nonaprenyl chains are abbreviated with R_8_ and R_9_, respectively. See Fig. S7-S9 for details on compound identification and quantification. Stars indicate *p* < 0.01 (**), *p* < 0.001 (***), and *p* < 0.0001 (****) for unpaired Student’s *t* tests relative to the empty vector.

Isotope labeling experiments further point to biochemical similarities between the HPQ biosynthesis pathways. Supplementation of cultures with ring-^13^C_6_-labeled substrates demonstrates that 4-HBA is the ring precursor in *Nitrospirota*, similar to *Cyanobacteriota* and *Pseudomonadota* (Fig. S5, SI Discussion). Further, experiments with methyl-^2^H_3_ methionine indicate that all three methyl groups of mPQ (at C2, C5, and C6) are derived from methionine via SAM-dependent methyltransferases (Fig. S2). We used this information to find multiple, homologous candidate genes for the biosynthetic pathway of mPQ in genomes of *Nitrospirota*. We suggest a gene nomenclature for the mPQ pathway (*mpq*) analogous to that of the UQ pathway and extend this to the PQ pathway (*plq*; Fig. 2). We identified a four-gene cluster in *Leptospirillum* spp. (Fig. S6), encoding a *ubiA* family prenyltransferase (*mpqA*; LFE_2122), *ubiC*-like chorismate pyruvate lyase (*mpqC*; LFE_2123), a cobalamin-binding radical *S*-adenosyl methionine (SAM) methyltransferase (LFE_2124), and a *ubiE*-like methyltransferase (*mpqE*; LFE_2125). The genes are not co-localized in other *Nitrospirota*, but *mpqA* and *mpqE* homologs are found in all aerobic *Nitrospirota*, in addition to a *ubiB*-like kinase (*mpqB*). By contrast, homologs of *mpqC, ubiD/X* (*mpqD/X*) and *plqQ* (*mpqQ*) occur only in a subset of aerobic *Nitrospirota* (Table S2). No clear *ubiH* homologs were identified. Consequently, aerobic *Nitrospirota* contain a mosaic pathway for mPQ biosynthesis composed of well-conserved (*mpqA, mpqB, mpqE*) and alternative genes (*mpqC, mpqD/X, mpqQ*).

Due to the lack of suitable genetic systems in *Nitrospirota*, we verified the mPQ candidate genes by assessing their functions in heterologous complementation assays using *Escherichia coli* mutants deficient in defined steps of UQ biosynthesis. When expressed in the *E. coli* Δ*ubiC* mutant, the *mpqC* from *L. ferrooxidans, Ca*. N. nitrificans, and *Ca*. M. noduliformans restored UQ biosynthesis up to wild-type levels (Fig. 2b). Likewise, the *mpqA* homologs from *L. ferrooxidans, N. moscoviensis*, and *Ca*. M. noduliformans restored UQ biosynthesis in an *E. coli* Δ*ubiA* mutant (Fig. 2, S7). Similarly, we observed recovery of UQ levels in *E. coli* Δ*ubiD* and Δ*ubiX* mutants upon expression of *mpqD/X* from *Ca*. M. noduliformans (*mpqD/X* do not occur in *Nitrospira* and *Leptospirillum* spp.; Fig. 2). Expression of the *plqQ* homolog (*mpqQ*) from *N. inopinata* in an *E. coli* Δ*ubiIF* mutant yielded PQ_8_ and mPQ_8_ (Fig. S8b-g). The Δ*ubiIFE* strain, in which the *E. coli ubiE* gene was additionally deleted, showed a strong increase in the amount of PQ_8_ and the disappearance of mPQ_8_ (Fig. 2e, S8b-g). Finally, expression of *mpqE* from *L. ferrooxidans, N. moscoviensis*, and *Ca*. M. noduliformans in the Δ*ubiIFE* strain led to the accumulation of mPQ_8_ (Fig. 2e, S8c).

Based on these heterologous expression and isotope labeling experiments, we reconstructed a tentative mPQ biosynthetic pathway that shares homology with the UQ and PQ pathways (Fig. 2). The ring precursor 4-HBA is generated from chorismate by MpqC and alternative enzymes, followed by prenylation of 4-HBA by MpqA and decarboxylation by MpqD/X. The following hydroxylation step at C1 is unresolved, but observations from *Pseudomonadota* indicate that a large diversity of benzoquinone C1 hydroxylases exist in nature (37, 53–55). Finally, methylations are introduced at C5 and C6 by MpqQ and at C2 by MpqE.

### Distribution and function of mPQ

Analysis of mPQ biosynthesis proteins in a representative selection of high- and medium-quality genomes and metagenome-assembled genomes revealed that mPQ is present in all aerobic lineages of *Nitrospirota* (*n*=85), but not found outside this phylum (*n*=482). A few early-branching lineages of *Nitrospirota*, which are anaerobes using the MK_mqn_ pathway, are devoid of mPQ biosynthesis proteins (Fig. 3). Since mPQ is the only respiratory quinone detected in aerobic *Nitrospirota*, it is likely involved in the ETC used for aerobic respiration (42–44), and the structural similarity between mPQ and UQ/PQ suggests that mPQ has a high redox potential. Since *Nitrospirota* grow slowly and to low cell densities, mPQ could not be isolated in quantities required for redox potential measurements. We therefore calculated the redox potential of mPQ, UQ, and PQ using density functional theory (56). For a given biologically relevant prenyl chain length, the calculated redox potential of mPQ (E^0^(Q/H_2_Q) = 517±8 mV) is lower than that of PQ (551±8 mV) but higher than that of UQ (480±8 mV; Table S3). Furthermore, all HPQs are described by significantly higher calculated redox potentials than the LPQ MK (364±8 mV), confirming the validity of our computational approach. Calculations for simple 1,4-benzoquinones indicate that redox potentials decrease by *∼*50 mV per methyl or methoxy group, with methoxy additions having the larger effect, which explains the higher potential of mPQ (trimethyl) relative to UQ (dimethoxy, methyl). These functional group combinations may reflect redox tuning of HPQs to specific components of the ETC in *Cyanobacteriota, Nitrospirota*, and *Pseudomonadota*. Due to the tight coupling of redox potentials of quinones to other ETC components (e.g., iron sulfur clusters of Rieske proteins and *b*-hemes in Rieske/cyt*b* complexes) (27), we infer that aerobic *Nitrospirota* have high potential ETCs. Indeed, we find that Rieske proteins of aerobic *Nitrospirota* contain the ‘SY’ motif (Table S4) characteristic for Rieske/cytochrome *b* complexes adapted to interact with HPQs in high potential ETCs (57). High potential ETCs would be advantageous for minimizing ROS generation and maximizing proton motive force (27) in the low energy-yielding chemoautotrophic metabolisms of aerobic *Nitrospirota*. Since the use of HPQs requires adaptation of the entire ETC to higher redox potential (27), such a decisive step may have been linked to a major event, such as Earth’s surface oxygenation (25, 27, 58).

**Fig. 3.**
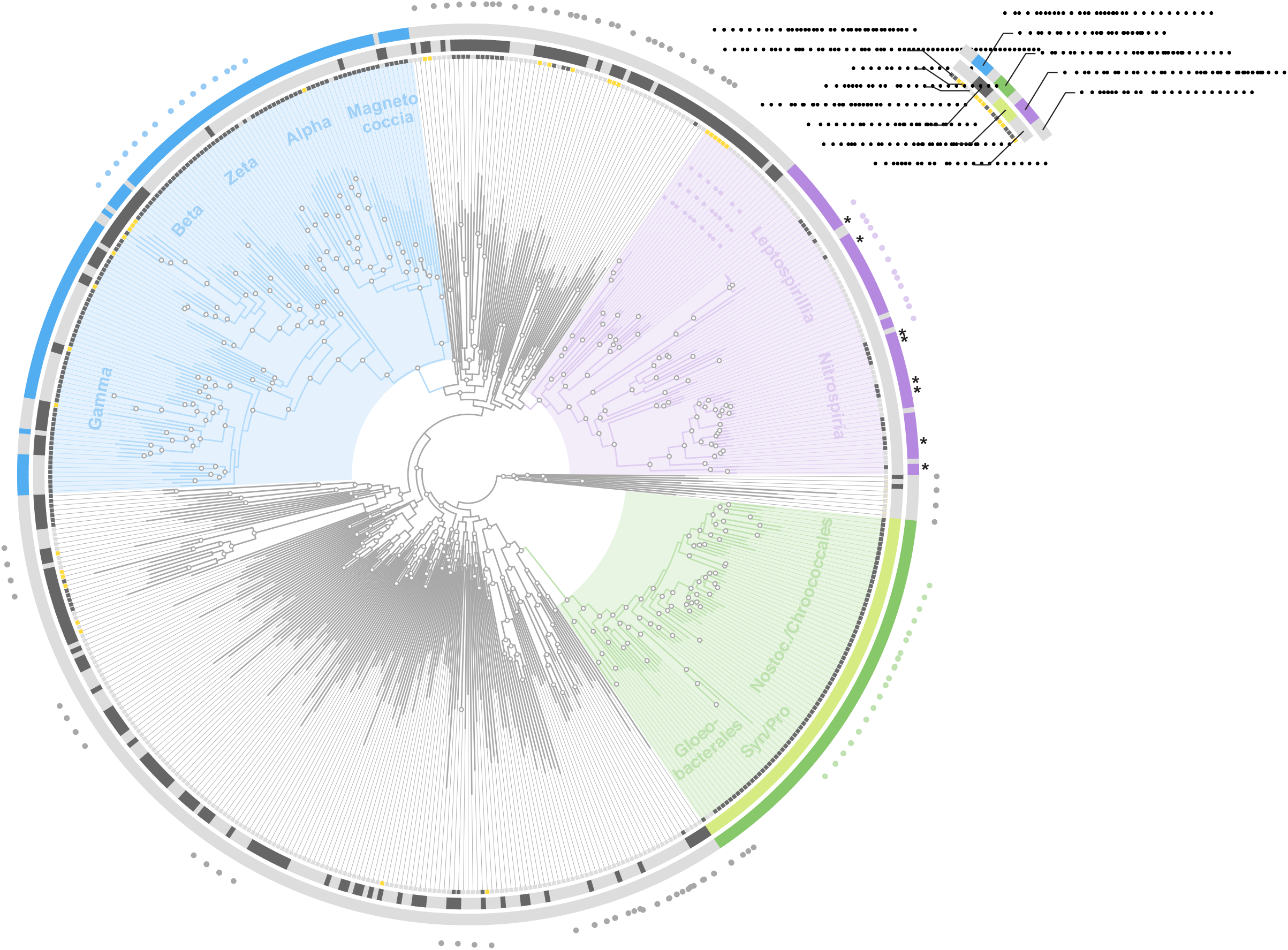
Phylogenetic tree of bacteria showing the occurrence of respiratory quinones. Quinones with high redox potential (UQ, PQ, mPQ) occur only in aerobic *Nitrospirota, Pseudomonadota*, and *Cyanobacteriota*. Low potential quinones occur in anaerobic *Nitrospirota* (MK), some *Pseudomonadota* (MK), and all *Cyanobacteriota* (PhQ). Asterisks indicate strains in which presence of mPQ has been verified experimentally. See Fig. S11-13 for detailed trees. The maximum-likelihood phylogenetic tree was constructed from 120 concatenated single copy marker proteins (59) of 547 isolate genomes and metagenome-assembled genomes, covering all bacterial phyla, and rooted using the DST group to approximate the bacterial root (60, 61). Quinone occurrences were derived from instrumental analysis of isolates or inferred from the presence of key biosynthesis genes (SI results; Supplementary Datafile S3; including literature data). Phenotype oxytolerance was curated from strain descriptions. Selected classes/orders denoted inside of rings. Selected phyla denoted outside of rings: ACD, *Aquificota*-*Campylobacterota*-*Deferribacterota*; Desulfob., *Desulfobacterota*; DST, *Deinococcota*-*Synergistota*-*Thermotogota*; BA, Bacillota-*Actinomycetota*; FCB, *Fibrobacterota*-*Chloroflexota*-*Bacteroidota*; Marg., *Candidatus* Margulisbacteria; Myxoc., *Myxococcota*; Nitrospin., *Nitrospinota*; PVC, *Planctomycetota*-*Verrucomicrobiota*-*Chlamydiota*; Seri., *Candidatus* Sericytochromatia; Vamp., *Vampirovibrionophyceae*. Circles indicate ultra-fast bootstrap support ≥95%.

### Ancient origin of high-potential quinones

The biochemical similarity of the HPQ biosynthesis pathways suggests a common ancestry. The mPQ biosynthesis proteins of *Nitrospirota* are most closely related to homologs from the UQ and PQ pathways of *Cyanobacteriota* and *Pseudomonadota*, which is unexpected given the phylogenetic distance between these phyla (Fig. 3). Specifically, the phylogenies of the key prenyltransferases and decarboxylases exhibit a consistent tree topology, with HPQ proteins being monophyletic relative to homologous proteins of the LPQ biosynthesis pathways (Fig. 4; see SI for expanded discussion). HPQ homologs from *Pseudomonadota* and *Nitrospirota* branch as sister lineages with respect to *Cyanobacteriota*. Other proteins of the HPQ pathways (chorismate-pyruvate lyase, decarboxylase cofactor) generally support this topology, although with lower branch support (Fig. S19-20). These patterns suggest a single, shared origin of the universal core of HPQ biosynthesis in bacteria.

**Fig. 4.**
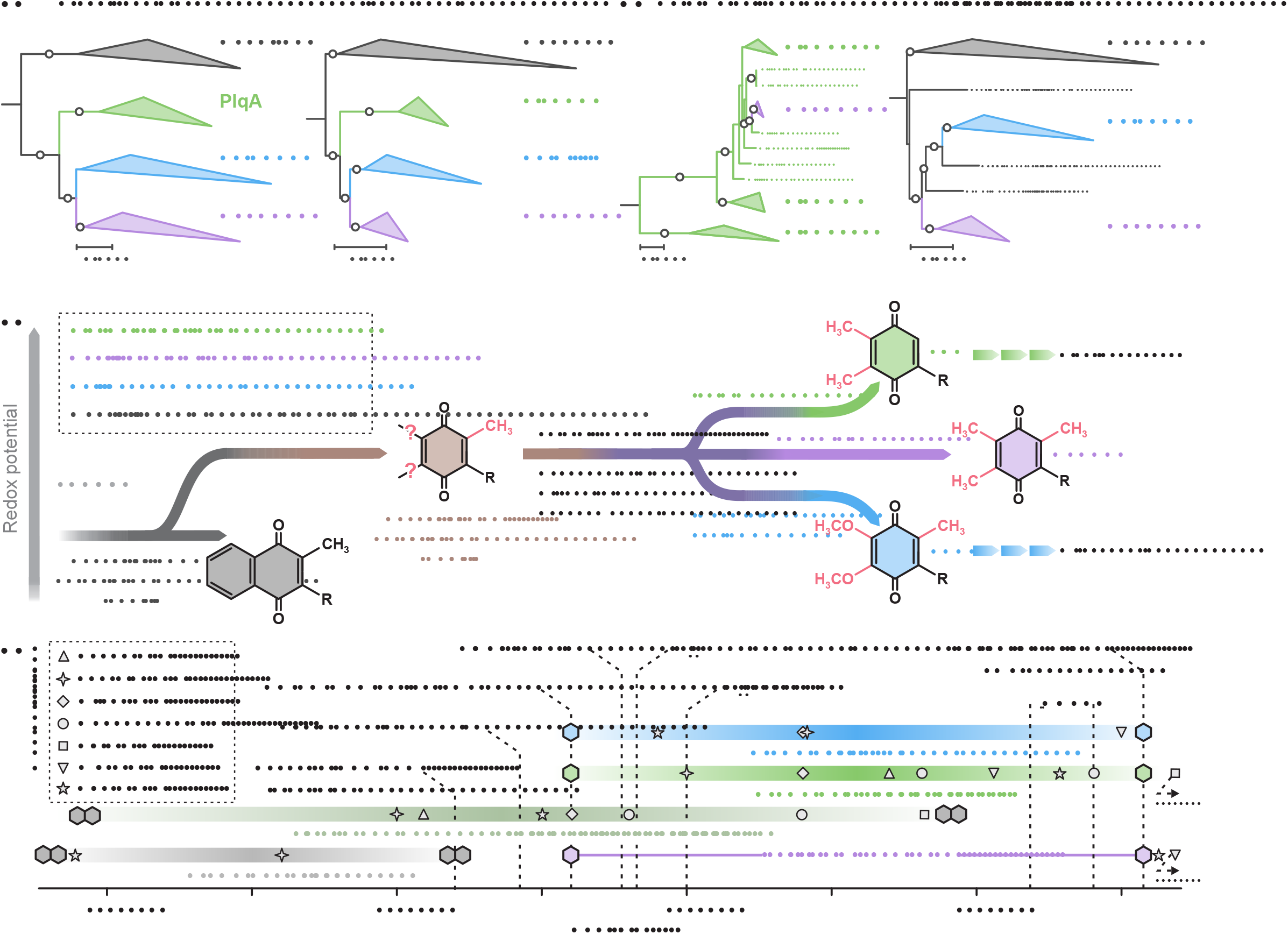
High-potential quinones (HPQ) share a single origin predating the great oxygenation event. **a**, Phylogenetic trees of HPQ biosynthesis proteins demonstrating that prenyltransferases and decarboxylases of the ubiquinone (UQ, UbiAD), plastoquinone (PQ, PlqAD), and methylplastoquinone (mPQ, MpqAD) pathways form sister clades of the archaeal and bacterial futalosine pathway for biosynthesis of menaquinone (MK, MqnPL). **b**, Phylogenetic trees of quinone C5/C6 (PlqQ, MpqQ) and C2 methyltransferases (UbiE, MpqE), showing a nested topology of C5/C6 methyltransferases and that C2 methyltransferases form a sister lineage of menaquinone-associated methyltransferases (MqnK). Outgroups used for rooting the trees are not shown but discussed in the Supplementary Information. Scale bars indicate 0.5 substitutions per site. Open circles indicate ultra-fast bootstrap support ≥95%. **c**, Conceptual sketch of HPQ evolution and resulting redox potentials, based on the trees in panels a-b. **d**, Timescale of LPQ and HPQ evolution (colors as in panel c; based on panels 4a-b and the observation that the last common ancestors of *Pseudomonadota, Cyanobacteriota*, and aerobic *Nitrospirota* contained UQ, PQ, and mPQ, respectively) in relation to geochemical changes (evidence for localized O_2_ oases (63–65), the great oxygenation event, GOE (7)) and biological innovations (Archean rapid genetic expansion (10), evolution of enzymes protecting against reactive oxygen species (ROS) (12), expansion of O_2_ reductase diversity (11)). Shaded hexagons indicate minimum and maximum estimates of HPQ evolution timescale. Open symbols indicate median ages (colored bars: uncertainty range; quinone symbols: upper/lower estimate) of relevant clades estimated by previous molecular clock analyses (Boden et al. (12); Davín et al. (66); Fournier et al. (67); Oliver et al. (68); Magnabosco et al. (69); Shih et al. (70); Ward et al. (71)). The earliest date of UQ/PQ/mPQ emergence is set as the earliest estimate of the radiation of crown Cyanobacteria, Pseudomonadota, and (aerobic) Nitrospirota assuming that UQ, PQ, or mPQ were present in the last common ancestor of each clade.

The distribution of LPQs, HPQs and their associated biosynthetic genes in *Bacteria* suggests that HPQ biosynthesis is conserved in all known lineages of *Cyanobacteriota, Pseudomonadota*, and aerobic *Nitrospirota* (Fig. 3, S11-13; see SI for expanded discussion). Conversely, HPQs are not found in anaerobic *Nitrospirota*, nor in the sister phyla of *Cyanobacteriota, Nitrospirota*, or *Pseudomonadota*, all of which produce LPQs via the MK_mqn_ pathway (Fig. 3; Table S5). Given that *Cyanobacteriota, Nitrospirota*, and *Pseudomonadota* are paraphyletic, vertical inheritance of HPQ pathways from a common ancestor is unlikely. Instead, HPQ occurrence and protein phylogenies indicate that an ancestral HPQ pathway was laterally acquired by stem-group *Cyanobacteriota, Pseudomonadota*, and aerobic *Nitrospirota* from an unknown or extinct donor lineage.

The ancestral HPQ pathway later diversified through changes to the C2, C5, and C6 substituents. Specifically, C2 methyltransferases are present in all LPQ and HPQ biosynthesis pathways except PQ. LPQ and HPQ C2 methyltransferases form sister clades (Fig. 4b) and C2 methyltransferases may thus be as old as the divergence between LPQ and HPQ pathways. Consequently, it is likely that the ancestral HPQ pathway contained a C2 methyltransferase that was lost prior to the radiation of crown group *Cyanobacteriota* (Fig. 4c). Lack of C2 methylation increases the redox potential of PQ (Table S3) and is essential for the functioning of the oxygen-evolving photosystem II (62). Loss of C2-methylation was likely linked to the evolution of oxygenic photosynthesis and therefore did not occur in *Pseudomonadota* and *Nitrospirota*. The evolution of the C5/C6 functional groups is less constrained. The C5/C6 methyltransferases of the PQ/mPQ pathways are poorly conserved in *Nitrospirota* and *Cyanobacteriota*, but at least one subgroup of *Nitrospirota* laterally acquired a C5/C6 methyltransferase from *Cyanobacteriota* (Fig. 4b; see SI discussion). C5/C6 methylation requires a single enzyme, whereas methoxylation to yield UQ requires at least two enzymes that are specific to *Pseudomonadota* (37, 55). Thus, the most parsimonious explanation is that the ancestral HPQ was methylated at C5/C6 in addition to C2, i.e., identical to mPQ, and that methoxylation evolved later (Fig. 4c).

It has been proposed that LPQs were present in the last universal common ancestor or evolved shortly thereafter, given their nearly universal presence in *Archaea* and *Bacteria* (72). Of the two LPQ biosynthetic pathways, the MK_mqn_ pathway is considered ancestral to basal *Archaea* and *Bacteria*, whereas the MK_men_ pathway was laterally transferred from *Bacteria* to a subset of *Archaea* (72). Homologous proteins suggest that the LPQ and HPQ biosynthetic pathways are evolutionarily related. The HPQ pathways share five homologs with the MK_mqn_ pathway (prenyltransferase, two-component decarboxylase, C2 methyltransferase, kinase) and two with the alternative MK_men_ pathway (prenyltransferase, C2 methyltransferase) (37). Our analysis shows that the MK_mqn_ homologs from *Archaea* and *Bacteria* form sister groups to the HPQ proteins, whereas the two homologs of the MK_men_ pathway are more distantly related to both HPQ and MK_mqn_ proteins (Fig. 4, S14-16, and SI discussion). This topology suggests that contrary to previous conclusions (73), the HPQ pathways did not descend directly from extant MK pathways.

Instead, the HPQ and MK pathways likely evolved from an ancestral quinone biosynthesis pathway that, like all extant pathways, used a chorismate derivative as precursor. In the case of HPQ, this precursor is prenylated in the second step, whereas prenylation is a late step in MK biosynthesis. The specificity of prenyltransferase for its quinone substrate (74) combined with early prenylation in the HPQ pathways may have facilitated evolutionary divergence of the HPQ and MK pathways. Existing machinery from the ancestral quinone biosynthesis pathway such as decarboxylase, C2 methyltransferase, and kinase were then co-opted by these new pathways. The deep phylogenetic divergence between HPQ and LPQ proteins (Fig. S14-16) suggests that the ancestral HPQ pathway could have emerged before the radiation of *Bacteria* and *Archaea* (72) 4.1-3.4 Ga ago in an extinct lineage coeval to the evolution of the extant LPQ pathways (Fig. 4c-d). Such an early origin of HPQs is not necessarily linked to aerobic respiration or oxygenic photosynthesis using high-potential ETCs. Instead, ancestral HPQs could have been involved in different functions, such as oxygen detoxification or a primordial form of high-potential photosynthesis (75, 76), and only later adopted into high-potential ETCs used for oxygenic photosynthesis and respiration using oxygen or other high-potential electron acceptors, such as nitric oxide (17, 77).

### Early evolution of aerobic metabolism

The association of HPQ biosynthesis with oxygenic photosynthesis and aerobic respiration in extant bacteria suggests that these traits became inseparably linked during evolution. The phylogeny of HPQ biosynthesis proteins therefore allows dating the origin of aerobic metabolisms using HPQs relative to Earth’s oxygenation. Oxygen first accumulated permanently in the atmosphere during the GOE (7) but geochemical tracers suggest oxygen was locally present during the late Archean (63, 64, 78). The likely presence of oxygen during the Archean aligns with the diversification of electron transport pathways, oxygenases, oxidoreductases, and antioxidant enzymes around 3.3-2.9 Ga (10–13, 79), i.e., long before the GOE. Alternative proposals place the emergence of crown group *Cyanobacteriota*, oxygenic photosynthesis, and aerobic respiration coeval to, or after, the GOE (4, 70, 80). Regardless of whether oxygenic photosynthesis emerged during the Archean or was coeval to the GOE, the phylogenetic split between HPQ and LPQ proteins and the presence of the MK_mqn_ pathway in the non-photosynthetic sister lineages (*Vampirovibrionophyceae*, “*Candidatus* Margulisbacteria”, “*Candidatus* Sericytochromatia”; Fig. 3, 4; Table S5; SI discussion) together indicate that the HPQ pathway in *Cyanobacteriota* originated from lateral transfer after their divergence from these sister lineages. Because PQ is central to the functioning of photosystem II in all extant oxygenic photosynthesizers (24, 81), emergence of PQ biosynthesis was likely tied to the evolution of oxygenic photosynthesis and thus may have existed before the radiation of crown group *Cyanobacteriota*. This supports earlier proposals that the extant oxygenic photosynthetic machinery originated in a lineage that diverged from the non-photosynthetic sister lineages (80, 82) but pre-dated the radiation of crown group *Cyanobacteriota* (76). Collectively, these constraints indicate that HPQs are at least as old as oxygenic photosynthesis by *Cyanobacteriota* and therefore predate the GOE.

Aerobic metabolism preceding the GOE is supported by the near-universal occurrence of aerobic respiration in crown group *Cyanobacteriota* and *Pseudomonadota*. All basal clades of *Cyanobacteriota* and *Pseudomonadota* possess HPQs and are capable of aerobic respiration, with only few late-branching *Pseudomonadota* being obligate anaerobes (Fig. 3, S10, S12-13). Molecular clocks calibrated using cyanobacteriotal fossils place the last common ancestor of crown group *Cyanobacteriota* and the emergence of basal, aerobic *Pseudomonadota* (*Magnetococcia*) around 2.5-3.2 Ga (10, 12, 66–68, 83), whereas aerobic *Nitrospirota* may have emerged shortly after the GOE (66, 71). HPQs were thus likely used for aerobic respiration by the time of the radiation of extant *Cyanobacteriota* and *Pseudomonadota*. Given the constraint that HPQs must have been present in stem-group *Cyanobacteriota* and *Pseudomonadota*, the minimum age of extant HPQs is between 2.5-3.2 Ga, whereas the ancestral HPQ pathway may be as old as crown-group *Bacteria* (3.4-4.1 Ga; Fig. 4d). We therefore suggest that aerobic respiration with high potential ETCs may have originated up to 800 Ma before oxygen permanently accumulated in the atmosphere during the GOE. Microbial mats could have provided a niche for chemoautotrophs and heterotrophs consuming oxygen provided by *Cyanobacteriota* directly (32), preventing escape to the atmosphere.

In modern ecosystems, some aerobic bacteria continue to use LPQ-rather than HPQ-dependent ETCs for aerobic respiration (23). In the presence of O_2_, reduced HPQs are relatively stable but reduced LPQs are rapidly autoxidized, resulting in the loss of reducing equivalents to O_2_ (26, 84). Further, aerobic respiration with LPQs leads to increased formation of deleterious reactive oxygen species (28, 85), requiring energy to be expended on the mitigation of cellular damage, thereby decreasing growth rates (28, 86). Finally, the use of LPQs instead of HPQs for proton pumping by complex I is less efficient (87, 88). Given these drawbacks, one might question why all aerobes have not switched from LPQs to HPQs to use oxygen without these disadvantages. The evolution of HPQs was a complex process closely tied to the evolution of the ETC itself, demanding not only the acquisition of a dedicated pathway for quinone biosynthesis, but also an upshift in redox potential of all other ETC components, including hemes and iron-sulfur clusters (27). These complex requirements may explain why the evolution of HPQ was successful only once and lateral transfers were rare. Distinct redox-tuning then led to the extant diversity of HPQ structures and biosynthetic pathways in *Cyanobacteriota, Pseudomonadota*, and *Nitrospirota*. Two of the three HPQ pathways were later obtained by eukaryotes, UQ via incorporation of an alphaproteobacterium as the mitochondrion and PQ via incorporation of a cyanobacterium as the chloroplast (58, 89) (Fig. 4d), while mPQ remained exclusively bacterial. Through their high potential ETCs, these lineages were able to rise to dominance in modern oxic ecosystems (32–36), as evidenced by the prevalence of HPQs in modern oxic environments (31, 90).

## Material & Methods

Detailed methods can be found in the SI Appendix. Cultures were grown in standard media with or without ^13^C-isotope-labeled compounds and harvested as described in the SI Appendix. Quinones were isolated using solvent extraction and chromatography and structurally characterized using high-performance chromatography coupled to high resolution tandem mass spectrometry or using nuclear magnetic resonance spectroscopy. Candidate genes and their phylogenies were identified using standard genomic and phylogenetic techniques and verified using heterologous complementation in *E. coli* mutants.

## Acknowledgements

We thank Susan Carter for laboratory assistance and Colleen Hansel, Gregory Fournier, and C.S. Raman for discussions. Linda L. Jahnke and Mary N. Parenteau are thanked for providing biomass of *G. violaceus*. Fay-Wey Li is thanked for providing the *A. panamensis* culture and Anja Engel for access to UPLC-Orbitrap-MS/MS. Two anonymous reviewers are thanked for comments that helped improve the quality of the manuscript. This work was funded through National Science Foundation grants 1702262 and 1843285 (to A.P.), the Gordon and Betty Moore Foundation (to A.P.), and the Deutsche Forschungsgemeinschaft grant 441217575 (to F.J.E). This work was also supported by the French National Research Agency through the grant ANR-21-CE02-0018 (to S.S.A.), and the “Investissements d’Avenir” program (ANR-15-IDEX-02) through the “Origin of Life” project and the IDEX-IRS 2020 call of the Grenoble-Alpes University (to S.S.A.). Research at MARUM was supported by the Deutsche Forschungsgemeinschaft via Germany’s Excellence Strategy, no. EXC-2077-390741603. T.W.E. acknowledges funding by the Alexander-von-Humboldt-Stiftung through the Feodor-Lynen-Fellowship. S.L. acknowledges funding by the Netherlands Organisation for Scientific Research (016.Vidi.189.050). Research at MIT was otherwise supported by the NASA Astrobiology Institute Exobiology Program grant number 18-EXXO18-0039. This result is part of a project that has received funding from the European Research Council (ERC) under the European Union’s Horizon 2020 research and innovation program (Grant agreement No. 946150, to J.D.H.). Research at Caltech was supported by the NASA Astrobiology Institute Exobiology grant no. 80NSSC19K0480. This research was supported through high-performance computing resources available at the Kiel University Computing Centre. Redox potential calculations were performed on the Euler cluster operated by the High-Performance Computing group at ETH Zurich. The authors thank Coline Fernandes from University Grenoble Alpes for running the ^1^H and ^13^C NMR experiments.

## Material & Data Availability

All data including alignments and phylogenetic trees are available through the Open Science Framework under doi:10.17605/OSF.IO/KNRW9. Data tables are additionally available in the supplementary materials. Strains are available from the authors upon request.

## Code Availability

All code for the calculation of redox potentials is available in the supplementary materials.

