## Supplementary information for "A novel quinone biosynthetic pathway illuminates the evolution of aerobic metabolism"

### Materials and Methods

#### ***Cultivation***

All cultures used in this study are listed in Table S4. *Nitrospira defluvii* A17, *N. lenta* BS10, and *N. moscoviensis* M-1 were grown chemolithoautotrophically in 10-L glass bottles containing 6 L freshwater medium with 7 mM sodium nitrite (1) and were harvested after nitrite was completely consumed. *N. marina* Nb-295 was grown chemolithoautotrophically in 10-L glass bottles containing 6 L artificial seawater medium (2) with 1 mM sodium nitrite. *N. inopinata* was grown chemolithoautotrophically in 0.5-L glass bottles containing 200 mL mineral ammonia oxidizer medium with 1 mM ammonium chloride(3). *Ca*. N. kreftii was maintained under chemolithoautotrophic conditions in a continuous membrane bioreactor in NOB mineral salts medium(4) receiving 2.5 mM ammonium chloride day$^{-1}$. *Leptospirillum ferrooxidans* DSM 2705 was grown chemolithoautotrophically in a 10-L glass bottle containing 6 L of DSMZ medium 882 and was harvested after 7 days. *Nitrospira* spp. and *L. ferrooxidans* cultures were inoculated with 1% of mid-growth phase pre-cultures and incubated without stirring at 28 °C in the dark. A co-culture of *Candidatus* Manganitrophus noduliformans and *Ramlibacter lithotrophicus* RBP-1 was grown aerobically and chemolithoautotrophically in a 2-L Erlenmeyer flask containing 1 L of defined minerals and vitamins basal medium buffered with MOPS (5 mM, pH 7.2), as previously described(5). The medium contained 40 g manganese(II) carbonate hydrate (MilliporeSigma) as sole electron donor and 1 mM sodium nitrate as sole source of fixed nitrogen. After prolonged incubation at 35 °C in the dark, the cells along with the particulate manganese(II/IV) oxides to which cells strongly attach, were harvested by centrifugation after completion of the oxidation of the manganese carbonate starting material. A clonal pure culture of *R. lithotrophicus* str. RBP-1 was grown heterotrophically and anaerobically under $N_2$ in a butyl rubber-stoppered, 1-L glass bottle containing 200 ml of the same defined minerals and vitamins basal medium buffered with MOPS (5 mM, pH 7.2) as used for the manganese oxidizing co-culture, above, but without manganese carbonate. The medium was amended with tryptone (5 g/L), L-aspartic acid sodium salt monohydrate (2 g/L), sodium formate (1 g/L), and sodium nitrate (10 mM). Cells were incubated at 35 °C in the dark and harvested by centrifugation as soon as the culture medium became turbid.

*Anthocerotibacter panamensis* was grown on BG-11 solid medium. Proteobacterial or eukaryotic contamination of the culture was suspected after detection of $UQ_{9:9}$ in *A. panamensis* quinone extracts. No contamination was detected during phase-contrast microscopy (1000x magnification). To further test for contamination, a single colony of *A. panamensis* was resuspended in BG-11 medium and streaked onto a range of heterotrophic media (potato-dextrose agar, LB agar, Saboraud agar, tryptic soy agar, or BG-11 with 0.01% yeast extract). Yellowish-white colonies of 2-3 µm long rod-shaped cells developed after four days of incubation on LB, tryptic soy, or BG-11-yeast extract agar at room temperature under ambient light. A single colony was picked from the BG-11-yeast extract plate, re-streaked onto the same substrate and used for Sanger sequencing of the 16S rRNA gene (GENEWIZ, Leipzig, Germany), revealing 99% sequence identity to *Pseudomonas lutea*. For quinone analysis, a colony from the same plate was grown in LB broth for 4 days and harvested by centrifugation.

For labeling experiments, the three aerobic *Nitrospirota* species *N. moscoviensis*, *N. lenta*, and *L. ferrooxidans*, were grown in triplicate in 0.5 L medium in 2-L bottles. Culture conditions were as described above with the exception that only 2 mM sodium nitrite was used for *N. moscoviensis* and 1 mM sodium nitrite for *N. lenta* and *N. marina*. Labeled tyrosine or 4-hydroxybenzoic acid (99% ring-$^{13}C_6$; Cambridge Isotope Laboratories, Andover, MA, USA) were added in early exponential phase at a final concentration of 10 µM (for *N. moscoviensis*, *N. lenta*, *L. ferrooxidans*). Additionally, *N. moscoviensis* was grown in the same way in the presence of 100 µM methyl-$^{2}H_3$-methionine (98% methyl-$^{2}H_3$; Cambridge Isotope Laboratories). Cultures were harvested at the onset of the stationary phase as described above. Labeling experiments were not performed with *M. noduliformans* due to the lack of a pure culture and the slow growth of the co-culture.

Biomass of *Nitrospira* spp. and *L. ferrooxidans* was harvested by filtration through two stacked, combusted (450 °C, 6 h) glass fiber filters (GF-75, 0.3 µm pore size, Advantec MFS, Dublin, CA, USA). Biomass of *T. islandicus*, the *Ca*. Manganitrophus noduliformans-*Ramlibacter lithotrophicus* co-culture, and the *R. lithotrophicus* pure culture were harvested using centrifugation. Biomass of *A.*

*panamensis* was grown in BG-11 medium at ambient light and room temperature and harvested by centrifugation. The *Anthocerotibacter panamensis* culture was a gift of Fay-Wey Li (Cornell University, Ithaca, NY, USA). Biomass of *Gloeobacter violaceous* was a gift from Linda L. Jahnke and Mary N. Parenteau (NASA Ames Research Center, Moffett Field, CA, USA).

***Identification of candidate genes***

Query sequences for proteins of the known quinone biosynthetic pathways (PQ, UQ, $MK_{mqn}$, $MK_{men}$) were selected from the UniProt (http://www.uniprot.org/) and InterPro 7 databases(6) based on two criteria, i) production of the respective quinone confirmed in culture and ii) confirmed protein function. These protein sequences were used as queries to search *Nitrospirota* genomes in the NCBI nr database and the JGI IMG database for homologs using blastp v2.7.1 (7). An expectation value of $10^{-5}$, coverage of >50%, and amino acid similarity of >20% were used as cutoffs. Candidate genes were identified through amino acid similarity, conserved functional domains, and by clustering with query sequences in phylogenetic trees. UbiC homologs could not be identified in *Nitrospirota* using blastp with query sequences from *Pseudomonadota* and *Cyanobacteriota*. UbiC homologs in *Nitrospirota* were identified through the conserved functional Pfam domain PF04345 (chorismate lyase) in Interpro. These homologs were then used as query sequences in blastp to identify further homologs in *Nitrospirota*.

To complement candidate proteins identified through blastp, Hidden Markov models (HMMs) were used for annotation of quinone biosynthesis proteins in *Nitrospirota* genomes. All 1117 genomes of the phylum *Nitrospirota* available at the NCBI database were downloaded (last accessed in October 2022) regardless of their level of assembly and completeness. Their quality was assessed using CheckM2 (version 0.1.3) (8). The genomes were considered either as high (Completion > 90% and Contamination < 5%), medium (Completion >= 50% and Contamination < 10 %) or low quality(9). Other genomes were added from the IMG database: *Candidatus* Manganitrophus morganii SA1, *Candidatus* Manganitrophus morganii SB1, Nitrospirae bacterium SCGC AC-732-L14, Nitrospirae sp. genome_bin_8, Nitrospirae bacterium SCGC AB-219-C22, and *Nitrospira marina* 295. Genomes without CDS annotation from the NCBI annotation pipeline were annotated with Bakta (v 1.8.1, database v5.0 full) (10). We used HMM profiles for ubiquinone, menaquinone, and plastoquinone pathways previously designed and presented in Pelosi et al. (11), Kazemzadeh et al. (12), and Chobert et al. (in preparation), amounting to a total of 93 HMM profiles used including 48 profiles for the genes of interest and 45 “decoy” profiles designed to increase the annotation specificity of some genes. These profiles were used to perform a similarity search on the set of *Nitrospirota* genomes with the hmmscan program (HMMER suite v3.3.2) (13). We selected the hits with an i-Evalue < 0.05 and a profile coverage value > 0.5. The profile coverage corresponds to the proportion of the profile length that aligns with the protein sequence. When several profiles matched a sequence, only the hit with the best i-Evalue was considered (Supplementary Datafile S1). Results from HMM profile searches identified the same set of quinone proteins found using blastp.

***Phylogenetic analysis***

For analysis of quinone occurrence, 284 representative genomes covering all major lineages (classes, orders) of Proteobacteria (*Pseudomonadota*; 130 genomes), *Cyanobacteriota* (74 genomes), and all *Nitrospirota* (84 genomes) were retrieved from the NCBI and JGI IMG databases. Genomes of *Cyanobacteriota* and *Pseudomonadota* were randomly selected. Genomes where occurrence of PQ and UQ had previously been confirmed experimentally were added manually and included in the analyses. Incomplete genomes and MAGs of *Cyanobacteriota* and *Nitrospirota* were included due to the small number of isolates for *Nitrospirota* and basal *Cyanobacteriota* (Gloeobacterales). Symbionts and fast-evolving lineages (Rickettsiales, Pelagibacterales, Holosporales, Legionellales, and Francisellales) were excluded. Seven genomes were used as outgroup for the species trees (*Chloroflexus aggregans* DSM 9485, *Pirellula staleyi* DSM 6068, *Desulfovibrio vulgaris* DP4, *Ramlibacter tataouinensis* TTB310, *Nitrospina gracilis* 3/211, *Acidobacterium capsulatum*, *Candidatus* Methylomirabilis oxyfera; *R. tataouinensis* omitted from outgroup in *Pseudomonadota* tree; analogous to ref. (14)). Genome completeness and contamination was determined using CheckM v1.2.2 (15). Only genomes that met the minimum criteria (9) for high (completeness >90%, contamination <5%) and medium quality genomes (completeness >50%, contamination <5%) were retained for downstream analyses. For species trees (Fig. 2, S11-13), protein sequences of 120 single copy markers were identified, aligned, filtered, and concatenated using GTDB-Tk v2.2.5 with reference data version r207 (16, 17). In brief, markers were identified and annotated using Prodigal V2.6.3 (18) and HMMER 3.3.2 (13), aligned using hmmalign 3.3.2 and filtered to ~5000 amino acids (i.e., 42 positions retained per marker for the final alignment) using the default for bacteria in GTDB-Tk. The alignments were trimmed using trimAl 1.2 using the automated trimming heuristic (19). The resulting alignments (~4500 positions) were used to construct maximum-likelihood trees using IQ-TREE v2.2.0.3(20) with Modelfinder(21) and 10,000 ultrafast bootstrap approximations (22). The best-fitting substitution models were LG+F+R8 (*Cyanobacteriota*), Q.yeast+F+I+R6 (*Nitrospirota*), and LG+F+R9 (*Pseudomonadota*). Phylogenetic trees were visualized using TreeViewer 2.1.0(23). Experimentally confirmed occurrences of quinone types of *Cyanobacteriota* (7/74) and *Pseudomonadota* (85/130) were manually curated from the literature in addition to those identified in this study (*Cyanobacteriota*: 2; *Nitrospirota*: 9; Supplementary Datafile S3). For isolates outside these phyla, quinone types were manually curated from published strain descriptions. For all other species/MAGs, quinone types were inferred from the genomes using the presence of at least one characteristic marker protein identified here and in other studies(24, 25) (UQ pathway: UbiA/UbiD/UbiE; PQ pathway: PlqA/PlqD; mPQ pathway: MpqA/MpqD/MpqE; MK pathways: MenA/MenB/MenF/MenG or MqnA/MqnC/MqnD/MqnL/MqnP). Quinone types could be inferred for 430 of 568 studied genomes using blastp searches (Supplementary Datafile S3). Oxytolerance (aerobe/anaerobe/facultative anaerobe) was manually curated from published strain descriptions of isolates (275 of 568 studied genomes).

For quinone biosynthetic gene phylogenies, protein query sequences from the PQ/mPQ/UQ pathways of *Cyanobacteriota*, *Nitrospirota*, and *Pseudomonadota* as well as the respective homologs of the $MK_{men}$ and $MK_{mqn}$ pathways were used as query sequences for searches using blastp (Supplementary Datafile S3). The search space included selected *Cyanobacteriota*, *Nitrospirota*, and *Pseudomonadota* genomes, as described above, as well as one randomly picked representative genome of each bacterial (182 genomes) and archaeal (20 genomes) phylum. To further enhance phylogenetic resolution, genomes of phyla closely related to *Nitrospirota* ("*Candidatus* Methylomirabilota", *Nitrospinota*), *Pseudomonadota* (*Campylobacterota*, *Myxococcota*, *Bdellovibrionota*,

*Thermodesulfobacteriota, Desulfobacterota*), and *Cyanobacteriota* ("*Candidatus* Margulisbacteria", "*Candidatus* Sericytochromatia", *Vampirovibrionophyceae*) were added by picking one random representative genome for each order. All random picks were drawn from the list of representative genomes in the Genome Taxonomy Database (version r207) using GToTree v1.8.3(26). The resulting 569 genomes (Supplementary Datafile S3) were used to construct a blast database. An expectation value of $10^{-5}$, coverage of >50%, and amino acid similarity of >20% were used as cutoffs. The resulting hits were added to the dataset derived from HMM searches (described above) and the whole dataset was dereplicated to retain only unique sequences. Suitable outgroups with distinct biochemical functions were identified through Interpro and Pfam protein families and validated through unrooted phylogenetic trees and protein family-level phylogenies. The outgroups were UbiA/MpqA/PlqA/MqnP: COX10; UbiB/MpqB/PlqB: Rio1; UbiC/MpqC/PlqC: TreR; UbiD/MpqD/PlqD: Fdc1+SmdK+TtnD+PpcB+PhdA; UbiE/MpqE: PmtA; MpqQ/PlqQ: PmtA; UbiX/MpqX/PlqX: Pad1+SmdJ+TtnC+PhdB. Sequences were aligned using MAFFT v7.388 (L-INS-I, BLOSUM62, gap open penalty: 1.53)(27). Alignments were trimmed using clipkit v2.1.1, using the smart-gap trimming mode(28), to reduce computation time. Phylogenetic trees were generated using IQ-TREE v2.2.0.3(20) with Modelfinder(21) and 1,000 ultrafast bootstrap approximations(22). The best-fitting models were LG+I+G4 for UbiX/MpqX, LG+F+I+G4 for UbiA/MpqA/PlqA and ubiC/mpqC/plqC, LG+F+G4 for UbiD/MpqD and UbiB/MpqB/PlqB, and Q.pfam+G4 for UbiE/MpqE and MpqQ/PlqQ.

Sequence motifs of Rieske proteins were identified by performing a tBastn or a blastp against the Rieske protein sequence from *Heliobacterium modesticaldum* on a representative selection of *Nitrospirota* genomes (Table S4). The retrieved sequences were analyzed to confirm the presence of the cluster binding motifs (CxxLGC. . . CPCHxXxY) and to note the two residues X and Y that determine the cluster redox midpoint potential. Only sequences for which the genes downstream of the Rieske protein sequence could be identified as cytochrome *b* or cytochrome $b_6$ and subunit IV were retained.

***Gene synthesis***

The genes tested experimentally in this study are listed in Supplementary Datafile S2. The genes were synthesized by the “Genecust” company and were cloned into the pBAD24i or pBAD33i vectors(12) downstream of an arabinose-inducible promoter using NcoI (5’ end) and HindIII (3’ end) restriction enzymes. The nucleotide sequences were optimized for expression in *E. coli* and are available in Supplementary Datafile S2. The ATG start codon of the genes is comprised in the NcoI site sequence (CCATGG), which imposes a G as the first nucleotide of the second codon. If the second amino acid did not correspond to a codon starting with a G, an alanine codon (GCG) was added before the second codon of the nucleotide sequence.

***Construction of E. coli deletion strains.***

The antibiotic resistance cassette of strain Δ*ubiIF* was eliminated using plasmid pCP20 as described previously(29), resulting in strain Δ*ubiIFc*. The *ubiE*::kan mutation was introduced into Δ*ubiIFc* by P1 vir transduction from the Δ*ubiE* donor strain, selecting for kan resistance(30). Clones of the resulting Δ*ubiIFE* strain were verified by PCR amplification of the *ubiF* and *ubiE* loci using primers UbiF5/UbiF3 and UbiE5/UbiE3, respectively (Table S7).

***In vivo complementation assay in E. coli strains***

*E. coli* strains were transformed by heat shock with plasmids containing *Nitrospirota* genes and selected on LB-agar plates containing the appropriate antibiotics. Individual clones were grown microaerobically overnight at 37 °C in 1.5 mL microcentrifuge tubes filled to the top with LB medium plus antibiotics. Then, 50 μL of the preculture was used to inoculate 5 mL LB with antibiotics and 0.05% arabinose to induce the expression of the genes cloned in the pBAD33i* or pBAD24i* plasmids. The glass tubes were incubated overnight at 37 °C with 180 rpm shaking and the cells were collected by centrifugation at 4000 × *g*, 10 min, 4°C. The pellet was rinsed in cold PBS buffer and cells were transferred to pre-weighed 1.5 mL microcentrifuge tubes. After centrifugation at 12,000 × *g* at 4 °C for 1 min and elimination of supernatant, the cell wet weight was determined (10 to 20 mg), and pellets were stored at -20 °C.

***HPLC-ECD-MS analysis of quinones produced in E. coli***

Quinone analysis by HPLC-ECD-MS (ElectroChemical Detection-Mass Spectrometry) was carried out as described previously(11) with a mobile phase composed of 50% methanol, 40% ethanol and 10% of a mix of 90% isopropanol, 10% ammonium acetate (1 M), and 0.1% trifluoroacetic acid. The dried lipid extracts were suspended in 100 μL ethanol and a volume corresponding to 1 mg of cells was injected. In our chromatographic system, a precolumn electrode set at +650 mV ensured the oxidation of all reduced quinones present in the extracts prior to arrival on the HPLC column. MS detection was performed on an MSQ spectrometer (Thermo Scientific) with electrospray ionization in positive mode (probe temperature, 400°C; cone voltage, 80 V). MS spectra were recorded between m/z 600 and 950 with a scan time of 0.3 s and single-ion monitoring (SIM) was used to detect the following compounds: OPP ($M+NH_4^+$, m/z 656 to 657), $OQ_8$ ($M+NH_4^+$, m/z 670 to 671), $PQ_8$ ($M+NH_4^+$, m/z 698 to 699), $mPQ_8$ ($M+NH_4^+$, m/z 712 to 713), $UQ_8$ ($M+NH_4^+$, m/z 744 to 745).

***High-resolution mass spectrometric analysis of quinones***

For structural analysis, quinones were generally extracted using a modified four-step Bligh and Dyer extraction(31, 32). The extraction procedure was further modified for the *Ca.* Manganitrophus noduliformans-*Ramlibacter lithotrophicus* co-culture. To dissolve the insoluble manganese oxide and manganese carbonate encrusting the biomass, the co-culture cell pellet was treated with citric acid for 1 hour at room temperature before extraction (15 ml saturated citric acid added to ~15 ml culture slurry). The first two steps of Bligh and Dyer extraction were then performed using citric acid instead of phosphate buffer, followed by two steps using trichloroacetic acid buffer. The same protocol was applied to extract *R. lithotrophicus* biomass. The total lipid extracts (TLEs) were gently dried under $N_2$, reconstituted in methanol, and stored at -20 °C.

Quinones were analyzed using high-performance liquid chromatography–mass spectrometry (HPLC-MS) as described previously(33, 34). Briefly, aliquots of the TLEs dissolved in methanol were injected into a coupled HPLC-MS system consisting of an Agilent 1200 series HPLC and an Agilent 6520 quadrupole time-of-flight MS equipped with an electrospray ionization interface operated in positive mode (Agilent Technologies, Santa Clara, CA, USA. $MS^2$ precursor selection was performed in data-dependent mode targeting the two most abundant ions per $MS^1$ scan with an isolation width of

4 Da and active exclusion after 2 spectra over 0.4 min. Due to instrument unavailability, a subset of samples was analyzed on a Dionex Ultimate 3000 HPLC coupled to a Bruker maXis Ultra-High Resolution quadrupole time-of-flight tandem MS equipped with an electrospray ionization ion source operating in positive mode (Bruker Daltonik, Bremen, Germany) coupled to the UHPLC system. The Bruker maXis MS was set to a resolving power of 27000 at *m*/*z* 1222 and every analysis was mass calibrated by loop injections of a calibration standard and correction by lock mass, leading to a mass accuracy of < 1–3 ppm. Ion source and other MS parameters were optimized by infusion of standards into the eluent flow from the HPLC system using a T-piece. For both HPLC-MS instruments, quinones were chromatographically separated using an ACE3 $C_{18}$ column (2.1 × 150 mm; 3 µm particle size; Advanced Chromatography Technologies, Aberdeen, Scotland) maintained at 45 °C as described previously(35). Quinones were identified by retention time, $MS^2$ fragment spectra, accurate molecular mass, and isotope pattern match of proposed sum formulas in full scan mode using Agilent MassHunter B.06.00 and Bruker DataAnalysis 4.4 software. A TLE extracted from spinach containing $UQ_{10:10}$, $MK_{4:1}$ (phylloquinone), and $PQ_{9:9,}$ as well as a commercial $MK_{4:4}$ standard (Sigma Aldrich) were used as quality controls for HPLC-MS and $MS^2$ analyses.

***Heterologous production of mPQ in E. coli and purification of mPQ***

The *E. coli* Δ*ubiJ* strain transformed with the plasmid pTU2a-[J23108-*mpqQ*]-[J23114-*plqH*] (Beltran et al., in preparation) was grown at 37 °C with shaking at 180 rpm in LB medium supplemented with chloramphenicol (Cm, 34 mg/L). Four 1-L cultures in LB+ Cm (in 5-L Erlenmeyer flasks) were each inoculated with 75 mL of overnight preculture and grown at 37 °C for 8 hours. Cultures were cooled at 4 °C for 2 hours and cells were harvested by centrifugation at 7,020 × g for 8 min at 4 °C. Cell pellets were suspended in 60 mL PBS and transferred in 50 mL centrifuge tubes for further centrifugation at 3,200 × *g*, 4 °C, 10 min. The supernatant was removed and the cell pellets (total weight 29 g) were frozen in liquid nitrogen and stored at -20 °C.

The pellets were thawed at room temperature and homogenized by vortexing in an aqueous solution of 1 mM ferricyanide and 0.15 M KCl (2 mL per gram of cells). The cell suspension was transferred to a 500 mL round-bottom flask together with methanol (10 mL per gram of cells) and placed in a sonication bath at room temperature for 10 min. Quinones were extracted with petroleum ether (40-60 °C boiling range, 7 mL per gram of cells) and sonication was repeated for 3 min. Phase separation was achieved by centrifugation at 7,020 × *g*, 4 °C, 8 min.

The top phase (petroleum ether) was transferred in another 500 mL round-bottom flask, and the extraction was repeated by adding petroleum ether to the remaining methanol phase. The second petroleum ether extract was combined with the first, and the solution was evaporated under reduced pressure at 50 °C, leaving an oily residue in the flask. This lipid extract was suspended in 400 mL hexane:water (1:1), mixed by several inversions, and sonicated for 3 min. After phase separation (obtained without centrifugation), the hexane phase was transferred to a pre-weighed 250 mL round-bottom flask, protected from light by an aluminum foil, and evaporated in the Rotavap. The hexane extraction was repeated a second time, and the solvent was evaporated in the same 250 mL flask, which was then sealed and stored overnight at 4 °C. A silica chromatography column was prepared using ~ 60 g silica (Macherey-Nagel, pore size 60 Å) suspended in hexane, with a thin layer of sand on top. The column was equilibrated with 1 volume of hexane and the quinone extract (suspended in ~4 mL of hexane) was deposited. Elution was performed with 3 column volumes of hexane:diethyl ether (97.5:2.5), followed by 3 column volumes of hexane:diethyl ether (90:10). The fractions were collected in test tubes protected from light with aluminum foil and were analyzed by TLC plates (Macherey-Nagel, Layer 0.2 mm, silica gel with fluorescent indicator $UV_{254}$). The quinones were visualized under UV light. $mPQ_8$ migrated with a $R_f$ of 0.67 (in hexane:diethyl ether, 90:10). The different quinones eluted from the column in the following order: $mPQ_8$, $MK_8$, $PQ_8$, $DMK_8$. The fractions containing $mPQ_8$ were further analyzed by HPLC-ECD-MS and the fractions with the highest purity of $mPQ_8$ were pooled and evaporated under vacuum. A total of 2.3 mg of $mPQ_8$ was obtained.

***Structural characterization of heterologously produced mPQ***

$mPQ_8$ produced heterologously in *E. coli* was further characterized by high-resolution mass spectrometry and nuclear magnetic resonance spectroscopy (NMR). Mass spectrometric analysis was performed using UHPLC-MS on a Vanquish Horizon UHPLC system equipped with a Waters Acquity BEH $C_{18}$ column (150 × 2.1 mm, 1.7 µm particle size) coupled to a Q Exactive Plus Orbitrap high-resolution MS (Thermo Fisher Scientific, Bremen, Germany) equipped with a heated electrospray ionization source (HESI) operating in positive ionization mode. Reversed-phase chromatographic conditions were as described previously(36). In brief, the injection volume was 10 µl, and mPQ was eluted at a constant flow rate of 0.4 ml $min^{-1}$ using linear gradients of methanol-water (85:15, vol/vol) (eluent A) to methanol-isopropanol (50:50, vol/vol; eluent B), both with 0.04% formic acid and 0.1% $NH_3$. The initial condition was 100% A held for 2 min, followed by a gradient to 15% B in 0.1 min and a gradient to 85% B in 18 min. The column was then washed with 100% B for 8 min. The column temperature was 65 °C. HESI and MS settings were optimized during infusion of a mixture of lipids including a $UQ_{10:10}$ standard (Sigma Aldrich, St Louis, MO, USA). HESI settings were: capillary temperature, 250 °C; sheath gas ($N_2$) pressure, 30 arbitrary units (AU); auxiliary gas ($N_2$) pressure, 10 AU; spray voltage, 3.5 kV; probe heater temperature, 350 °C; S-lens 75 V. mPQ was detected scanning from *m/z* 150 to 2000, followed by data-dependent $MS^2$ (isolation window 1.2 *m/z*) using resolving powers of 140,000 and 17,500 (FWHM at *m/z* 200) in full-scan MS and $MS^2$, respectively. Optimal fragmentation of mPQ was achieved using a stepped normalized collision energy of 15, 25, and 50. A mass calibration was performed using the Thermo Scientific Pierce LTQ Velos ESI Positive Ion Calibration Solution prior to analysis.

NMR spectra were acquired at the NMR-ICMG platform of Grenoble on a Bruker Avance-400 (400 MHz for $^{1}H$) and a Bruker Avance-500 instrument for 13C (125 MHz for $^{13}C$) at room temperature. Chemical shifts (δ) are reported in parts per million (ppm) relative to $CDCl_3$ as the solvent used [$^{1}H$: $\delta(CDCl_3)$ = 7.27 ppm, $^{13}C$: $\delta(CDCl_3)$ = 76 ppm].

***Calculation of quinone redox potentials***

The p*K*a and redox potential, $E^0$, for a suite of quinone compounds, including MK, PQ, mPQ, and UQ with varying isoprenoid sidechain lengths were estimated using density functional theory calculations. These methods broadly follow those of Huynh et al.(37). The p*K*a and $E^0$ values for any quinone can be calculated relative to experimentally determined values for a reference quinone, typically taken to be 1,4-benzoquinone, using isodesmic reactions. For the reduction of quinone Q to semiquinone $Q^{-}$ and simultaneous oxidation of reference semiquinone $R^{-}$ to R, the net reaction can be written as the sum of two half-reactions:

$$(1)\quad \begin{array}{cccccccc|c} Q & + & e^{-} & \rightarrow & Q^{-} & & & & -F\mathrm{E}_{1}^{0} \\ R^{-} & & & \rightarrow & R & + & e^{-} & & F\mathrm{E}_{\mathrm{ref},1}^{0} \\ \hline Q & + & R^{-} & \rightarrow & Q^{-} & + & R & & \Delta\mathrm{G}_{r,1}^{0} \end{array}$$

where subscript "1" indicates these values are for reaction 1 and subscript "ref" designates the $\mathrm{E}^0$ value for the reference quinone. Here, we have utilized the fact that $\Delta G^0 = -nF\mathrm{E}^0$ for a reduction reaction relative to the normal hydrogen electrode (NHE), where $F$ is Faraday's constant and $n = 1$ here since a single $e^{-}$ transfer is involved. Subsequent reduction of semiquinone $Q^{-}$ to reduced quinone $Q^{2-}$ can similarly be written as:

$$(2)\quad \begin{array}{cccccccc|c} Q^{-} & + & e^{-} & \rightarrow & Q^{2-} & & & & -F\mathrm{E}_{2}^{0} \\ R^{2-} & & & \rightarrow & R^{-} & + & e^{-} & & F\mathrm{E}_{\mathrm{ref},2}^{0} \\ \hline Q^{-} & + & R^{2-} & \rightarrow & Q^{2-} & + & R^{-} & & \Delta\mathrm{G}_{r,2}^{0} \end{array}$$

Thus, the $\mathrm{E}_i^0$ value for the quinone of interest can be calculated as the sum of the right-hand terms:

$$(3)\qquad \mathrm{E}_i^0 = \frac{\Delta\mathrm{G}_{r,i}^0}{F} + \mathrm{E}_{\mathrm{ref},i}^0$$

where $i = 1, 2$ is the reaction of interest, $\Delta\mathrm{G}_{r,i}^0$ is determined computationally, and $\mathrm{E}_{\mathrm{ref},i}^0$ is known experimentally. Similarly, for the deprotonation of quinol $H_2Q$ to $HQ^{-}$ and simultaneous protonation of reference quinol $HR^{-}$ to $H_2R$, the net reaction can be written as:

| | | | | | | | | |
|---|---|---|---|---|---|---|---|---|
| | $H_2Q$ | | | → | $HQ^-$ | + | $H^+$ | $RT\ln(10)\mathrm{p}K\mathrm{a}_3$ |
| (4) | $HR^-$ | + | $H^+$ | → | $H_2R$ | | | $-RT\ln(10)\mathrm{p}K\mathrm{a}_{\mathrm{ref},3}$ |
| | $H_2Q$ | + | $HR^-$ | → | $HQ^-$ | + | $H_2R$ | $\Delta G^0_{r,3}$ |

Here, we have utilized the fact that $\Delta G^0 = -RT\ln(10)\mathrm{p}K\mathrm{a}$ for a protonation reaction, where $R$ is the ideal gas constant and $T$ is temperature in Kelvin. Subsequent deprotonation of $HQ^-$ to $Q^{2-}$ can similarly be written as:

| | | | | | | | | |
|---|---|---|---|---|---|---|---|---|
| | $HQ^-$ | | | → | $Q^{2-}$ | + | $H^+$ | $RT\ln(10)\mathrm{p}K\mathrm{a}_4$ |
| (5) | $R^{2-}$ | + | $H^+$ | → | $HR^-$ | | | $-RT\ln(10)\mathrm{p}K\mathrm{a}_{\mathrm{ref},4}$ |
| | $HQ^-$ | + | $R^{2-}$ | → | $Q^{2-}$ | + | $HR^-$ | $\Delta G^0_{r,4}$ |

Similar to above, the $\mathrm{p}K\mathrm{a}_i$ value for the quinone of interest can thus be calculated as

(6) $$\mathrm{p}K\mathrm{a}_i = \frac{\Delta G^0_{r,i}}{RT\ln(10)} + \mathrm{p}K\mathrm{a}_{\mathrm{ref},i},$$

where $i = 3, 4$ is the reaction of interest, $\Delta G^0_{r,i}$ is determined computationally as above, and $\mathrm{p}K\mathrm{a}_{\mathrm{ref},i}$ is known experimentally. Finally, the redox potential for the complete reduction and protonation of quinone Q to quinol $H_2Q$ can be determined as:

| | | | | | | | | |
|---|---|---|---|---|---|---|---|---|
| | Q | + | $e^-$ | | | → | $Q^-$ | $-FE^0_1$ |
| (7) | $Q^-$ | + | $e^-$ | | | → | $Q^{2-}$ | $-FE^0_2$ |
| | $HQ^-$ | | | + | $H^+$ | → | $H_2Q$ | $-RT\ln(10)\mathrm{p}K\mathrm{a}_3$ |

| | | | | | | | |
|---|---|---|---|---|---|---|---|
| $Q^{2-}$ | | | + | $H^+$ | → | $HQ^-$ | $-RT\ \ln(10)\mathrm{p}K\mathrm{a}_4$ |
| Q | + | $2e^-$ | + | $2H^+$ | → | $H_2Q$ | $\Delta G^0_{r,\mathrm{net}}$ |

where absolute values for $E^0_1$, $E^0_2$, p$K$a$_3$, and p$K$a$_4$ are determined using Eqs. 3, 6 and experimentally determined values for the reference compound, 1,4-benzoquinone. Summing the right-hand equations in Eq. 7 yields:

(8) $$\Delta G^0_{r,\mathrm{net}} = -F(E^0_1 + E^0_2) - RT\ \ln(10)(\mathrm{p}K\mathrm{a}_3 + \mathrm{p}K\mathrm{a}_4).$$

Again, using the identity that $\Delta G^0 = -nFE^0$ relative to NHE for a reduction reaction, now with $n$ = 2 for the overall reaction, this can be rewritten as

(9) $$E^0_{\mathrm{net}} = \frac{1}{2}(E^0_1 + E^0_2) + \frac{RT\ \ln(10)}{2F}(\mathrm{p}K\mathrm{a}_3 + \mathrm{p}K\mathrm{a}_4).$$

Equation 9 is the same as the governing equation used in ref. (37) (although note that they are missing a factor of ln(10) in their manuscript); this is the equation used to calculate $E^0_{\mathrm{net}}$ values for all quinones presented in this study.

Solving Eq. 9 requires knowledge of $\Delta G^0_{r,i}$ for each reaction presented in Eqs. 1–4. Because these values are difficult or impossible to determine experimentally for some of our quinones of interest (particularly mPQ), we estimate them computationally using the ORCA quantum chemistry program package(38). That is, for each quinone of interest—including the reference 1,4-benzoquinone—we determine the optimum geometry and calculate the frequencies for each compound shown in the following scheme:

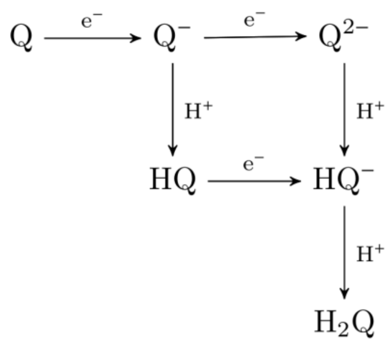

Specifically, we performed calculations by density functional theory (DFT) using the B3LYP functional (39, 40) and the 6-31G(d) basis set(41, 42); this combination of theory and basis set was chosen to balance computational efficiency and solution accuracy. In contrast, more accurate basis sets such as 6-31++G(d,p) (refs. (41, 42)) are computationally prohibitive for larger molecules, including the 9-isoprenoid side-chain $mPQ_{9:9}$, $PQ_{9:9}$, and $UQ_{9:9}$ compounds included here. We additionally calculated results for a "training set" of compounds whose p*K*a and $E^0$ values have been measured experimentally (2-methyl-1,4-benzoquinone; 2,3-dimethyl-1,4-benzoquinone; trimethyl-1,4-benzoquinone; tetramethyl-1,4-benzoquinone; 2-methoxy-1,4-benzoquinone; and 2,6-dimethoxy-1,4-benzoquinone) in order to generate scaling factors and to assess the impact of methyl and methoxy groups on the resulting $E^0_{\mathrm{net}}$ values.

Following Huynh et al.(37), all geometries were optimized in water using the conductor-like polarizable continuum solvation model (CPCM)(43) with Bondi atomic radii(44) and including non-electrostatic contributions of dispersion, repulsion, and cavitation energies(45). Thermochemistry results were calculated at $T$ = 298.15 K using reported Gibbs free energies that include the sum of zero-point energy, electronic, entropic, and solvation effects. Gibbs energy change for each reaction in Eqs. 3, 6 were then calculated as $\Delta G^0_r = \sum_p G^0 - \sum_r G^0$, where "$p$" represents the products, "$r$" represents the reactants, and $G^0$ is the Gibbs free energy for each compound as calculated by Gaussian. All calculations were performed on the Euler cluster operated by the High Performance Computing Group at ETH Zurich.

To estimate absolute p*K*a and $E^0_{net}$ values for each quinone of interest, we scaled all computational results to experimentally determined values using a training set (2-methyl-1,4-benzoquinone; 2,3-

dimethyl-1,4-benzoquinone; trimethyl-1,4-benzoquinone; tetramethyl-1,4-benzoquinone; 2-methoxy-1,4-benzoquinone; and 2,6-dimethoxy-1,4-benzoquinone). All experimental values are reported in Huynh et al.(37) and references therein. Such scaling factors are commonly used in computational chemistry to account for biases that arise due to incomplete functionals and basis sets. Resulting calculated *vs.* experimental $E^0_{net}$ plots are shown in Fig. S17; as expected, calculated $E^0_{net}$ exhibits a strong, positive correlation with experimentally measured $E^0_{net}$ values. All calculated values were thus scaled using the orthogonal distance regression (ODR) lines in Fig. S17 and uncertainty is taken as the regression root mean square error (RMSE). All computed, experimental, and scaled results are reported in Supplementary Datafile S4.

### Supplementary Results & Discussion

#### *Structural characterization of methyl plastoquinone*

Mass spectrometric characterization of mPQ revealed fragmentation spectra similar to PQ but with a base peak at *m/z* 165.0923 ($[C_{10}H_{12}O_2+H]^+$, theoretical mass 165.0910) instead of 151.0759 for PQ ($[C_9H_{10}O_2+H]^+$, theoretical mass 151.0754). This mass reflects a distinctive trimethyl-benzoquinone moiety connected to an isoprenoid tail (Fig. 1, Fig. S1, Table S1). Labeling experiments with methyl-$^2H_3$ methionine confirm that the three methyl groups must be located at C2, C5, and C6 of the benzoquinone moiety (Fig. S2). Specifically, mPQ with nine $^2H$ is observed in these experiments. This shows that the methyl groups are introduced at different positions because sequential methylation at the same ring position (i.e., formation of ethyl or propyl substituent) would have led to the loss of one or two $^2H$ during the formation of ethyl or propyl, respectively. Thus, mPQ is structurally related to UQ, by methylation at C2, and to PQ, by methylation at C5 and C6 of the benzoquinone moiety. Isoprenoid tails of mPQ varied in length (7 to 13 prenyl units) and degree of saturation (fully unsaturated to partly saturated) depending on the species (Fig. 1, Fig. S1). Previous misidentification of mPQ as MK in *Leptospirillum ferrooxidans*(46) likely resulted from identical retention times of MK and mPQ during HPLC analysis with UV detection, which we resolved by using HPLC-MS (Fig. 1).

The structure of mPQ was further confirmed using NMR spectroscopy. As mPQ could not be produced in quantities sufficient for NMR analysis from *Nitrospirota* cultures, $mPQ_8$ was produced in

*E. coli* Δ*ubiJ* by co-expression of *mpqQ* from *N. inopinata* and *plqH*, a C1-hydroxylase involved in PQ biosynthesis in the cyanobacterium *Synechnococcus* PCC6803 (Beltran *et al.*, in preparation). $mPQ_8$ was purified to homogeneity and was characterized by mass spectrometry and NMR. High-resolution tandem mass spectrometry shows identical elution times, exact masses, and fragmentation spectra for $mPQ_8$ from *E. coli* and *N. moscoviensis* (Fig. S3). In addition to the mass spectrometry data, the presence of an additional methyl group at the C2 position of the PQ scaffold can be constrained by NMR (Fig. S4). The first evidence is the absence of the signal of the proton linked to the C-2 of the quinone moiety which should appear above 6.65 ppm ($^1$H-NMR). Second, the presence of a third methyl group linked to the quinone moiety can be counted among the proton signals at 2.06-2.01 ppm. The presence of three methyl groups linked to the quinone moiety is confirmed by $^{13}$C NMR where three signals are observed at 11.14, 11.33, and 11.36 ppm. In conclusion, NMR characterization confirms the structure of the trimethyl headgroup of mPQ.

***Additional discussion on heterologous expression and isotope labeling experiments***

<u>Chorismate-pyruvate lyase:</u> The lack of *mpqC* in all *Nitrospira* spp. except *N. nitrificans* suggests the existence of alternative 4-HBA biosynthesis pathways or alternative ring precursors. In the PQ and UQ pathways, 4-HBA is generated from chorismate by the chorismate-pyruvate lyase UbiC(47). However, an alternative pathway to 4-HBA biosynthesis (e.g., XanB2)(48) or the use of substitute ring precursors (e.g., tyrosine, homogentisate) are known from bacteria and eukaryotes(49–51). The alternative XanB2 chorismate-pyruvate lyase is present in only two mPQ-producing species. Tyrosine could be used as a ring precursor in *Nitrospirota* lacking UbiC and XanB2, analogous to PQ biosynthesis in eukaryotes(52). Our experiments with *Nitrospirota* lacking UbiC, XanB2, or both, supplemented with ring-$^{13}C_6$-labeled substrates demonstrate that 4-HBA, but not tyrosine, is the ring precursor in *Nitrospirota* (Fig. S7). We therefore conclude that the use of 4-HBA as mPQ precursor is evolutionarily conserved but that alternative pathways to 4-HBA besides MpqC and XanB2 must exist in *Nitrospirota*.

*<u>Methyltransferases:</u>* The *plqQ* gene from *Synechocystis* sp. PCC6803 encodes a SAM-dependent methyltransferase proposed to perform the methylations at C5 and C6 in the PQ pathway, but this has

never been clearly demonstrated(53–55). These methylation steps do not exist in the UQ pathway, which instead has methoxy groups at C5 and C6 formed via hydroxyl intermediates (Fig. 3a). An *E. coli* mutant lacking the C5- and C6-hydroxylases (Δ*ubiIF*) accumulated 3-octaprenyl-4-hydroxyphenol ($4\text{-}HP_8$; Fig. S8a)(56), which has the same structure as the substrate of PlqQ in the proposed cyanobacteriotal PQ pathway (Fig. 3a). In the Δ*ubiIF* mutant, we indeed detected 3-octaprenyl ortho-quinone ($OQ_8$), the oxidized form of $4\text{-}HP_8$ (Fig. S8b). By contrast, expression of the methyltransferase (LFE_2124) from the four-gene cluster in *Leptospirillum* spp. and its homologues in *N. moscoviensis* and *M. noduliformans*, did not result in methylation of $PQ_8$ or $mPQ_8$ in Δ*ubiIF* and Δ*ubiIFH* mutants (Fig. S9).

Although *mpqQ* is missing from many *Nitrospirota* genomes, isotope labeling experiments indicate that these bacteria must contain functionally equivalent methyltransferases. In *N. moscoviensis*, which lacks *mpqQ*, supplementation of methyl-$^2H_3$-labeled methionine generates additional mPQ ions with +3, +6, and +9 Da in the benzyl moiety, indicating that all hydrogens of the methionine methyl groups are transferred intact to form the three methyl groups of mPQ (Fig. S2). This argues for a SAM-dependent methyltransferase similar to PlqQ/MpqQ and against an alternative reaction mechanism via radical SAM methyltransferases, which would subtract one $^2H$ from the methionine methyl group, leading to increases of +2, +4, and +6 Da. Finally, we took advantage of the accumulation of $PQ_8$ in the Δ*ubiIFE* mutant expressing *mpqQ* to test the activity of the *Nitrospirota ubiE* homologs (*mpqE*). Expression of *mpqE* from *L. ferrooxidans*, *N. moscoviensis*, and *M. noduliformans* led to the accumulation of $mPQ_8$ (Fig. 3e, S8c), demonstrating that all three proteins catalyze the C2-methylation of $PQ_8$ to $mPQ_8$.

***Occurrence of aerobic metabolism and HPQs in Cyanobacteria, Nitrospirota, and Pseudomonadota***

The distribution of quinone types and quinone biosynthesis genes has so far not been systematically studied in *Cyanobacteriota*, *Nitrospirota*, and *Pseudomonadota*. Here, we compiled literature data on quinone occurrence in these phyla, new analyses of quinone occurrence from basal *Cyanobacteriota* (*Gloeobacter violaceus* and *Anthocerotibacter panamensis*) and *Nitrospirota* (9 species), and inferred quinone type from the presence of key biosynthesis genes in 430 of 568 genomes from diverse archaeal and bacterial species (Fig. 2; Supplementary Datafile S3). While the occurrence of UQ in *Pseudomonadota* has been extensively studied, PQ has been experimentally confirmed only in late-branching cyanobacteriotal lineages (Supplementary Datafile S3). It has been suggested that the earliest-branching cyanobacteriotal lineage, *Gloeobacterales*, may not produce PQ due to the lack of some PQ biosynthesis genes(57). Our quinone analysis of two cultivated species of *Gloeobacterales*, *Gloeobacter violaceus* and *Anthocerotibacter panamensis*, demonstrates that PQ is indeed found in this lineage (Fig. S10). Our compilation of quinone occurrence data and biosynthesis gene inventories further suggests that HPQ biosynthesis is conserved not only in *Cyanobacteriota* but also in *Pseudomonadota* and aerobic *Nitrospirota* (Fig. S11-13). Further, HPQs or the required biosynthetic genes are not found outside these three phyla.

Representatives of the clades most closely related to *Cyanobacteriota* (the non-photosynthetic "*Candidatus* Margulisbacteria", "*Candidatus* Sericytochromatia", and *Vampirovibrionophyceae,* which are in the literature variably considered as separate phyla or classes of the candidate phylum "Melainobacteriota"), *Pseudomonadota* (*Bdellovibrionota*, *Campylobacterota*, *Desulfobacterota*, *Myxococcota*), and aerobic *Nitrospirota* (*Thermodesulfovibrionales* within *Nitrospirota*; "*Candidatus* Methylomirabilota", *Nitrospinota*) all are predicted to produce MK via the $MK_{mqn}$ pathway. $MK_{mqn}$ biosynthesis genes were consistently found within the *Vampirovibrionophyceae* (5/6 genomes) but were fewer in "*Candidatus* Margulisbacteria" (2/18) and "*Candidatus* Sericytochromatia" (2/15). By contrast, the MK derivative phylloquinone (PhQ, $MK_{4:1}$) is produced by all *Cyanobacteriota* (*sensu stricto*) through the $MK_{men}$ pathway (Table S5; Fig. 2). This split between $MK_{men}$ in *Cyanobacteriota* and $MK_{mqn}$ in closely related clades suggests that the $MK_{men}$ or the $MK_{mqn}$ pathway was laterally transferred after the divergence of *Cyanobacteriota* from their non-photosynthetic relatives. MK is also produced by some *Pseudomonadota* via the $MK_{men}$ pathway (Table S5). Due to the fragmentary distribution of this pathway in *Pseudomonadota* and the lack of MK in early branching lineages (Fig. 2) as well as the presence of the $MK_{mqn}$ pathway in the most closely related phyla, it appears most likely that the $MK_{men}$ pathway was acquired after the emergence of the UQ pathway and the radiation of crown group *Pseudomonadota*. As discussed below, biosynthesis proteins of the MK pathways found in

*Cyanobacteriota*, *Nitrospirota*, and *Pseudomonadota* form sister lineages to the HPQ proteins and thus do not suggest that the HPQ pathways evolved from the MK pathways in these three phyla or other closely related phyla.

Aerobic metabolism is common in *Cyanobacteriota*, *Nitrospirota*, and *Pseudomonadota* while anaerobic metabolism is rare. In *Pseudomonadota,* 75 of the 128 surveyed species are aerobes, 41 are facultative anaerobes, and 9 are strict anaerobes (Fig. 2; Supplementary Datafile S3). All described isolates of the basal clades *Magnetococcia* and *Zetaproteobacteria* are obligate aerobes, although genomes indicate that some strains could be facultative anaerobes(58, 59). Within the Alpha-, Beta-, and *Gammaproteobacteria*, some species are facultative anaerobes and few are anaerobes. In agreement with the dominance of aerobic metabolisms, 105/128 species use UQ as sole quinone, 19/128 use MK and UQ, and only 4/128 strains use MK as sole quinone. This indicates that the use of UQ (or the derivative rhodoquinone) for anaerobic respiration is common in *Pseudomonadota*, in support of earlier studies(11, 60). Given the prevalence of UQ and aerobic respiration in *Pseudomonadota*, particularly in basal clades, we suggest that their last common ancestor was an aerobe using UQ. The use of strictly anaerobic metabolisms may have evolved later and in some cases included lateral acquisition of the $MK_{men}$ pathway. Similarly, all described *Cyanobacteriota* (*sensu stricto*, Oxyphotobacteria) including the basal *Gloeobacterales* are aerobes capable of oxygenic photosynthesis (Fig. 2; Supplementary Datafile S3), with the exception of some obligate endosymbiotic *Cyanobacteriota*(61, 62). Likewise, aerobic respiration is universally conserved in *Cyanobacteriota*(63), suggesting that their last common ancestor was an aerobe using PQ. In *Nitrospirota*, basal clades are not well resolved (Fig. S11). However, the basal *Thermodesulfovibrionia* are anaerobes using MK and some basal *Nitrospiria* MAGs may be anaerobes(64), which suggests that aerobic respiration using mPQ only evolved after the divergence of the classes *Thermodesulfovibrionia* and *Nitrospiria*.

***Phylogenetics of quinone biosynthesis proteins***

The following sections expand on the description of the function and phylogeny of quinone biosynthesis proteins provided in the main text. Specifically, the following hypotheses are evaluated: 1) that the three HPQ pathways share a common origin, 2) that HPQ biosynthesis evolved prior to the radiations of *Cyanobacteriota*, *Pseudomonadota*, and *Nitrospirota*, 3) that the HPQ pathways are closely related to, but not derived from, the classical ($MK_{men}$) or futalosine pathways ($MK_{mqn}$) for MK biosynthesis.

It has previously been proposed that HPQs originated in *Cyanobacteriota* and that the UQ pathway originated from lateral transfer of PQ pathway genes from *Cyanobacteriota* to *Pseudomonadota*(65). Further, it has been argued that genes of the PQ pathway were laterally transferred from *Cyanobacteriota* to *Pseudomonadota* twice, once to Alphaproteobacteria and once to Zetaproteobacteria(65). Finally, it has been argued that the HPQ pathways were derived from parts of the $MK_{men}$ and $MK_{mqn}$ pathways(66). Our analyses do not support these arguments as homologs of HPQ biosynthesis proteins in *Cyanobacteriota* form a sister lineage to *Pseudomonadota* and all HPQ pathway proteins form sister lineages to homologous proteins of the MK biosynthetic pathways (as described in detail below).

*Prenyltransferase UbiA/PlqA/MpqA:* The *ubiA* gene of *E. coli* encodes a prenyltransferase that attaches a polyprenol side chain to 4-HBA during ubiquinone biosynthesis(67). Homologs of *ubiA* are found in *Cyanobacteriota* (*plqA*) and *Nitrospirota* (*mpqA*). The *plqA* gene from *Synechocystis* PCC6803 has previously been shown to complement *E. coli* Δ*ubiA* mutants(25). Similarly, the *mpqA* homologs from three tested *Nitrospirota* complement an *E. coli* Δ*ubiA* mutant (Fig. S6). Labeling experiments implicate 4-HBA as the prenyltransferase substrate in *Synechocystis* PCC6803 (ref. (25)) and *Nitrospirota* (Fig. S7). This 4-HBA prenylation activity of UbiA/PlqA/MpqA is distinct from quinone prenyltransferases of the MK pathways, which are not able to prenylate 4-HBA(68). This indicates that the HPQ prenyltransferases perform a conserved function and may be evolutionarily related.

Phylogenetic analyses support the close relationship of HPQ prenyltransferases. The HPQ prenyltransferases are monophyletic with regard to other prenyltransferases of the UbiA family (Fig. 4, S14, S18). This protein family comprises prenyltransferases with diverse functions, including biosynthesis of heme O and heme A (COX10), MK biosynthesis (MenA: $MK_{men}$ pathway; MqnP: $MK_{mqn}$ pathway)(68), bacteriochlorophyll (BchG), tocopherol/plant PQ (HPT), and archaeal membrane lipids (DGGGPS). The HPQ prenyltransferases are most closely related to the MqnP sister lineage, while both are only distantly related to MenA. Within the HPQ prenyltransferases, UbiA and MpqA form a sister lineage of PlqA (Fig. 4). Based on the prenyltransferase tree topology, we conclude that the HPQ pathways did not evolve from the extant MK pathways. The split of HPQ and LPQ pathways may be as old as the $MK_{mqn}$ pathway and therefore may be as old as the first emergence of MK. The closest relatives of each *Cyanobacteriota* ("*Candidatus* Margulisbacteria", "*Candidatus* Sericytochromatia", *Vampirovibrionophyceae*), *Pseudomonadota* (*Campylobacterota*, *Myxococcota*, *Bdellovibrionota*, *Desulfobacterota*), and aerobic *Nitrospirota* (*Thermodesulfovibrionales* within *Nitrospirota*; "*Candidatus* Methylomirabilota", *Nitrospinota*) all use the $MK_{mqn}$ pathway (Table S5). Their MqnP homologs cluster with other MqnP sequences, and not with HPQ prenyltransferases. We thus suggest that HPQ prenyltransferases did not originate in any of the extant lineages of *Cyanobacteriota*, *Pseudomonadota*, and aerobic *Nitrospirota* including their stem groups. Instead, HPQ methyltransferases may have originated in a yet undescribed or extinct lineage and were laterally transferred to *Cyanobacteriota*, *Pseudomonadota*, and aerobic *Nitrospirota*. Since HPQs are ancestral features of these phyla, any lateral transfer between them must have occurred before their radiation.

<u>Chorismate pyruvate-lyase UbiC/PlqC/MpqC:</u> The *ubiC* gene of *E. coli* encodes for a chorismate pyruvate-lyase that catalyzes the conversion of chorismate to 4-HBA(69), which is the ring precursor of UQ. This step is commonly regarded as the first committed step of UQ biosynthesis(70), although it has been suggested that 4-HBA may also serve other functions(71). Homologs of *ubiC* occur in many but not all UQ-producing *Pseudomonadota*(48, 70). The alternative chorismate lyase XanB2 has been shown to be present in *Pseudomonadota* species that lack *ubiC*(48). Homologs of *ubiC* (*plqC*) have been found in many *Cyanobacteriota* but lack in others, such as *Gloeobacter* species. The *plqC* gene of the cyanobacterium *Synechocystis* sp. PCC6803 (*sll1797*) has been shown to be able to restore UQ biosynthesis in an *E. coli* Δ*ubiC* mutant and is required for PQ biosynthesis(72). Likewise, a *ubiC* homolog (*mpqC*) occurs in aerobic *Nitrospirota* and can complement an *E. coli* Δ*ubiC* mutant (Fig. 3b). While *mpqC* is found in all three described aerobic genera of *Nitrospirota* (*Leptospirillum*, *Manganitrophus*, *Nitrospira*), it is lacking from most *Nitrospira* spp. Only few *Nitrospira* spp. have the

alternative *xanB2*. Homologs of CH-IV, another recently proposed alternative gene found in some *Pseudomonadota*(71), are found in a few *Nitrospira* spp. lacking *ubiC*. Still, some *Nitrospira* spp. such as *N. moscoviensis* lack all of these genes (*mpqC*, *xanB2*, CH-IV) but still use 4-HBA as precursor via unknown alternative enzymes, as demonstrated by labelling experiments (see main text and Fig. S7). This demonstrates that despite the fragmented conservation of chorismate lyase homologs, the use of 4-HBA as quinone precursor is a conserved biochemical feature of *Pseudomonadota*, *Cyanobacteriota*, and *Nitrospirota*.

The phylogeny of UbiC/PlqC/MpqC is broadly coherent with those of the other HPQ biosynthesis genes that suggest a common origin (Fig. S19). The GntR transcriptional regulator from *Bacillus subtilis* was used as the outgroup. Similar to UbiC, GntR is part of the superfamily IPR028978, which shares a conserved domain with chorismate pyruvate-lyases. However, many branches are not well supported and *Nitrospirota* are split into two branches, possibly due to long branch attraction of the distantly related outgroup, the fragmented phylogenetic coverage, and low sequence similarity. Still the topology of the UbiC tree is broadly similar to those of the other HPQ genes: Apart from a few sequences, homologs from Cyanobacteriota form a divergent sister lineage to homologs from *Pseudomonadota* and *Nitrospirota*. The coherence of the UbiC/PlqC/MpqC phylogeny with those of other HPQ biosynthesis genes suggests that the fragmentary distribution of UbiC/PlqC/MpqC results from losses in multiple lineages.

Decarboxylase UbiDX/MpqDX: UbiD catalyzes the decarboxylation of 3-prenyl-4-HBA using a prenylated co-factor provided by UbiX(70, 73, 74). Homologs of the *ubiDX* genes are found in *Cyanobacteriota* and aerobic *Nitrospirota*. The homologs *plqD/plqX* (*sll0936*, *slr1099*) from *Synechocystis* sp. PCC6803(72) and *mpqD/mpqX* from *M. noduliformans* can complement *E. coli* Δ*ubiD* and Δ*ubiX* mutants (Fig 3d). Among aerobic *Nitrospirota*, only the *Manganitrophus* lineage contains *ubiDX*. However, these genes are also missing from some *Pseudomonadota*(65, 75), indicating that additional decarboxylases must exist(70). Homologs of the putative alternative decarboxylase *ubiZ*(66) are not present in any of the studied *Nitrospirota*. By contrast, *ubiDX* homologs (MqnL and MqnM) are present in most anaerobic *Nitrospirota*. MqnL and MqnM are likely involved in the $MK_{mqn}$

pathway, although this step has not yet been proven experimentally(70, 76). This could indicate a shared origin of *ubiDX* homologs in the HPQ pathways and the $MK_{mqn}$ pathway.

Phylogenetic analyses support the monophyly of UbiDX and do not support an origin from the $MK_{mqn}$ pathway. For these analyses, several UbiD-like (de-)carboxylases (Fdc1, SmdK, TtnD, PpcB, PhdA) and their associated UbiX-like prenyltransferases (Pad1, SmdJ, TtnC, PhdB) were used as outgroups. These proteins are related to *E. coli* UbiDX but catalyze the (de-)carboxylation of different substrates(77). The UbiD/MpqD/PlqD phylogeny supports the same branching pattern as UbiA/MpqA/PlqA: UbiD and MpqD are sister lineages to each other, the UbiD/MpqD form a sister clade to PlqD, and the UbiD/MpqD/PlqD clade is monophyletic with regard to the sister group consisting of MqnL (including MqnL of anaerobic *Nitrospirota*, and sister phyla of *Pseudomonadota* and *Cyanobacteriota*; Fig 3a, S15). This topology is broadly preserved in the UbiX/MpqX/PlqX phylogeny, but support is low for some branches, such as the MpqX and PlqX branches which cluster with some MqnM homologs (Fig. S20). The consistency between the UbiD/MpqD/PlqD and UbiA/MpqA/PlqA tree topologies suggests that the lack of UbiDX in *Leptospirillum* and *Nitrospira* spp. results from gene loss. The monophyly of UbiD/PlqD/MpqD separate from MqnL further supports the hypothesis that the HPQ pathways share a common origin and did not originate from the extant $MK_{mqn}$ pathway.

<u>*C2 methyltransferase UbiE/MpqE:*</u> In both MK and HPQ biosynthesis, methylation of the quinone ring moiety is a late step in the pathway. In the $MK_{men}$ pathway, MenG catalyzes methylation at the C2 position of the napthoquinone moiety and a homolog for the same step has been proposed for the $MK_{mqn}$ pathway (MqnK)(76). In *Pseudomonadota*, UbiE catalyzes methylation at the C2 position of the benzoquinone moiety to form UQ. However, in *E. coli* (and likely other *Pseudomonadota*), the C2 methyltransferase UbiE is bifunctional and can also methylate MK (derived from the $MK_{men}$ pathway in *E. coli*). A homologous methyltransferase found in *Cyanobacteriota* and plastids is strictly monofunctional, methylating only MK but not PQ(78). Homologs of UbiE (MpqE) are found in all aerobic *Nitrospirota*. Expression of MpqE from several aerobic *Nitrospirota* in an *E. coli* Δ*ubiE* mutant restores the synthesis of MK and UQ, indicating that MpqE is bifunctional (Fig. S7). Previously it has

been suggested, based on protein phylogenies, that UbiE of *Pseudomonadota* is more closely related to the $MK_{men}$ pathway (MenG) than the $MK_{mqn}$ pathway (MqnK) and that it was derived from those pathways(66).

Here, we re-analyzed the phylogeny of C2 methyltransferases, finding that the HPQ C2 methyltransferases did not originate from any of the two MK pathways and were likely lost in *Cyanobacteriota* after the divergence of HPQ pathways. To broadly explore the sequence space of quinone methyltransferases, sequences covering all known HPQ methyltransferases (UbiE, MpqE, MpqQ, PlqQ) were used as blastp queries. Phylogenetic trees were constructed using the phosphatidylethanolamine methyltransferase PmtA as an outgroup that is distantly related to both UbiE/MpqE/MqnK/MenG as well as MpqQ/PlqQ. Due to the large number of hits (>800) a preliminary tree was generated using Fasttree. For the final alignment and tree, only the UbiE/MpqE, MqnK, and MenG clusters and the deeply branching neighbor lineages were retained (Fig. S16). The phylogenetic analysis shows that UbiE/MpqE are more closely related to MqnK ($MK_{mqn}$ pathway) than to MenG ($MK_{men}$ pathway), while MqnK and UbiE/MpqE form sister lineages. Therefore, the earlier hypotheses that HPQ C2 methyltransferases evolved from one of the MK pathways is not supported. This suggests that the bifunctionality of UbiE/MpqE was already present before the pathway was transferred to *Pseudomonadota* and *Nitrospirota*. Further experiments with MqnK expressed in *E. coli ΔubiE* mutants are needed to confirm whether bifunctionality was already present earlier, before the divergence of MqnK and UbiE/MpqE and thus prior to the evolution of all HPQ pathways. The genes may then have been lost in *Cyanobacteriota* or was gained by *Pseudomonadota* and *Nitrospirota* after the divergence of the HPQ pathways. However, given the fact that a C2 methylase is present in all quinone biosynthetic pathways (except PQ) and in all domains of life, it appears plausible that *Cyanobacteriota* lost their C2 methylase homolog. This loss must have occurred after the divergence of HPQ pathways and before the radiation of crown group *Cyanobacteriota*. Since loss of C2 methylation is required for the functioning of photosystem II(78), this loss was likely linked to the evolution of oxygenic photosynthesis in *Cyanobacteriota*.

*C5/C6 methyltransferase MpqQ/PlqQ:* It has previously been suggested that the radical SAM methyltransferase Sll0418 (PlqQ) of *Synechocystis* spp. PCC6803 catalyzes methylation at the C5/C6 position to yield PQ in addition to its role in tocopherol biosynthesis(54, 55, 79). However, deletion of *plqQ* in *Synechocystis* spp. PCC6803 results in only slightly lower PQ levels(54, 55, 80). This indicates redundancy in quinone methyltransferases in *Cyanobacteriota*. Homologs of PlqQ (MpqQ) are found in a few species of *Nitrospirota*, limited to the genus *Nitrospira*. MpqQ from *N. inopinata* shows C5/C6 methyltransferase activity, resulting in the production of PQ and mPQ in *E. coli* Δ*ubiIF* and of PQ only in the Δ*ubiIFE* mutant (Fig. S8b). To broadly explore the sequence space of quinone methyltransferases, sequences covering all known HPQ methyltransferases (UbiE, MpqE, MpqQ, PlqQ) were used as blastp queries. Phylogenetic trees were constructed using the phosphatidylethanolamine methyltransferase PmtA as an outgroup that is distantly related to both UbiE/MpqE/MqnK/MenG as well as MpqQ/PlqQ. Due to the large number of hits (>800) a preliminary tree was generated using Fasttree. Besides the MpqQ/PlqQ cluster, no other clusters containing closely related sequences from *Nitrospirota* and *Cyanobacteriota* were found. For the final alignment and tree (Fig. 4b), only the MpqQ/PlqQ cluster and the deeply branching neighbor lineages were retained.

The MpqQ and PlqQ methyltransferases are not universally conserved. Only few *Nitrospira* spp., affiliated exclusively with *Nitrospira* sublineage II, possess an MpqQ homolog. These sequences are nested as a single cluster within sequences from *Cyanobacteriota* (Fig. 4b). The phylogeny thus suggests that the gene was derived by lateral transfer from *Cyanobacteriota* into sublineage II of the genus Nitrospira. However, only 11% (7 of 62) of the studied *Nitrospirota* genomes with mPQ pathways possess MpqQ. Similarly, only 56% (40 of 71) of the studied *Cyanobacteriota* genomes contain a PlqQ homolog, including basal lineages such as Gloeobacterales. We therefore suggest that the methylations at C5/C6 must be performed by alternative enzymes in the majority of *Nitrospirota* and *Cyanobacteriota*.

Identification of alternative enzymes will provide valuable information on the evolution of PQ and mPQ biosynthesis. We therefore tested a candidate alternative enzyme, a cobalamin-dependent methyltransferase, which is located next to the *mpqE* and *mpqA* genes in *L. ferrooxidans* (LFE2124;

Fig. S5). Homologs of the LFE2124 gene occur in most *Nitrospira* spp., in *Manganitrophus* spp., and all *Leptospirillum* spp. and are most closely related (~40% identity) to sequences from *Cyanobacteriota*, such as slr0320 from *Synechocystis* PCC6803. It was recently proposed that *slr0320* is involved in PQ biosynthesis, as deletion resulted in defects in electron transfer, although the authors did not analyze quinones to test this hypothesis(81). Heterologous expression of LFE2124 and the homologs from *N. moscoviensis* and *M. noduliformans* in *E. coli* Δ*ubiIF* or Δ*ubiIHF* mutants did not result in the methylation of 4-$HP_8$ (detected under its oxidized form $OQ_8$, Fig. S9a-b), suggesting that these enzymes are not involved in the mPQ biosynthetic pathway. Sufficient cobalamin should have been present in the LB medium used for these experiments. Alternatively, involvement of LFE2124 homologs in mPQ biosynthesis may rely on the supply of an unknown cofactor in addition to cobalamin. Given the current evidence, it cannot be established that the capacity for C5/C6 methyltransferase was uniformly, laterally transferred from *Cyanobacteriota* to *Nitrospirota*. However, alternative pathways for specific quinone biosynthesis steps have emerged across all domains of life(54, 66, 70, 82) while the chemical structure of the resulting quinone remained unchanged. Therefore, we suggest that C5/C6 methylation is a biochemically conserved step in PQ/mPQ biosynthesis and likely an ancestral feature of HPQs, although it is not performed by homologous enzymes across all extant PQ/mPQ producers.

<u>Quinone biosynthesis-related kinase UbiB/PlqB/MpqB:</u> UbiB is required for UQ biosynthesis in *E. coli*(83), where it is thought to be involved in the extrusion of UQ precursors from the membrane(84). UbiB homologs occur in *Cyanobacteriota* and *Nitrospirota*, but it remains unresolved if they have the same function as UbiB from *E. coli*. The *ubiB* homolog *mpqB* from *L. ferrooxidans* was not able to restore UQ biosynthesis in *E. coli ΔubiB* mutants (Fig. S9). Lack of function may indicate that MpqB and UbiB use distinct substrates, or, that MpqB is not involved in quinone biosynthesis.

Phylogenetic analyses of UbiB reveals a pattern distinct from the other studied quinone biosynthesis proteins. For these analyses, a selection of Rio1 kinase sequences was used as the outgroup. Rio1 kinases are distantly related to UbiB but both contain the same protein kinase-like domain (IPR011009). UbiB phylogenies are complicated by the existence of two or more homologs in many *Pseudomonadota* and *Cyanobacteriota* (but not *Nitrospirota*; Fig. S21). The cluster of sequences that contains the only two bacterial proteins with confirmed functionality, UbiB from *E. coli*(83, 84) and *Xanthomonas campestris*(85), also contains putative MpqB and PlqB homologs. In contrast to all other mPQ biosynthesis genes, the UbiB-like sequences of anaerobic *Nitrospirota* (*Thermodesulfovibrio*, *Sulfobium*, etc.; all of which use the $MK_{mqn}$ pathway) cluster together with MpqB of aerobic *Nitrospirota* (*Leptospirillum, Nitrospira*, *Manganitrophus* spp.; mPQ pathway). Similarly, UbiB-like sequences from "*Candidatus* Margulisbacteria", "*Candidatus* Sericytochromatia", and *Vampirovibrionophyceae* ($MK_{mqn}$ pathway) cluster with PlqB from *Cyanobacteriota* ($MK_{men}$ & PQ pathway). This suggests a vertical inheritance pattern with UbiB-like kinases being interchangeable between different quinone biosynthesis pathways, although this would need to be studied in more detail. The evolutionary history of UbiB family proteins thus appears unrelated to that of other HPQ biosynthesis genes. Due to the lack of confirmation of MpqB/PlqB function, we conclude that UbiB family proteins are currently not informative for understanding the evolution of HPQ pathways.

### Supplementary Figures

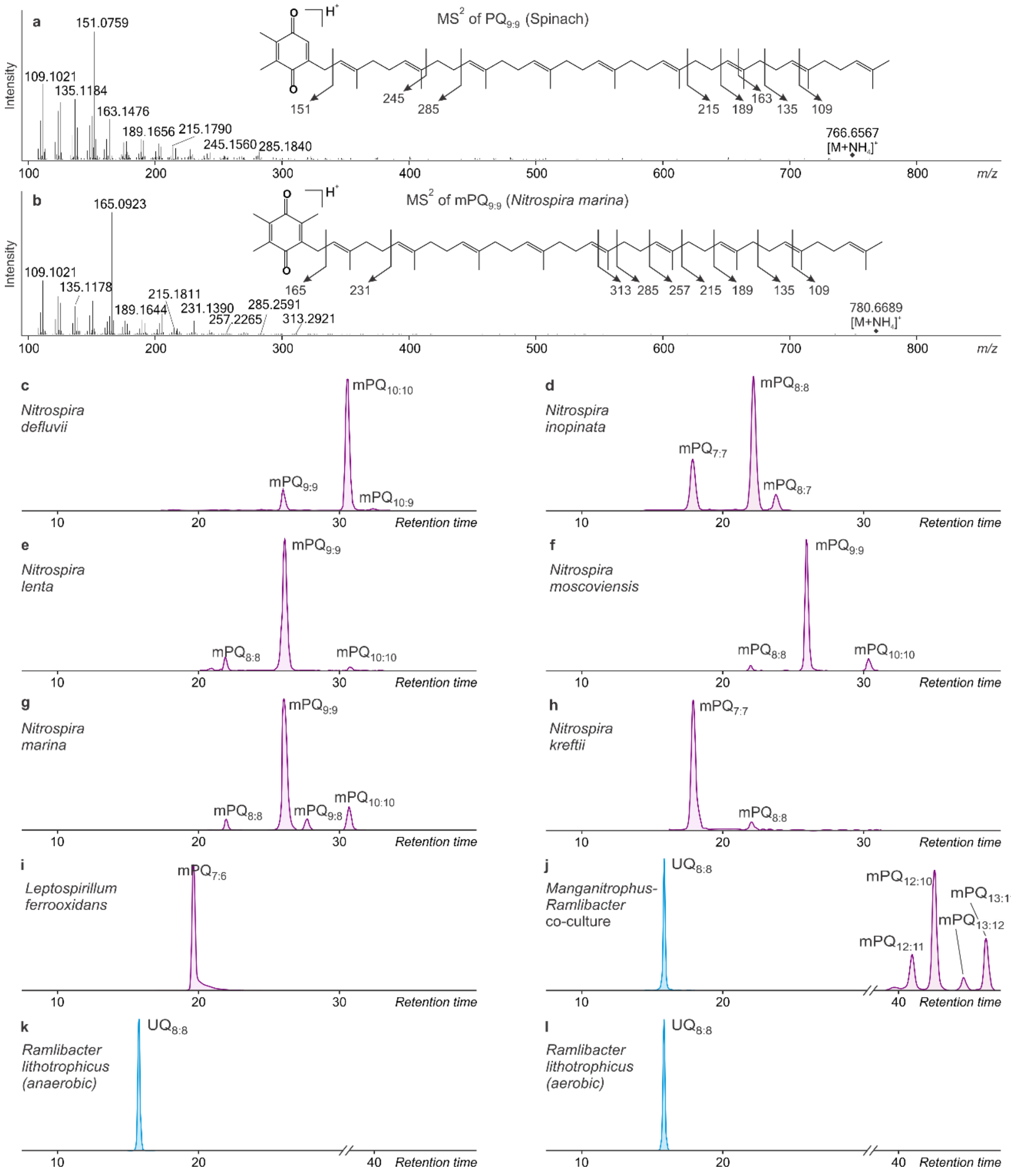

**Fig. S1.** Mass spectra and chromatogram of quinones in *Nitrospirota* and spinach (PQ$_{9:9}$). **a-b**, MS$^2$ fragmentation spectra and tentative structural characterization of PQ$_{9:9}$ ($NH_4^+$ adduct) and mPQ$_{9:9}$ ($NH_4^+$ adduct). Accurate masses and proposed sum formulas of characteristic fragment ions are shown in Table S1. **c-l**, Composite extracted ion chromatograms of respiratory quinones in extracts of *Nitrospira defluvii*, *N. inopinata*, *N. lenta*, *N. moscoviensis*, *N. marina*, *Ca.* N. kreftii, *Leptospirillum ferrooxidans*, a *Ca.* Manganitrophus noduliformans-*Ramlibacter lithotrophicus* co-culture, *R. lithotrophicus* (grown aerobically), and *R. lithotrophicus* (grown anaerobically).

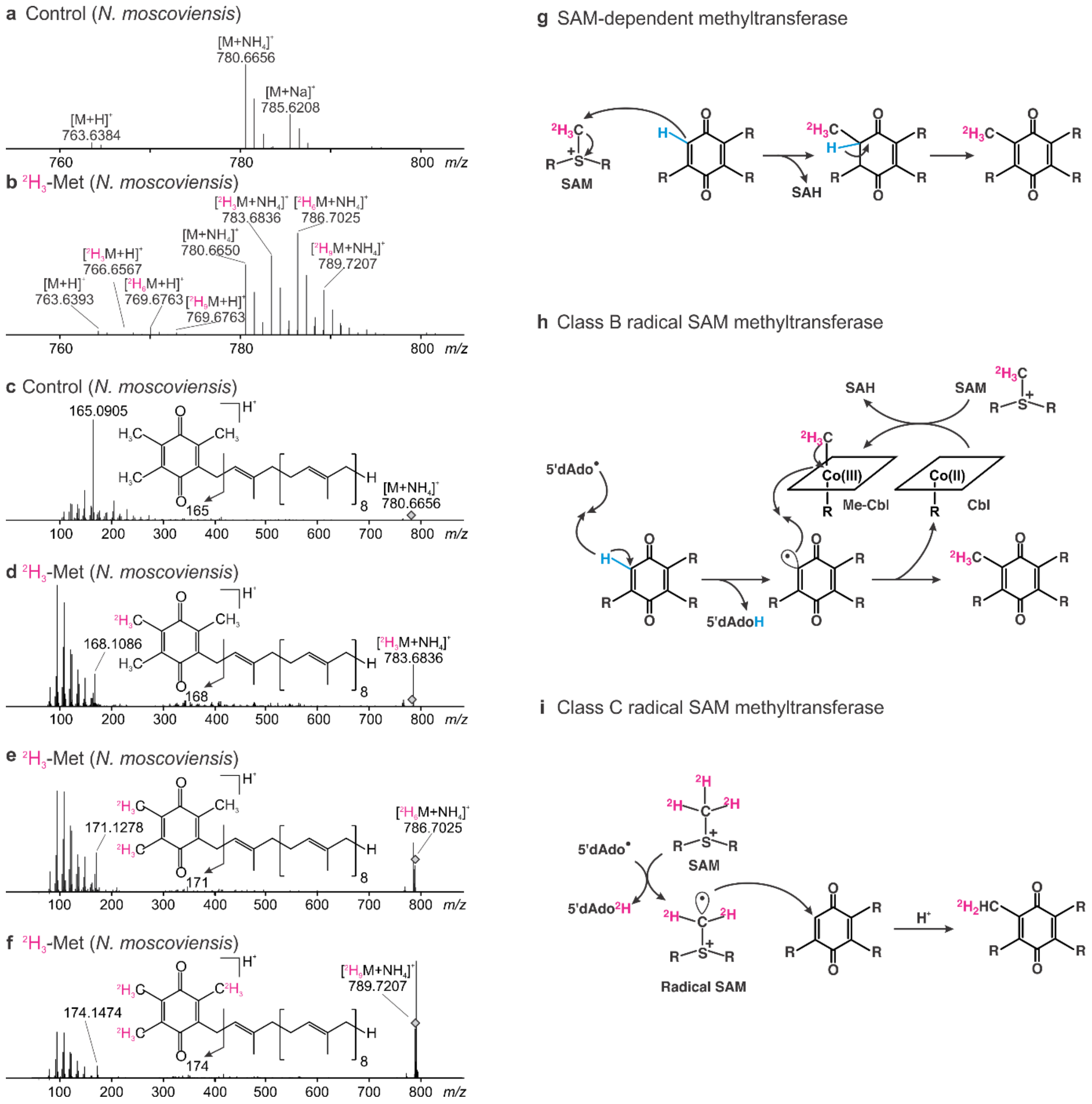

**Fig. S2. a-f**, Mass spectra of mPQ$_{9:9}$ in *Nitrospira moscoviensis* grown with methyl-$^2H_3$-methionine ($^2H_3$-Met), showing appearance of +3, +6, and +9 Da peaks as molecular ions and in the benzyl moiety fragments compared to the unlabeled control, indicating respectively the incorporation of one, two, and three intact $^2H_3$-methyl groups from methionine. Incorporation of three intact $^2H_3$-methyl groups from methionine indicates that the methyl groups must be located at C-2, C-5, and C-6 as methylation at any other position would result in loss of one or more $^2H$. **h-i**, Tentative reaction mechanisms for methylation at C5 and C6 of mPQ$_{9:9}$ (arbitrarily illustrated here for C6) for three types of methyltransferases. SAM-dependent methyltransferases and class B (cobalamin-dependent; Cbl) radical SAM methyltransferases are expected to preserve all three methyl-$^2H$ whereas class C radical

SAM methyltransferases are expected to lead to the loss of one methyl-$^{2}$H during the formation of the SAM radical. Preservation of all three methyl-$^{2}$H from methionine in $mPQ_{9:9}$ thus suggests that the enzyme responsible is a SAM-dependent methyltransferase or a class B radical SAM methyltransferase but not a class C radical SAM methyltransferase. *N. moscoviensis* was grown in a defined medium without cobalamin addition but may be able to produce cobalamin. However, the cobalamin-auxotroph *N. marina*(2) was grown without cobalamin addition but still produces mPQ, which does not support a class B radical SAM methyltransferase mechanism.

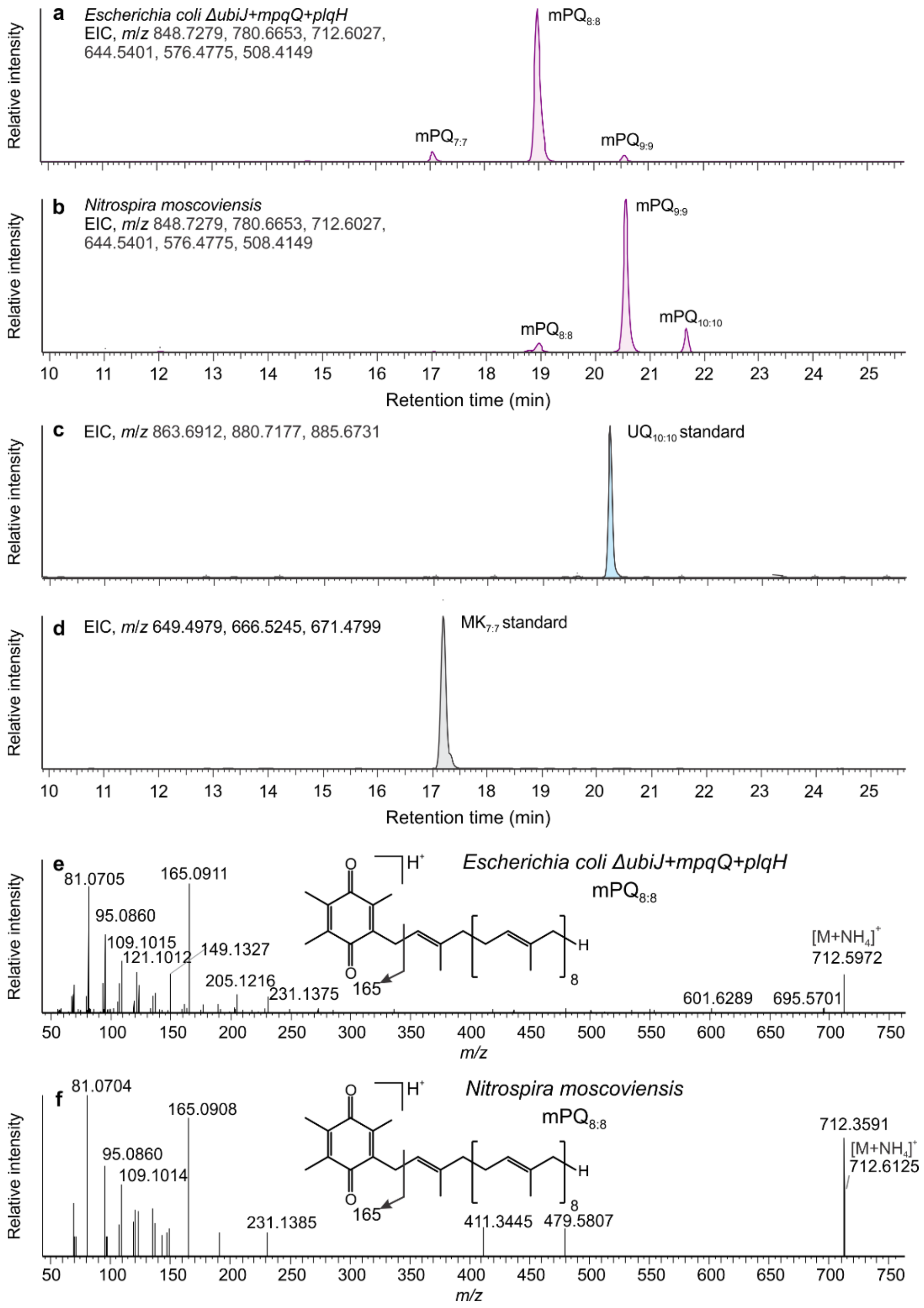

**Figure S3. a-b,** Extracted ion chromatograms (EIC) of mPQs ($[M+NH_4]^+$ ions) purified from *E. coli* Δ*ubiJ* expressing *mpqQ* from *N. inopinata* and *plqH* from *Synechococcus* PCC6803 (*E. coli* Δ*ubiJ+mpqQ+plqH*; **a**) and from extracts of *N. moscoviensis* (**b**) showing identical elution times of $mPQ_{8:8}$. $MK_{7:7}$ and $UQ_{10:10}$ standards are shown for reference (**c-d**). **e-f**, $MS^2$ fragmentation spectra of $mPQ_{8:8}$ from *E. coli* Δ*ubiJ+mpqQ+plqH* and *N. moscoviensis* showing identical exact masses of the molecular ions and identical fragmentation spectra (less than 10 mDa deviation from calculated exact mass).

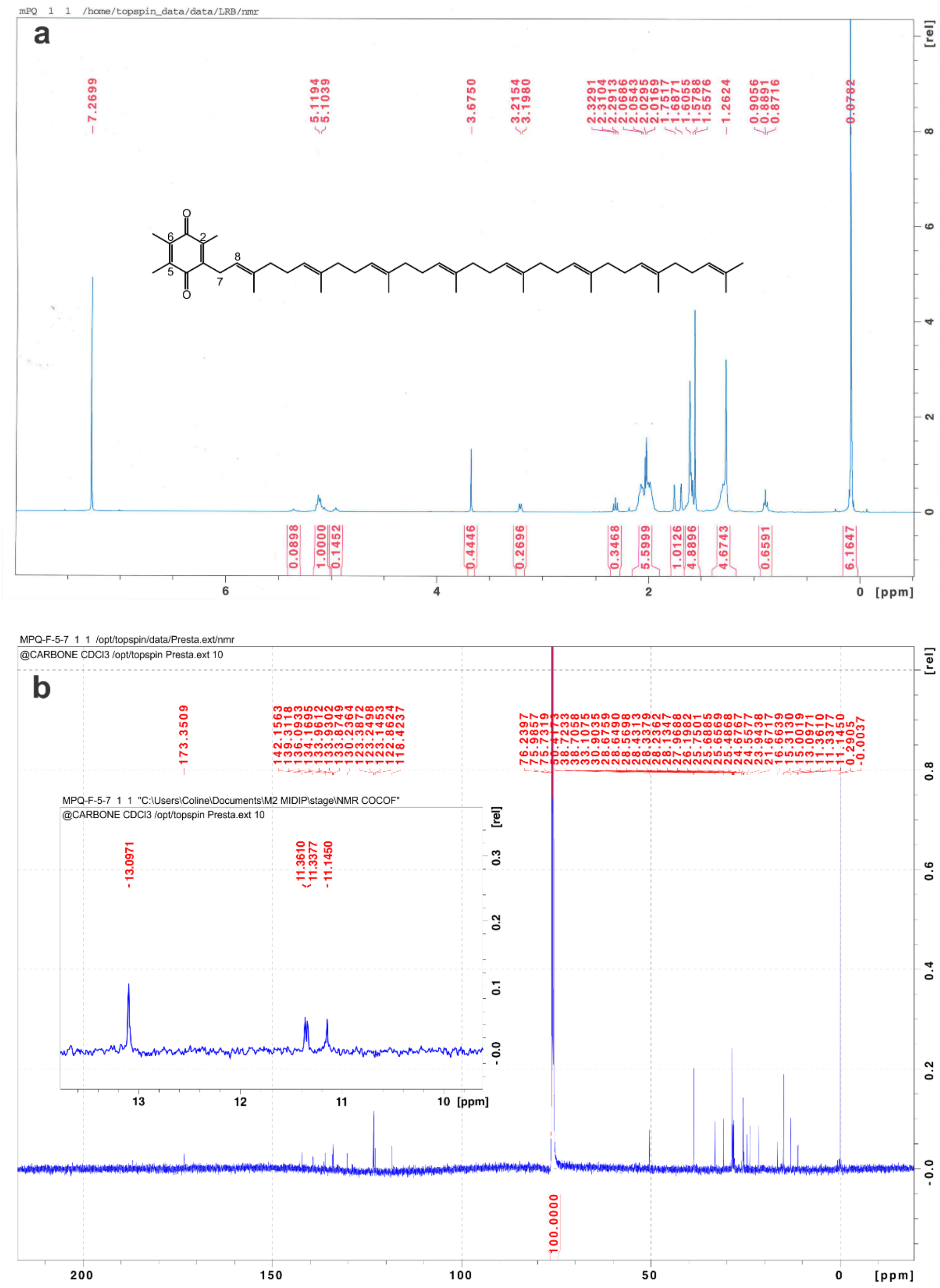

**Figure S4. a,** [1]H and **b,** [13]C NMR spectrum of $mPQ_8$ (in $CDCl_3$, 400 MHz for [1]H, 125 MHz for [13]C) purified from *E. coli* Δ*ubiJ* expressing *mpqQ* from *N. inopinata* and *plqH* from *Synechococcus* PCC6803. Inset in **b** is a close-up view of the range ~10-14 ppm showing the signals of the three methyl groups of the quinone moiety. Inset in panel **a** shows the structure of $mPQ_8$ with assigned carbon atoms numbered. δ: 5.25-5.0 (m, 7H, olefinic protons H); 4.95 (t, *J* = 6.9 Hz, H-8); 3.20 (d, *J* = 6.9 Hz, 2H, H-7); 2.06-2.01 (m, 41H, $-CH_2-CH_2$, $CH_3$ 2,5,6), 1.60-1.55 (m, isoprenyl $CH_3$).

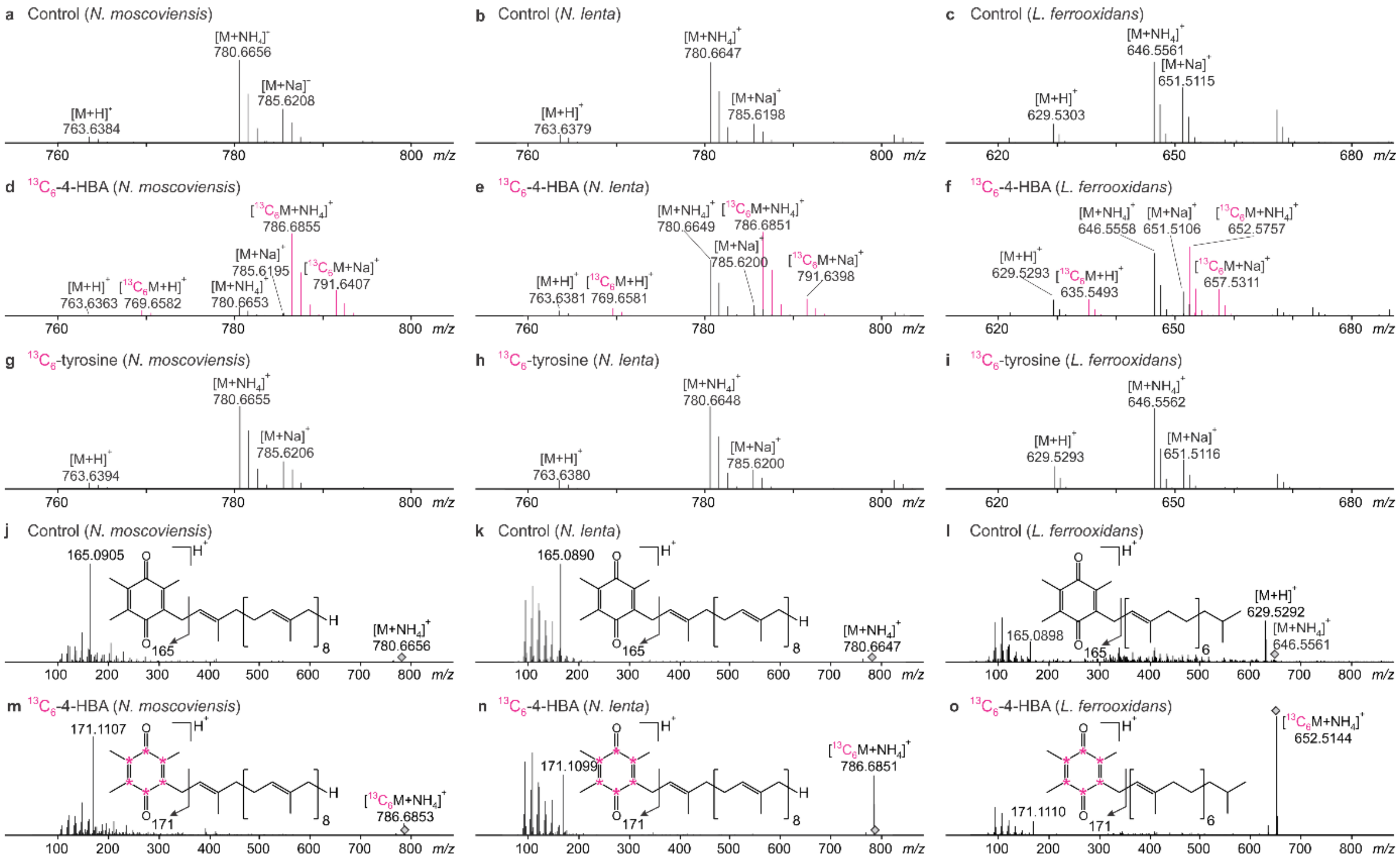

**Fig. S5.** $^{13}$C-labeling experiments demonstrating that 4-HBA, but not tyrosine, is the precursor of mPQ in *Nitrospirota* with or without known 4-HBA biosynthetic genes: *Nitrospira moscoviensis* lacks canonical chorismate-pyruvate lyase (*ubiC*) and the alternative *xanB2*; *N. lenta* lacks *ubiC* but has *xanB2*; *Leptospirillum ferrooxidans* has *ubiC* but lacks *xanB2*. *N. moscoviensis*, but not *N. lenta* and *L. ferrooxidans* species, possesses a 4-hydroxyphenylpyruvate dioxygenase homolog, which could be part of a pathway for biosynthesis of a ring precursor via tyrosine(52, 86), but this is not supported by the labeling experiments. **a-i**, MS spectra of $mPQ_{9:9}$ (*N. moscoviensis*, *N. lenta*) and $mPQ_{7:6}$ (*L. ferrooxidans*) showing appearance +6 Da peaks (pink) in cultures supplemented with ring-$^{13}C_6$-4-HBA (4-hydroxybenzoic acid) or ring-$^{13}C_6$-tyrosine, as well as control cultures grown without these substrates. **j-o**, $MS^2$ fragmentation spectra of $mPQ_{9:9}$ and $mPQ_{7:6}$ in the control and ring-$^{13}C_6$-4-HBA cultures showing incorporation of the intact ring moiety from ring-$^{13}C_6$-4-HBA (+6 Da) into the mPQ benzoquinone moiety. Diamonds in panels **j-o** indicate the precursor ion. Asterisks in panels j-o indicate tentative localization of $^{13}C$ in the benzyl ring moieties.

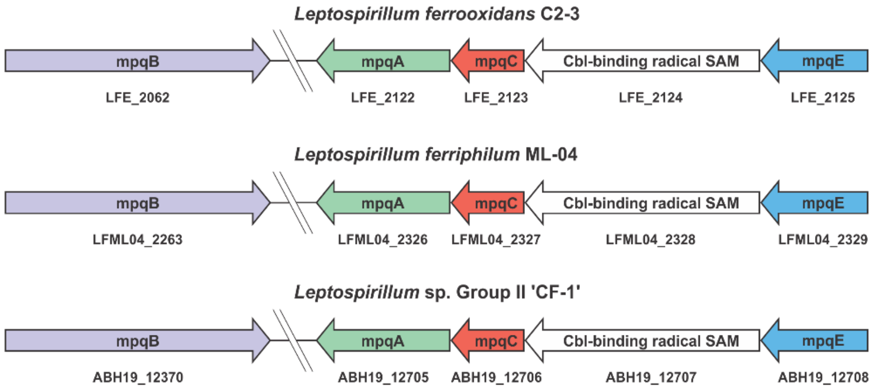

**Fig. S6.** Organization of mPQ biosynthesis genes in genomes of *Leptospirillum* spp. The genes are not co-localized in other *Nitrospirota*.

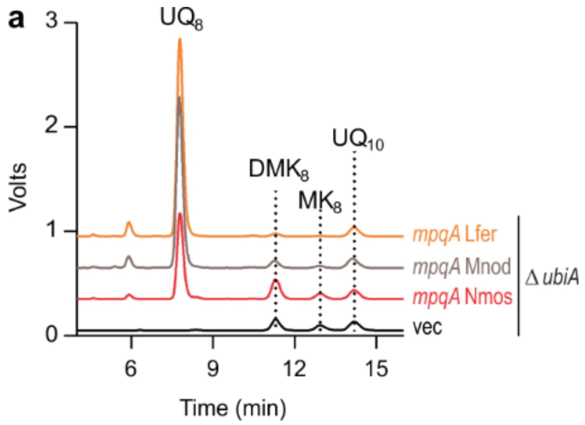

**Fig. S7.** HPLC-electrochemical detection of the total quinone content in lipid extracts of *E. coli* Δ*ubiA E. coli* mutants complemented with *mpqA* from *L. ferrooxidans* (Lfer), *N. moscoviensis* (Nmos) and *M. noduliformans* (Mnod) or containing an empty vector (vec). Representative chromatograms from 3 independent experiments. $DMK_8$ and $MK_8$ correspond respectively to dimethyl-menaquinone 8 and menaquinone 8. $UQ_{10}$ added as internal standard.

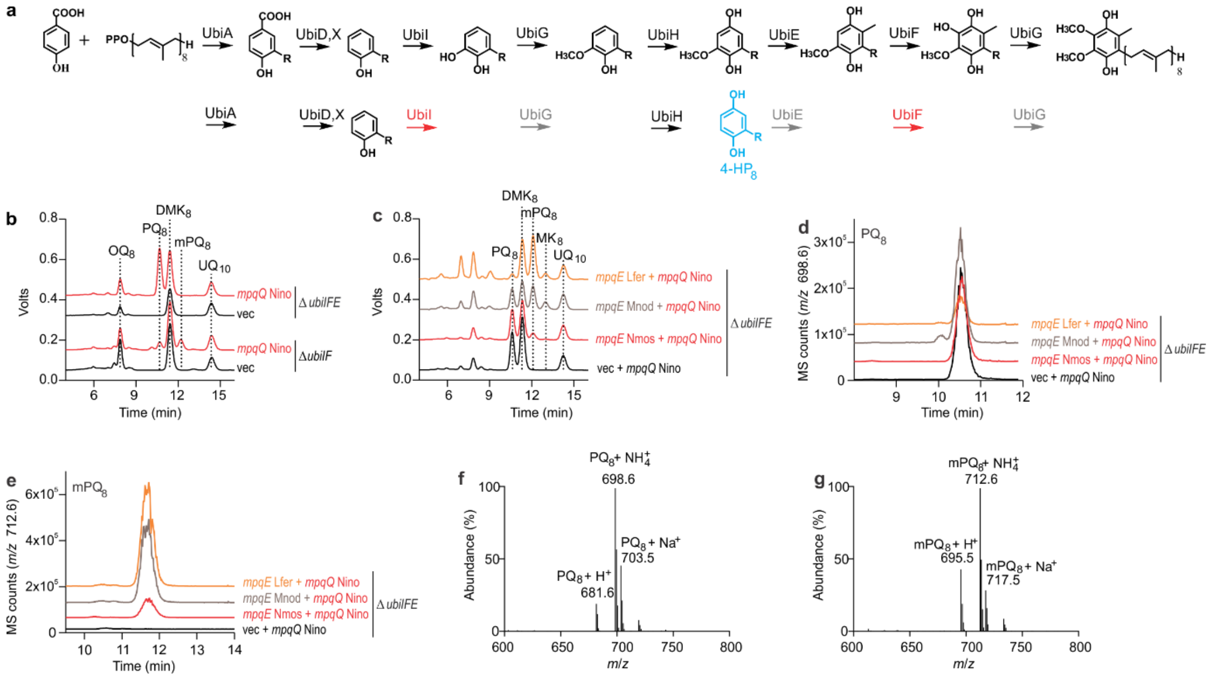

**Fig. S8.** Biosynthesis of PQ and mPQ in *E. coli*. **a**, *E. coli* UQ biosynthesis pathway (top) and steps which are inactivated (bottom) in the Δ*ubiIF* strain (grey/red for inactive/genetic deletion) leading to the accumulation of 4-$HP_8$ (blue); see ref. (87) for further details. **b**, HPLC-electrochemical detection (ECD) of the total quinone content in lipid extracts of *E. coli* Δ*ubiIF* and Δ*ubiIFE* mutants expressing *mpqQ* from *N. inopinata* (Nino) or empty vectors (vec). **c**, ECD chromatograms of HPLC-ECD-MS analyses of the total quinone content in lipid extracts of an *E. coli* Δ*ubiIFE* mutant with *mpqQ* from *N. inopinata* (Nino) in combination with *mpqE* from *L. ferrooxidans* (Lfer), *N. moscoviensis* (Nmos) and *M. noduliformans* (Mnod), or a vector with *mpqQ* from *N. inopinata*. **d-e** ion chromatograms and **f-g** mass spectra (main adducts $H^+$, $NH_4^+$ and $Na^+$ are labeled) of HPLC-ECD-MS analyses in panel **c**, detecting $PQ_8$ ($[M+NH_4]^+$, *m/z* 698.6; eluting at 10.7 min in panel **c**) and $mPQ_8$ ($[M+NH_4]^+$, *m/z* 712.6; eluting at 12.2 min in panel **c**). Representative chromatograms from 3 independent experiments. $UQ_{10}$ added as internal standard.

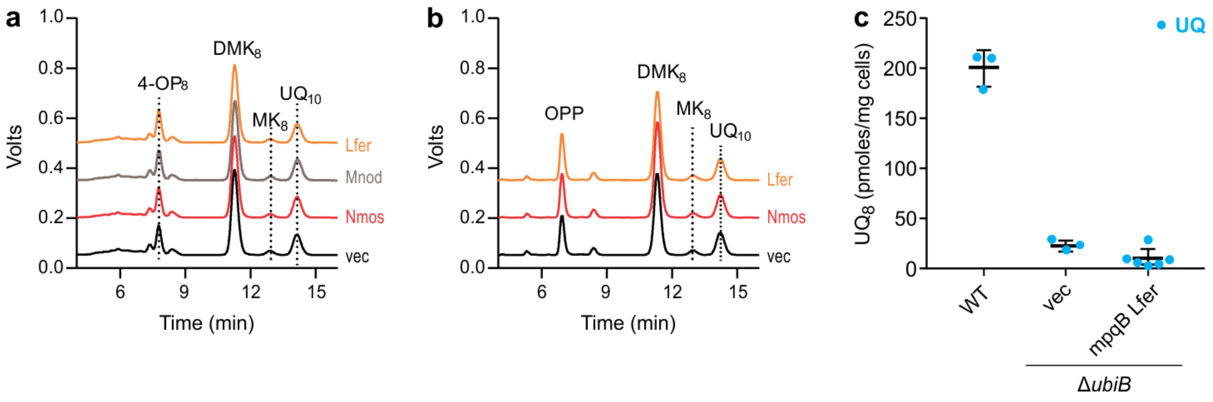

**Fig. S9. a-b**, HPLC-electrochemical detection of the total quinone content in lipid extracts from *E. coli* Δ*ubiIF* cells (**a**) or Δ*ubiIHF* cells (**b**) containing an empty vector (vec) or vectors coding for cobalamin-binding radical SAM methyltransferase from *L. ferrooxidans* (Lfer), *N. moscoviensis* (Nmos), and *M. noduliformans* (Mnod). Representative chromatograms are shown (*n*=2-3). **c**, $UQ_8$ quantifications from *E. coli* WT and Δ*ubiB* cells containing either an empty vector (vec) or a vector containing a putative *ubiB* homolog *mpqB* from *L. ferrooxidans* (means ± standard deviation, *n* = 3-6). The role of MpqB is unclear, similar to the unresolved role of UbiB in UQ and PQ biosynthesis(56, 83, 84).

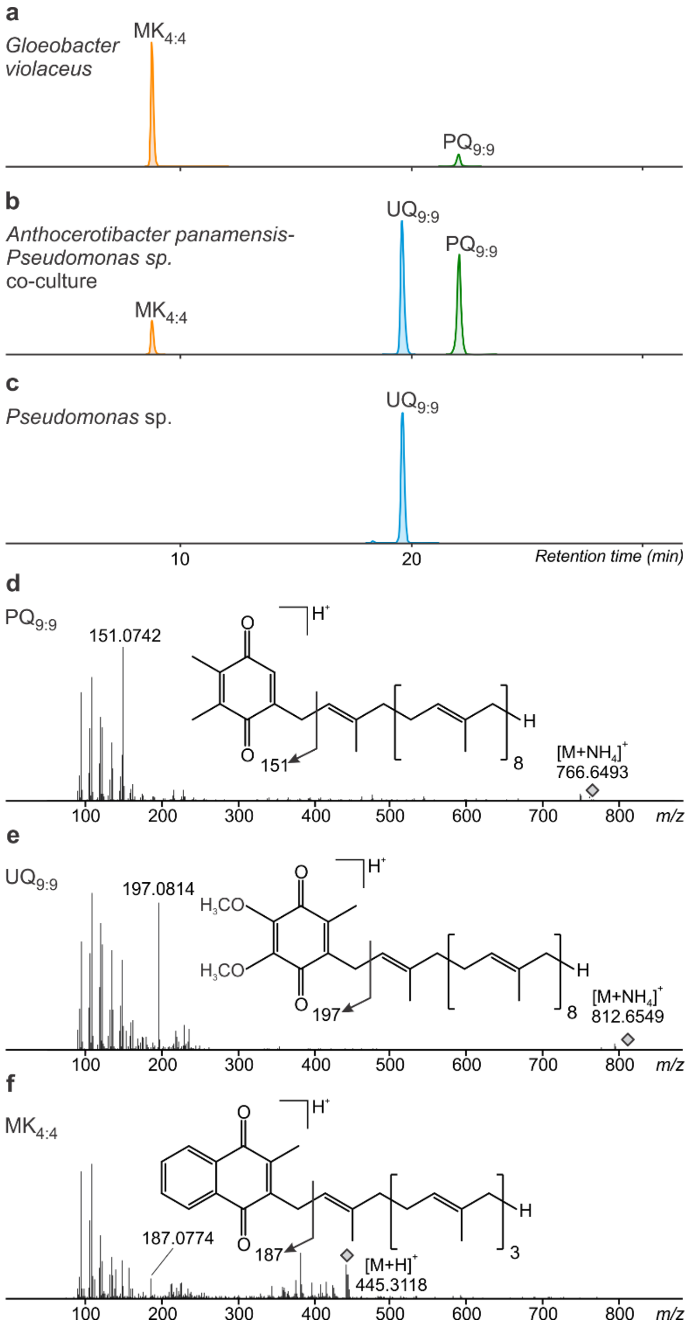

**Fig. S10.** Mass spectra and chromatograms of quinones in basal *Cyanobacteriota*. **a-c**, MS[2] fragmentation spectra and tentative structural characterization of $PQ_{9:9}$ ($NH_4^+$ adduct) and $UQ_{9:9}$ ($NH_4^+$ adduct) from an *Anthocerotibacter panamensis-Pseudomonas* sp. co-culture and $MK_{4:4}$ ($H^+$ adduct) from *Gloeobacter violaceus*. Accurate masses and proposed sum formulas of characteristic fragment ions are shown in Table S1. **d-f**, Composite extracted ion chromatograms of respiratory quinones in extracts of *G. violaceus*, the co-culture, and a *Pseudomonas sp.* isolated from the co-culture.

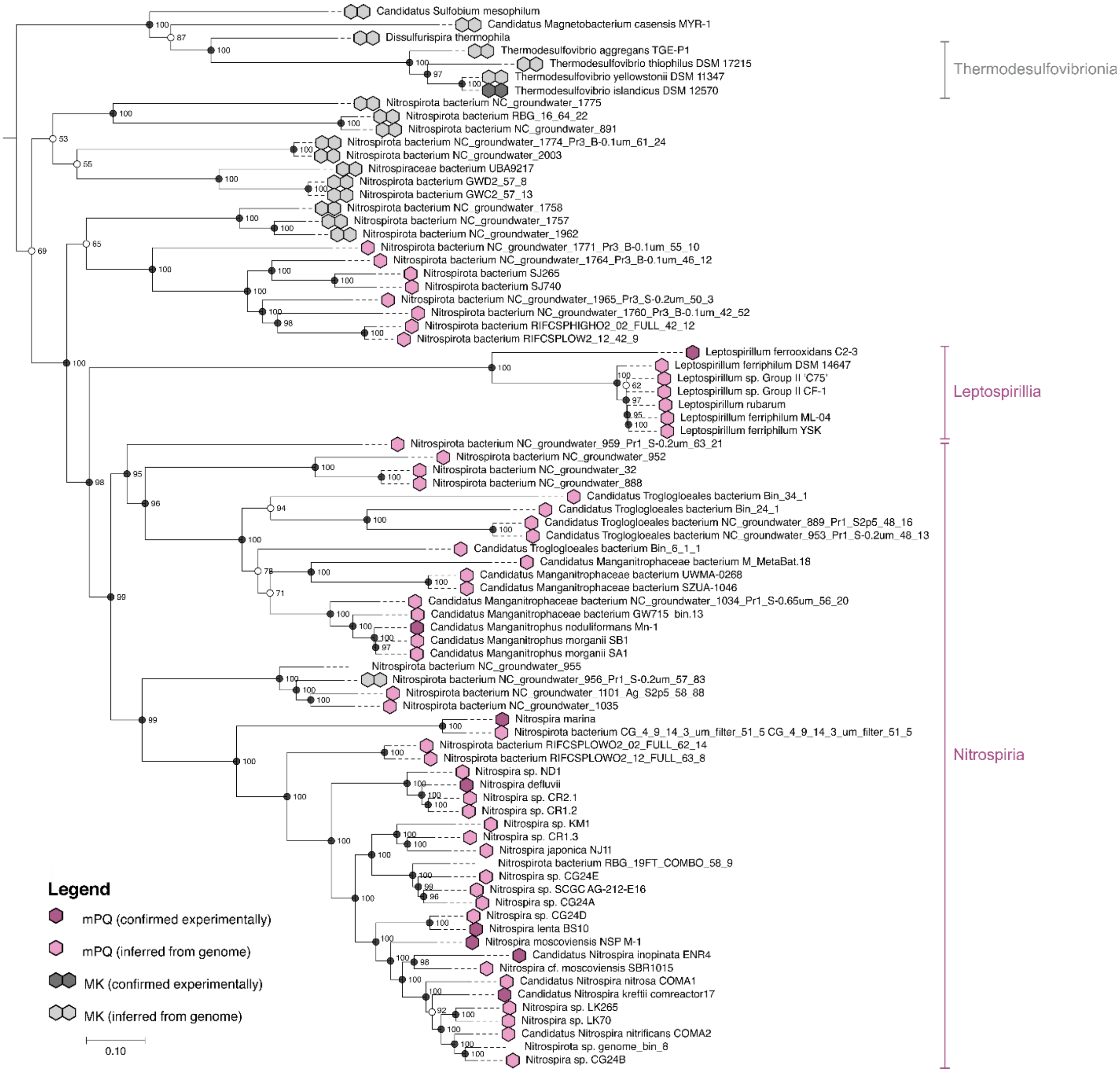

**Fig. S11.** mPQ is present in all studied aerobic *Nitrospirota*. The presence of mPQ (purple) and MK (grey) was experimentally confirmed ($n$=8) or inferred from genomic data ($n$=52) and plotted onto a maximum-likelihood protein phylogeny of 120 single-copy markers from 81 *Nitrospirota* genomes. mPQ biosynthesis could not be inferred from some incomplete genomes. Names of *Nitrospirota* classes are denoted on the right. The scale bar represents 0.1 substitutions per site. Numbers next to nodes indicate ultrafast bootstrap (UFBoot) support. Black circles represent UFBoot support >95%; white circles represent bootstrap support <95%.

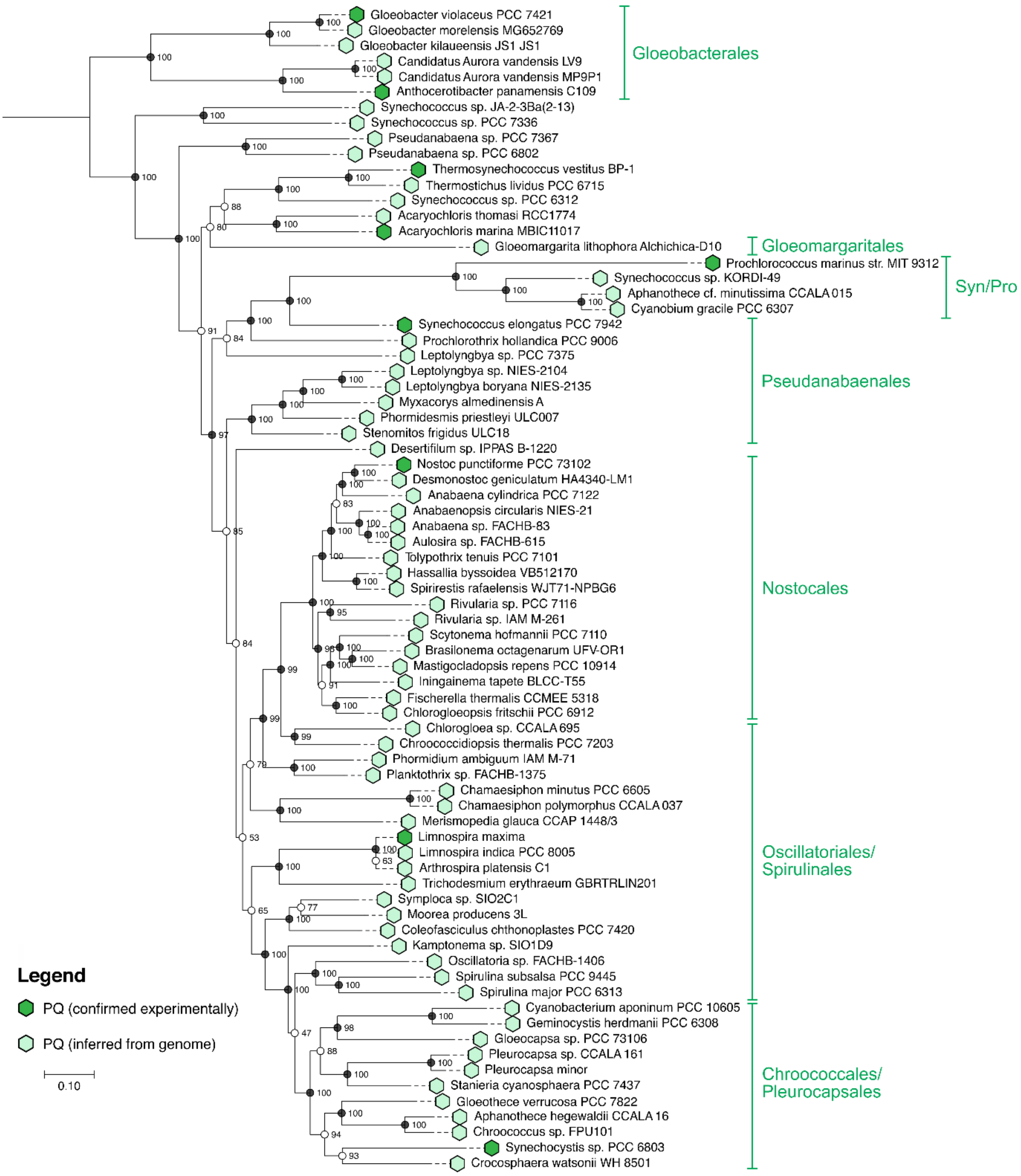

**Fig. S12.** PQ is present in all orders of *Cyanobacteriota*. The presence of PQ was experimentally confirmed (*n*=9) or inferred from genomic data (*n*=66) and plotted onto a maximum-likelihood protein phylogeny of 120 single copy markers from 75 *Cyanobacteriota* genomes. Names of *Cyanobacteriota* orders are denoted on the right. The scale bar represents 0.1 substitutions per site. Numbers next to nodes indicate ultrafast bootstrap (UFBoot) support. Black circles represent UFBoot support >95%; white circles represent bootstrap support <95%.

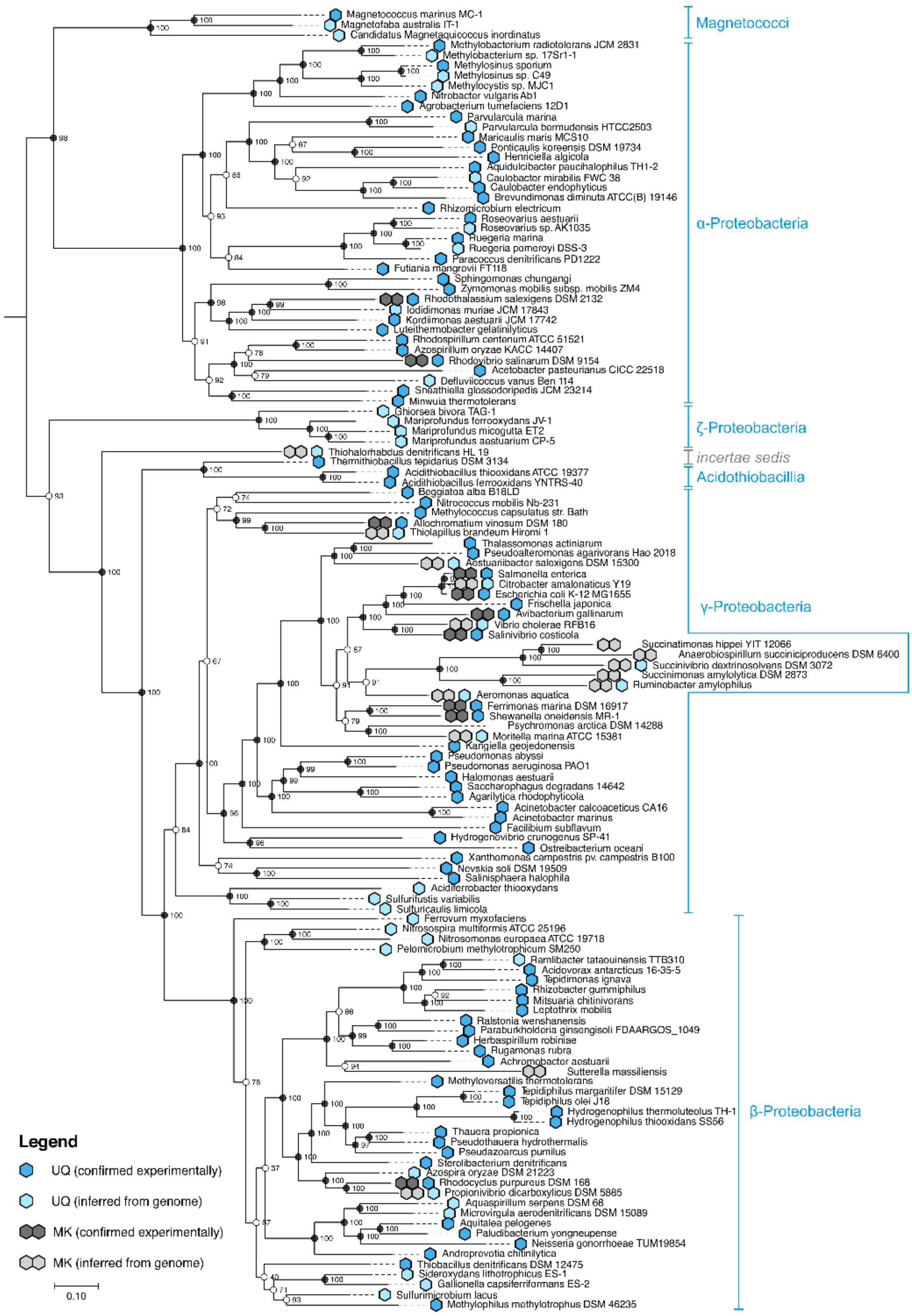

**Fig. S13.** UQ is predominant in all orders of *Pseudomonadota* and MK is present in few, isolated lineages. The presence of UQ and MK was experimentally confirmed (*n*=83) or inferred from genomic data (*n*=45) and plotted onto a maximum-likelihood protein phylogeny of 120 single copy markers from 128 *Pseudomonadota* genomes. Names of *Pseudomonadota* classes are denoted on the right. The scale bar represents 0.1 substitutions per site. Numbers next to nodes indicate ultrafast bootstrap (UFBoot) support. Black circles represent UFBoot support >95%; white circles represent bootstrap support <95%.

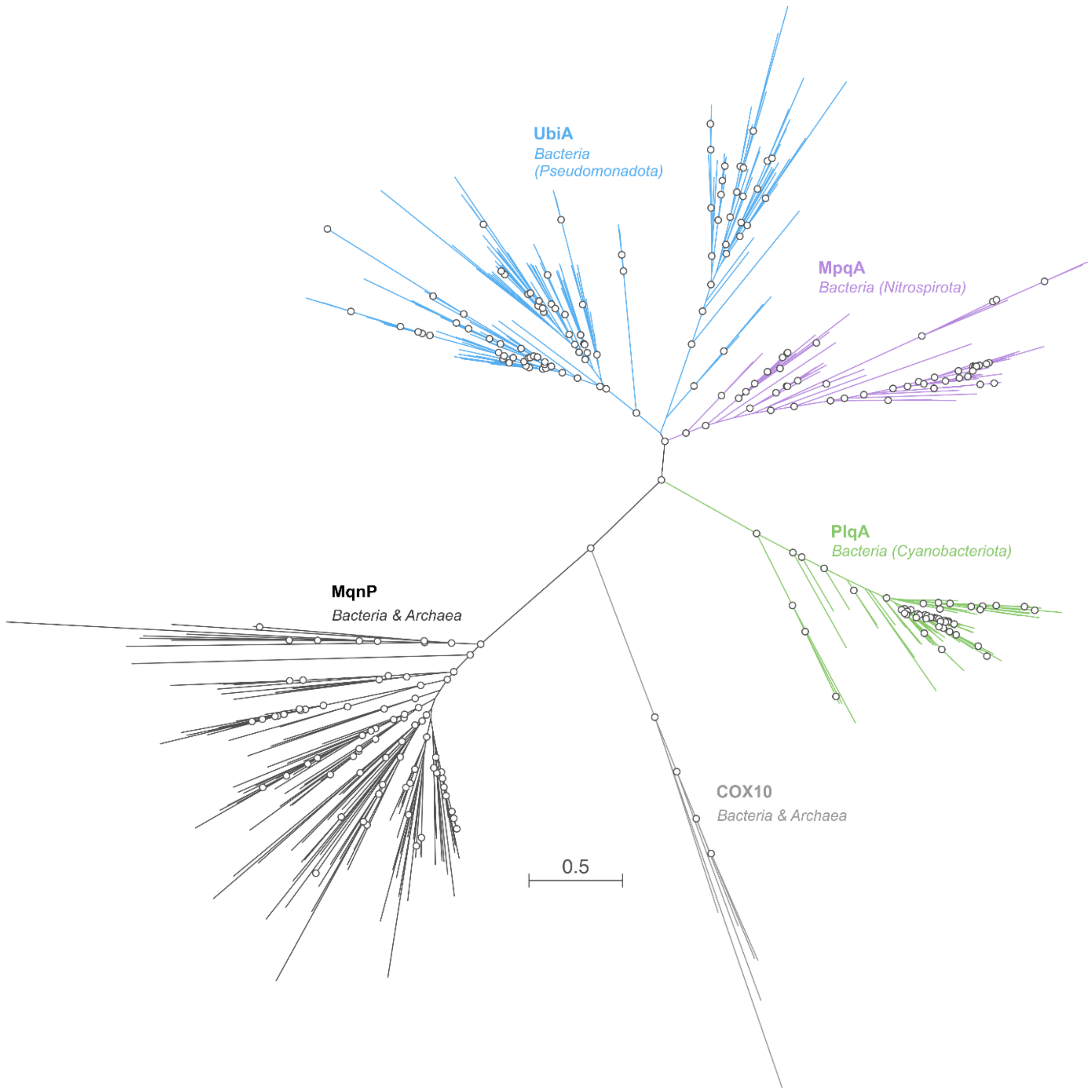

**Fig. S14.** Unrooted phylogenetic tree of UbiA-family prenyltransferase homologs: bacterial prenyltransferase homologs of the HPQ pathways, i.e., UbiA (blue), MpqA (purple), and PlqA (green), as well as archaeal and bacterial prenyltransferase homologs of the $MK_{mqn}$ pathway MqnP (black) and COX10 prenyltransferase homologs (light grey; used as outgroup in main text Fig. 4). Circles indicate ultra-fast bootstrap support ≥90%. The scale bar represents 0.5 amino acid substitutions per site.

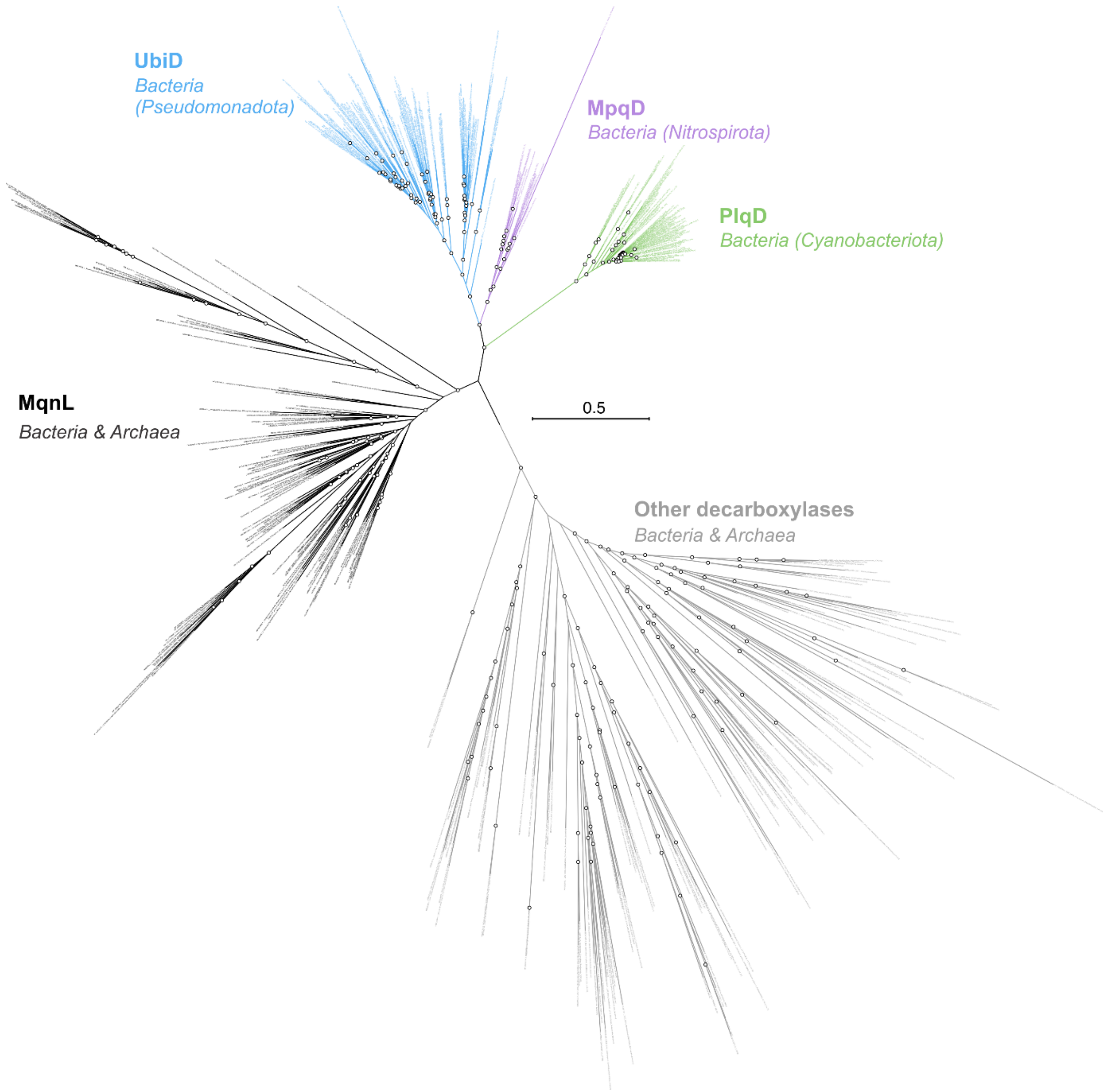

**Fig. S15.** Unrooted phylogenetic tree of UbiD-family decarboxylase homologs: bacterial decarboxylase homologs of the HPQ pathways, i.e., UbiD (blue), MpqD (purple), and PlqD (green), as well as archaeal and bacterial decarboxylase homologs of the $MK_{mqn}$ pathway MqnL (black) and decarboxylases of other functions (light grey; used as outgroup in main text Fig. 4). Circles indicate ultra-fast bootstrap support ≥90%. The scale bar represents 0.5 amino acid substitutions per site.

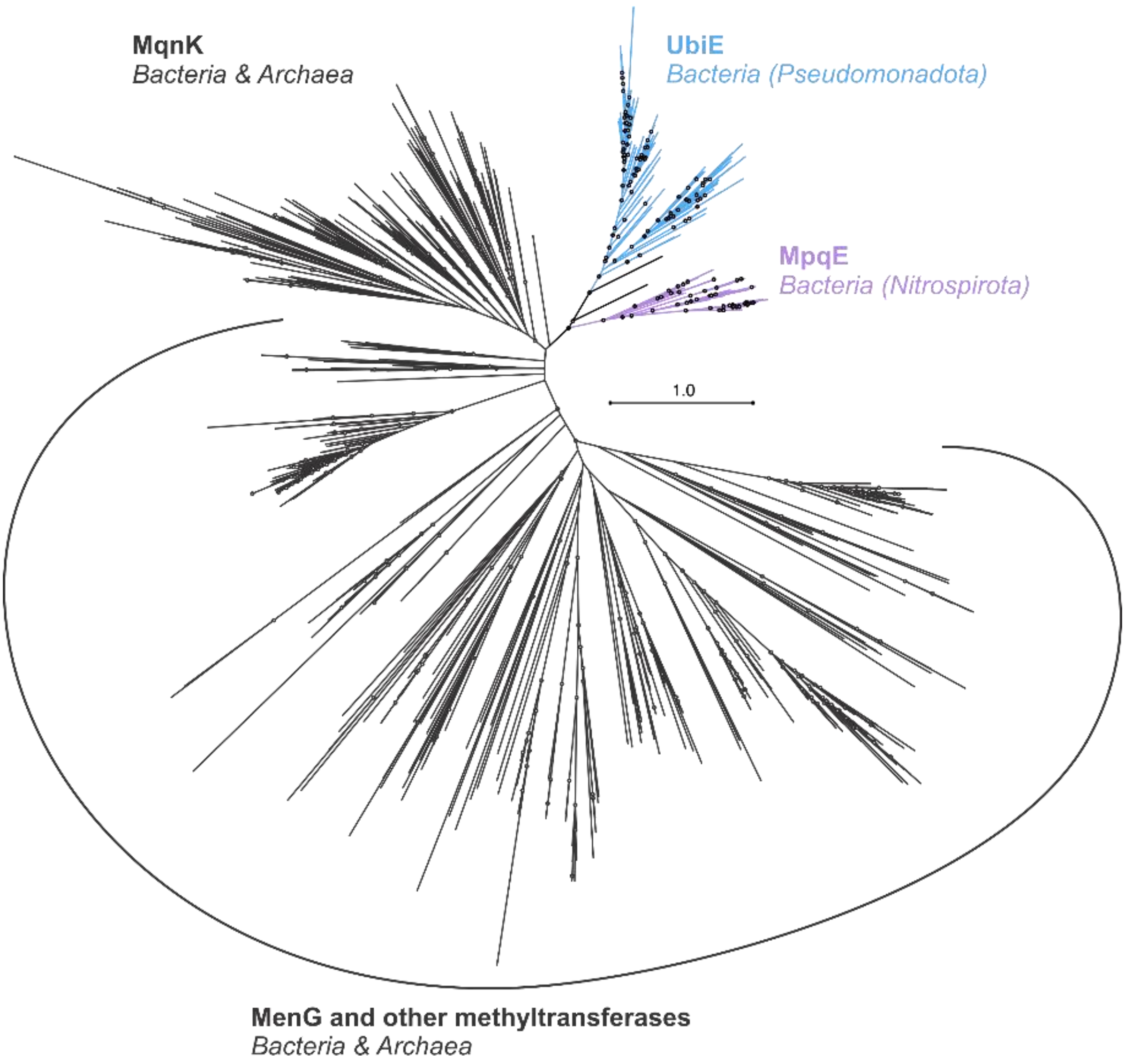

**Fig. S16.** Unrooted phylogenetic tree of quinone methyltransferase homologs: bacterial homologs of HPQ C2 methyltransferases UbiE (blue) and MpqE (purple) as well as archaeal and bacterial homologs of MK C2 methyltransferases MqnK and MenG (black) and other related methyltransferases (used as outgroup in main text Fig. 4). Circles indicate ultra-fast bootstrap support ≥90%. The scale bar represents 1 amino acid substitution per site.

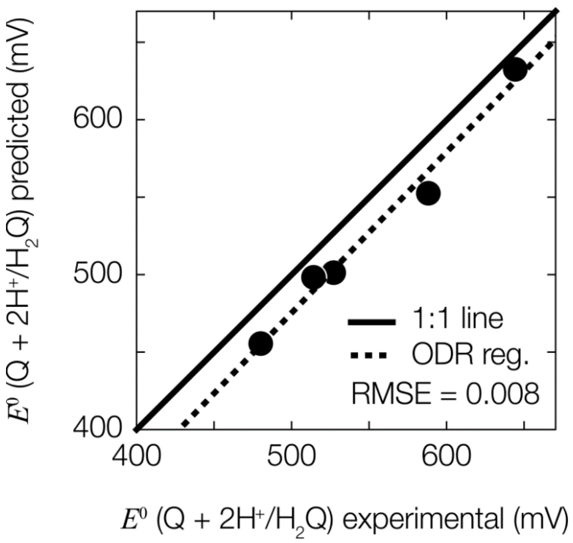

**Fig. S17.** Training set scaling factor results using B3LYP/6-31G(d) functionals and basis sets. Calculated $E^0$ exhibits a strong, positive correlation with experimentally measured $\mathrm{E}^0$ values, as expected. ODR = orthogonal distance regression; RMSE = root mean square error.

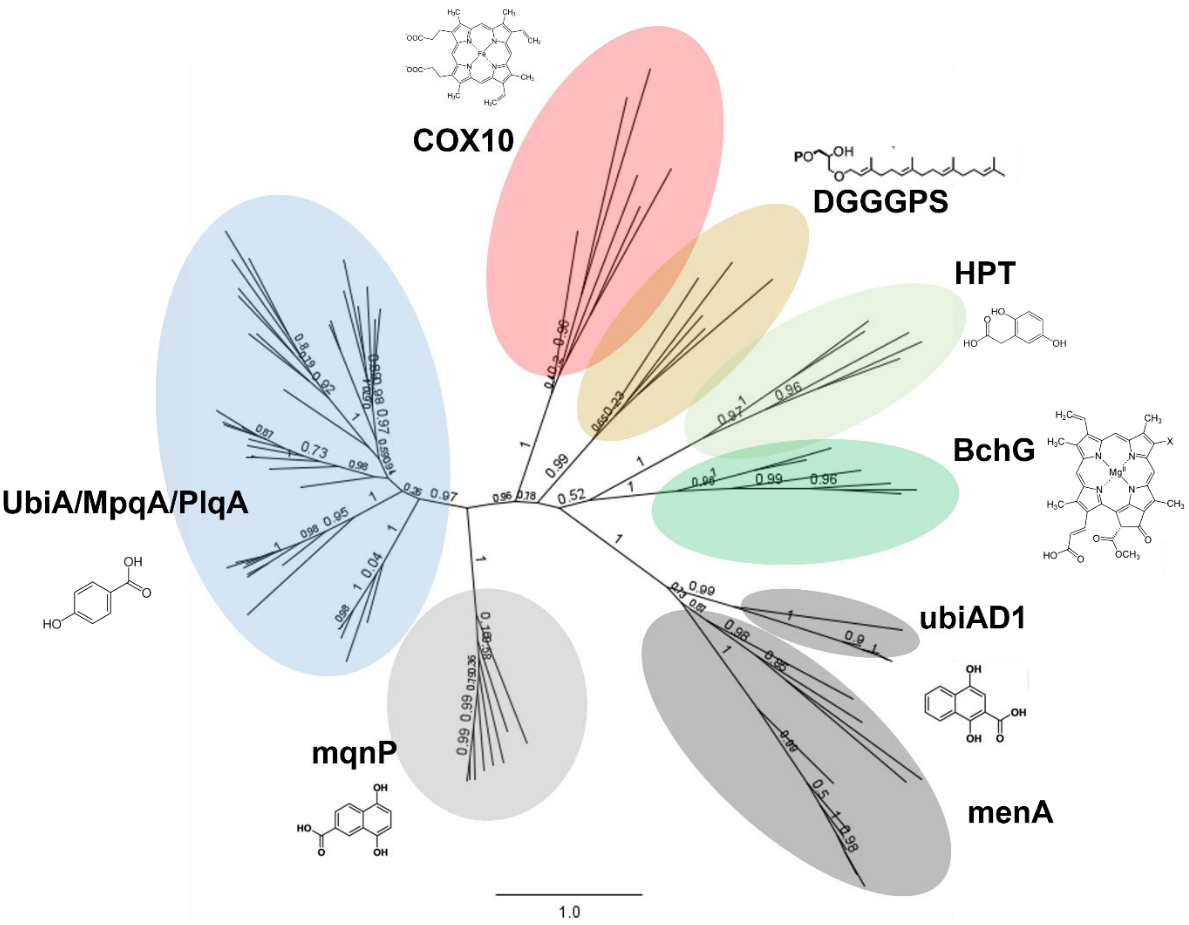

**BchG:** Chlorophyll biosynthesis
**COX10:** Heme biosynthesis
**DGGGPS:** Archaeal membrane lipid biosynthesis
**HPT:** Tocopherol/plant PQ biosynthesis
**MenA/ubiAD1:** Menaquinone biosynthesis (classical)
**MqnP:** Menaquinone biosynthesis (futalosine)
**UbiA/MpqA/PlqA:** UQ/mPQ/PQ biosynthesis

**Fig. S18.** Phylogenetic tree of UbiA family prenyltransferases and their substrates. The prenyltransferases of the HPQ pathways (UQ, PQ, mPQ) are more closely related to the putative prenyltransferase of the $MK_{mqn}$ pathway (MqnP) than to the prenyltransferase of the $MK_{men}$ pathway for menaquinone biosynthesis (MenA). All clusters contain both archaeal and bacterial sequences, except for UbiA/PlqA/MpqA, HPT, and BchG. Numbers on branches indicate ultra-fast bootstrap support. The scale bar represents 1 amino acid substitution per site.

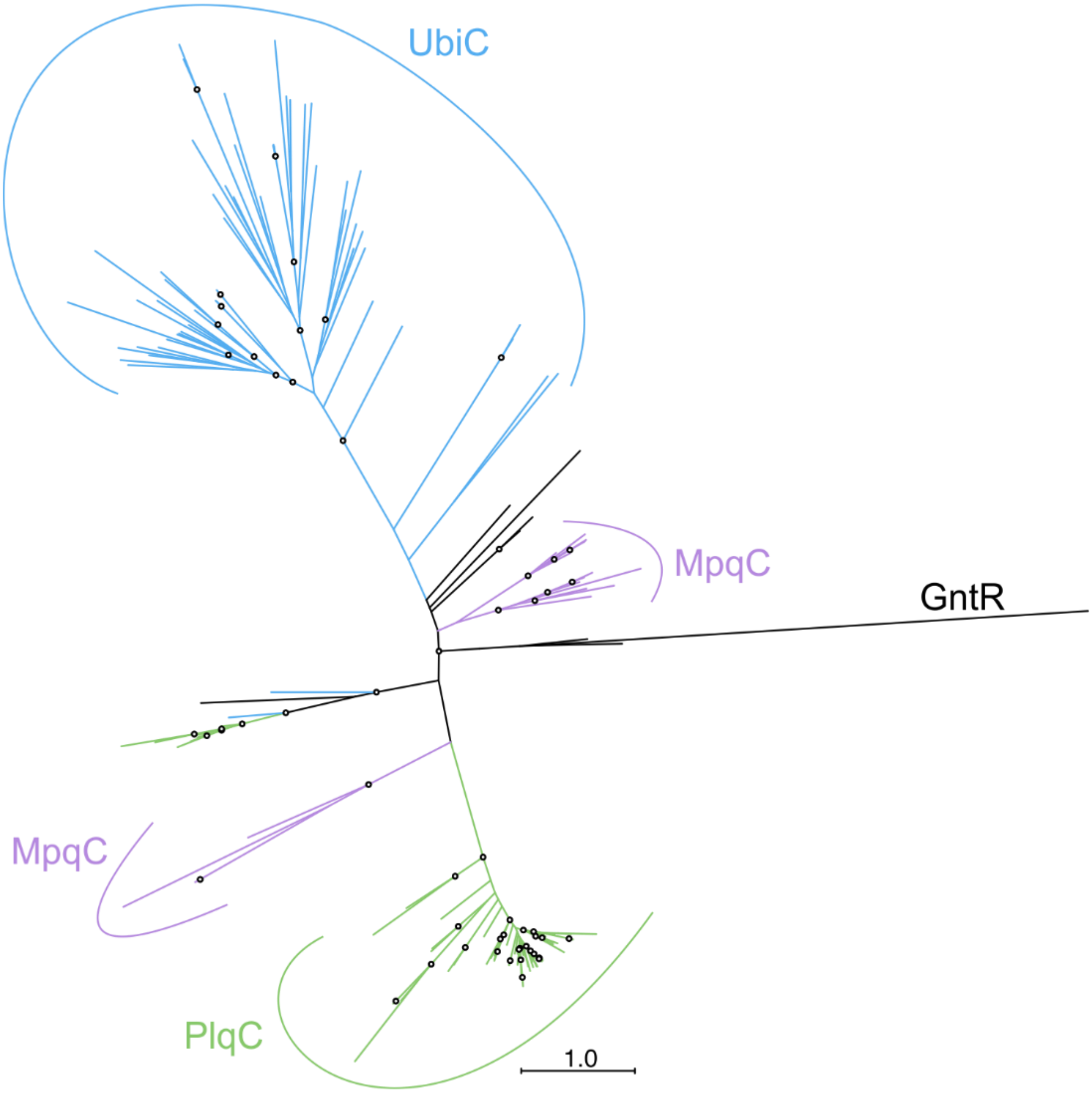

**Fig. S19.** Unrooted phylogenetic tree of chorismate pyruvate-lyase homologs from *Pseudomonadota* (UbiC; blue), aerobic *Nitrospirota* (MpqC; purple), and *Cyanobacteriota* (PlqC; green). Circles indicate ultra-fast bootstrap support ≥90%. The scale bar represents 1 amino acid substitution per site.

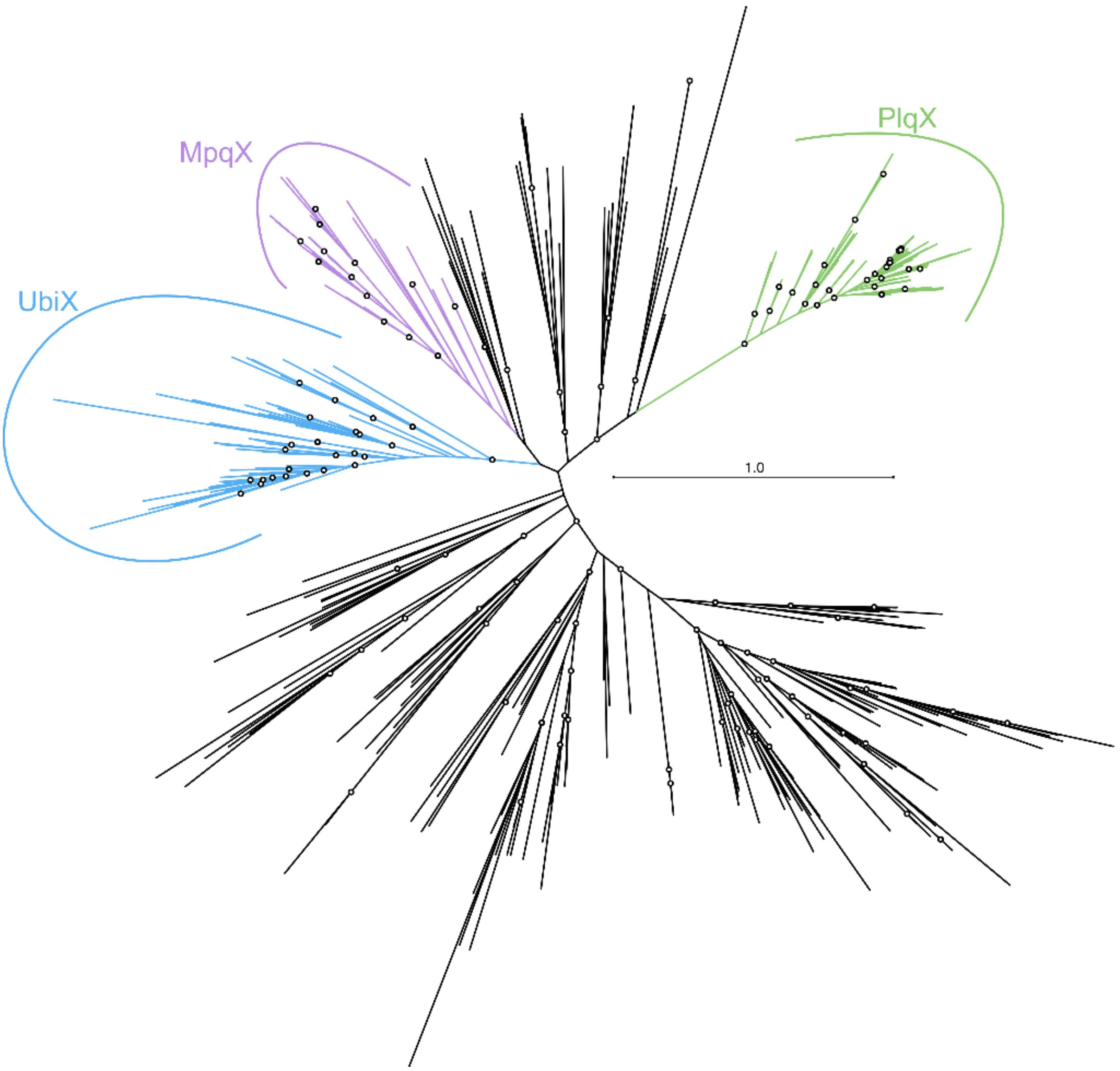

**Fig. S20.** Unrooted phylogenetic tree of flavin prenyltransferase homologs UbiX (blue) and MpqX (purple) as well as other flavin prenyltransferases (black; including putative MqnM homologs). Circles indicate ultra-fast bootstrap support ≥90%. The scale bar represents 1 amino acid substitution per site.

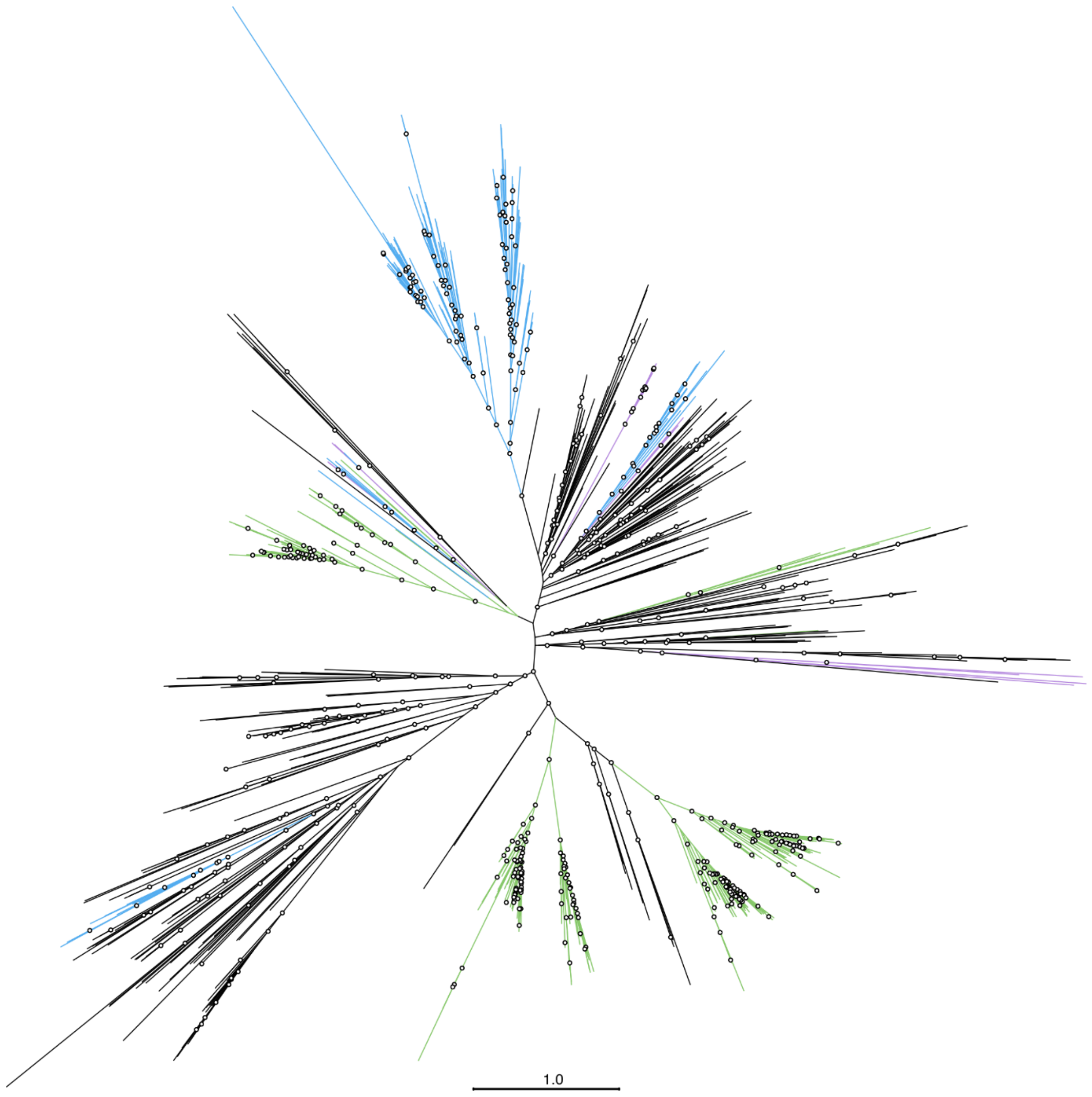

**Fig. S21.** Unrooted phylogenetic tree of putative quinone kinase UbiB homologs in *Pseudomonadota* (blue), *Cyanobacteriota* (green), and aerobic *Nitrospirota* (purple) as well as other quinone kinase homologs (black). Circles indicate ultra-fast bootstrap support ≥90%. The scale bar represents 1 amino acid substitution per site.

### Supplementary Tables

**Table S1.** Observed mass, predicted mass, predicted sum formula, measurement error inferred from predicted and observed masses, and double-bond equivalents (Dbl. eq.) of molecular ions (underlined) and diagnostic fragment ions in representative $MS^2$ spectra of quinones from *Leptospirillum ferrooxidans*, *Nitrospira defluvii*, *N. lenta*, *N. marina*, *N. moscoviensis*, *Ca.* Manganitrophus noduliformans, *Ramlibacter lithotrophicus*, *Thermodesulfovibrio islandicus*, *Gloeobacter violaceus*, *Pseudomonas sp.*, and spinach.

| Compound | Organism | Ion | Observed mass (Da) | Predicted mass (Da) | Predicted formula | Error (ppm) | Error (mDa) | Dbl. eq. |
|---|---|---|---|---|---|---|---|---|
| $MK_{4:1}$ | Spinach | $[M+H]^+$ | 451.3608 | 451.3571 | $C_{31}H_{47}O_2$ | -8.3 | -3.7 | 8.5 |
| | | $[M-C_{19}H_{36}]^+$ | 187.0741 | 187.0754 | $C_{12}H_{11}O_2$ | 6.7 | -1.3 | 7.5 |
| $MK_{4:4}$ | *G. violaceus* | $[M+H]^+$ | 445.3118 | 445.3101 | $C_{31}H_{41}O_2$ | -3.8 | -1.7 | 11.5 |
| | | $[M-C_{19}H_{30}]^+$ | 187.0774 | 187.0754 | $C_{12}H_{11}O_2$ | -10.9 | -2.0 | 7.5 |
| $MK_{6:3}$ | *T. islandicus* | $[M+H]^+$ | 587.4823 | 587.4823 | $C_{41}H_{63}O_2^+$ | -2.8 | -1.6 | 10.5 |
| $MK_{6:4}$ | *T. islandicus* | $[M+H]^+$ | 585.4678 | 585.4666 | $C_{41}H_{61}O_2^+$ | -2.0 | -1.2 | 11.5 |
| $MK_{6:5}$ | *T. islandicus* | $[M+H]^+$ | 583.4516 | 586.4510 | $C_{41}H_{59}O_2^+$ | -1.1 | -0.6 | 12.5 |
| $MK_{6:6}$ | *T. islandicus* | $[M+H]^+$ | 581.4367 | 581.4353 | $C_{41}H_{57}O_2^+$ | -2.4 | -1.4 | 13.5 |
| $MK_{7:4}$ | *T. islandicus* | $[M+H]^+$ | 655.5460 | 655.5449 | $C_{46}H_{71}O_2^+$ | -1.7 | -1.1 | 11.5 |
| $MK_{7:5}$ | *T. islandicus* | $[M+H]^+$ | 653.5305 | 653.5292 | $C_{46}H_{69}O_2^+$ | -2.0 | -1.3 | 12.5 |
| $MK_{7:6}$ | *T. islandicus* | $[M+H]^+$ | 651.5152 | 651.5136 | $C_{46}H_{67}O_2^+$ | -2.5 | -1.6 | 13.5 |
| $MK_{7:7}$ | *T. islandicus* | $[M+H]^+$ | 649.4995 | 649.4979 | $C_{46}H_{65}O_2^+$ | -2.5 | -1.6 | 14.5 |
| $MK_{8:6}$ | *T. islandicus* | $[M+H]^+$ | 721.5931 | 721.5918 | $C_{51}H_{77}O_2^+$ | -1.8 | -1.3 | 13.5 |
| $MK_{8:7}$ | *T. islandicus* | $[M+H]^+$ | 719.5761 | 719.5762 | $C_{51}H_{75}O_2^+$ | -0.1 | 0.0 | 14.5 |
| $MK_{8:8}$ | *T. islandicus* | $[M+NH_4]^+$ | 734.5873 | 734.5871 | $C_{51}H_{76}NO_2^+$ | -0.3 | -0.2 | 14.5 |
| | | $[M-C_{39}H_{65}N]^+$ | 187.0750 | 187.0754 | $C_{12}H_{11}O_2^+$ | 1.9 | 0.4 | 7.5 |
| $mPQ_{7:6}$ | *L. ferrooxidans* | $[M+NH_4]^+$ | 646.5460 | 646.5558 | $C_{44}H_{72}NO_2^+$ | 15.1 | 9.8 | 9.5 |
| | | $[M-C_{34}NH_{59}]^+$ | 165.0898 | 165.0910 | $C_{10}H_{13}O_2^+$ | 7.3 | 1.2 | 4.5 |
| $mPQ_{7:7}$ | *N. inopinata* | $[M+NH_4]^+$ | 644.5396 | 644.5401 | $C_{44}H_{70}NO_2^+$ | 0.8 | 0.5 | 10.5 |
| | | $[M-C_{34}NH_{57}]^+$ | 165.0899 | 165.0910 | $C_{10}H_{13}O_2^+$ | 6.7 | 1.1 | 4.5 |
| $mPQ_{8:7}$ | *N. inopinata* | $[M+NH_4]^+$ | 714.6158 | 714.6184 | $C_{49}H_{80}NO_2^+$ | 3.6 | 2.6 | 10.5 |
| | | $[M-C_{39}NH_{67}]^+$ | 165.0905 | 165.0910 | $C_{10}H_{13}O_2^+$ | 3.1 | 0.5 | 4.5 |
| $mPQ_{8:8}$ | *N. marina* | $[M+NH_4]^+$ | 712.6065 | 712.6072 | $C_{49}H_{78}NO_2^+$ | -5.3 | -3.8 | 11.5 |
| | | $[M-C_{39}NH_{65}]^+$ | 165.0916 | 165.0910 | $C_{10}H_{13}O_2^+$ | -3.6 | -0.6 | 4.5 |
| $mPQ_{9:8}$ | *N. marina* | $[M+NH_4]^+$ | 782.6858 | 782.6810 | $C_{54}H_{88}NO_2^+$ | -6.2 | -4.8 | 11.5 |
| | | $[M-C_{44}NH_{75}]^+$ | 165.0924 | 165.0910 | $C_{10}H_{13}O_2^+$ | -8.4 | -1.4 | 4.5 |
| $mPQ_{9:9}$ | *N. marina* | $[M+H]^+$ | 763.6425 | 763.6388 | $C_{54}H_{83}O_2^+$ | -4.9 | -3.7 | 13.5 |
| | | $[M-C_{44}H_{70}]^+$ | 165.0915 | 165.0910 | $C_{10}H_{13}O_2^+$ | -3.0 | -0.5 | 4.5 |
| | | $[M-C_{46}H_{70}O_2]^+$ | 109.1021 | 109.1012 | $C_8H_{13}^+$ | -8.5 | -0.9 | 2.5 |
| | | $[M-C_{44}H_{68}O_2]^+$ | 135.1178 | 135.1168 | $C_{10}H_{15}^+$ | -7.2 | -1.0 | 3.5 |
| | | $[M-C_{40}H_{62}O_2]^+$ | 189.1644 | 189.1638 | $C_{14}H_{21}^+$ | -3.3 | -0.6 | 4.5 |
| | | $[M-C_{42}H_{64}O_2]^+$ | 215.1811 | 215.1794 | $C_{16}H_{23}^+$ | -7.8 | -1.7 | 5.5 |
| | | $[M-C_{39}H_{64}]^+$ | 231.1390 | 231.1380 | $C_{15}H_{19}O_2^+$ | -4.5 | -1.0 | 6.5 |
| | | $[M-C_{35}H_{54}O_2]^+$ | 257.2265 | 257.2264 | $C_{19}H_{29}^+$ | -0.5 | -0.1 | 5.5 |
| | | $[M-C_{33}H_{58}O_2]^+$ | 285.2591 | 285.2577 | $C_{21}H_{33}^+$ | -5.0 | -1.4 | 5.5 |
| | | $[M-C_{31}H_{62}O_2]^+$ | 313.2921 | 313.2890 | $C_{23}H_{37}^+$ | -10.0 | -3.1 | 5.5 |
| $mPQ_{10:9}$ | *N. defluvii* | $[M+NH_4]^+$ | 850.7428 | 850.7436 | $C_{59}H_{96}NO_2^+$ | 0.9 | 0.8 | 12.5 |
| | | $[M-C_{49}NH_{83}]^+$ | 165.0887 | 165.0910 | $C_{10}H_{13}O_2^+$ | 14.0 | 2.3 | 4.5 |
| $mPQ_{10:10}$ | *N. marina* | $[M+H]^+$ | 831.6991 | 831.7014 | $C_{59}H_{91}O_2^+$ | 2.7 | 2.3 | 14.5 |
| | | $[M-C_{49}H_{78}]^+$ | 165.0894 | 165.0910 | $C_{10}H_{13}O_2^+$ | 9.7 | 1.6 | 4.5 |
| $mPQ_{12:10}$ | *M. noduliformans* | $[M+NH_4]^+$ | 988.8864 | 988.8844 | $C_{69}H_{114}NO_2^+$ | -2.0 | -2.0 | 13.5 |
| | | $[M-C_{59}H_{101}N]^+$ | 165.0987 | 165.0910 | $C_{10}H_{13}O_2^+$ | -46.6 | -7.7 | 4.5 |
| $mPQ_{12:11}$ | *M. noduliformans* | $[M+NH_4]^+$ | 986.8697 | 986.8688 | $C_{69}H_{112}NO_2^+$ | -1.0 | -0.9 | 14.5 |
| | | $[M-C_{59}H_{99}N]^+$ | 165.0920 | 165.0910 | $C_{10}H_{13}O_2^+$ | -6.0 | -1.0 | 4.5 |
| $mPQ_{13:11}$ | *M. noduliformans* | $[M+NH_4]^+$ | 1056.9474 | 1056.9470 | $C_{74}H_{122}NO_2^+$ | -0.4 | -0.4 | 14.5 |
| | | $[M-C_{64}H_{109}N]^+$ | 165.0910 | 165.0910 | $C_{10}H_{13}O_2^+$ | 0 | 0 | 4.5 |
| $mPQ_{13:12}$ | *M. noduliformans* | $[M+NH_4]^+$ | 1054.9349 | 1054.9314 | $C_{74}H_{120}NO_2^+$ | -3.4 | -3.5 | 15.5 |
| | | $[M-C_{64}H_{107}N]^+$ | 165.0899 | 165.0910 | $C_{10}H_{13}O_2^+$ | 6.7 | 1.1 | 4.5 |

| | | | | | | | | |
|---|---|---|---|---|---|---|---|---|
| $mPQ_{13:13}$ | *M. noduliformans* | $[M+NH_4]^+$ | 1052.9201 | 1052.9157 | $C_{74}H_{118}NO_2^+$ | -4.2 | -4.4 | 16.5 |
| $PQ_{9:9}$ | Spinach | $[M+NH_4]^+$ | 766.6567 | 766.6497 | $C_{53}H_{84}NO_2^+$ | -9.2 | -7.0 | 12.5 |
| | | $[M-C_{44}H_{73}N]^+$ | 151.0759 | 151.0754 | $C_9H_{11}O_2^+$ | -3.6 | -0.5 | 4.5 |
| | | $[M-C_{45}H_{71}NO_2]^+$ | 109.1021 | 109.1012 | $C_8H_{13}^+$ | -8.5 | -0.9 | 2.5 |
| | | $[M-C_{43}H_{69}NO_2]^+$ | 135.1184 | 135.1168 | $C_{10}H_{15}^+$ | -11.6 | -1.6 | 3.5 |
| | | $[M-C_{41}H_{65}NO_2]^+$ | 163.1476 | 163.1481 | $C_{12}H_{19}^+$ | 3.2 | 0.5 | 3.5 |
| | | $[M-C_{39}H_{63}NO_2]^+$ | 189.1656 | 189.1638 | $C_{14}H_{21}^+$ | -9.6 | -1.8 | 4.5 |
| | | $[M-C_{37}H_{61}NO_2]^+$ | 215.1790 | 215.1794 | $C_{16}H_{23}^+$ | 2.0 | 0.4 | 5.5 |
| | | $[M-C_{37}H_{63}N]^+$ | 245.1560 | 245.1536 | $C_{16}H_{21}O_2^+$ | -9.8 | -2.4 | 6.5 |
| | | $[M-C_{34}H_{59}N]^+$ | 285.1840 | 285.1849 | $C_{19}H_{25}O_2^+$ | 3.2 | 0.9 | 7.5 |
| $UQ_{8:8}$ | *R. lithotrophicus* | $[M+NH_4]^+$ | 744.5940 | 744.5925 | $C_{49}H_{78}NO_4^+$ | -2.0 | -1.5 | 11.5 |
| | | $[M-C_{49}H_{81}N]^+$ | 197.0798 | 197.0808 | $C_{10}H_{13}O_4^+$ | 5.3 | 1.0 | 4.5 |
| $UQ_{9:9}$ | *Pseudomonas* sp. | $[M+NH_4]^+$ | 812.6554 | 812.6551 | $C_{54}H_{86}NO_4^+$ | -0.3 | -0.3 | 12.5 |
| | | $[M-C_{44}H_{73}N]^+$ | 197.0810 | 197.0808 | $C_{10}H_{13}O_4^+$ | -0.8 | -0.2 | 4.5 |
| $UQ_{10:10}$ | Spinach | $[M+NH_4]^+$ | 880.7253 | 880.7177 | $C_{59}H_{94}NO_4^+$ | -8.6 | -7.6 | 13.5 |
| | | $[M-C_{49}H_{81}N]^+$ | 197.0817 | 197.0808 | $C_{10}H_{13}O_4^+$ | -4.4 | -0.9 | 4.5 |

**Table S2.** Potential mPQ biosynthesis gene homologs in cultivated *Nitrospirota*. Homologs assayed *in vivo* in *E. coli* are highlighted in bold.

| | *ubiC*-like Chorismate lyase *mpqC* | *xanB2*-like Chorismate lyase | *ubiA*-like Prenyltransferase *mpqA* | *ubiD*-like Decarboxylase *mpqD* | *ubiX*-like Decarboxylase *mpqX* | *ubiB*-like Kinase *mpqB* | *ubiE*-like Methyltransferase *mpqE* | *sll0418*-like Methyltransferase *mpqQ* |
|---|---|---|---|---|---|---|---|---|
| *Nitrospira defluvii* | - | - | WP_083817284.1 | - | - | - | WP_013249320.1 | - |
| *Nitrospira* sp. ND1 | - | - | WP_080878760.1 | - | - | - | WP_080877778.1 | - |
| *Nitrospira japonica* NJ1 | - | - | WP_080886482.1 | - | - | - | WP_080886247.1 | WP_080888637.1 |
| *Ca.* Nitrospira kreftii | - | - | QPD05446.1 | - | - | QPD05554.1 | QPD05757.1 | QPD04045.1 |
| *Nitrospira moscoviensis* M1 | - | - | **WP_053378563.1** | - | - | - | WP_083447688.1 | - |
| *Ca.* Nitrospira nitrosa COMA1 | - | - | WP_090742531.1 | - | - | - | WP_090745869.1 | WP_090749253.1 |
| *Nitrospira inopinata* ENR4 | - | WP_062481980.1 | WP_062484573.1 | - | - | WP_197549219.1 | **WP_082633614.1** | **WP_062482398.1** |
| *Ca.* Nitrospira nitrificans COMA2 | **WP_090896071.1** | - | WP_090900288.1 | - | - | WP_090896854.1 | WP_090897166.1 | WP_090895890.1 |
| *Nitrospira lenta* BS10 | - | WP_121989121.1 | WP_121988012.1 | - | - | - | WP_121988272.1 | - |
| *Nitrospira marina* 295 | - | - | 2597410107 | - | - | - | 2597410471 | - |
| *Leptospirillum rubarum* | WP_014962020.1 | - | EAY58208.1 | - | - | - | EAY58205.1 | - |
| *Leptospirillum ferriphilum* ML-04 | WP_036080762.1 | - | WP_038506322.1 | - | - | WP_077305007.1 | WP_036080765.1 | - |
| *Leptospirillum ferrooxidans* C2-3 | **WP_014450279.1** | - | **WP_014450278.1** | - | - | **WP_014450220.1** | **WP_041774405.1** | - |
| *Leptospirillum ferrodiazotrophum* | EES52035.1 | - | EES52034.1 | - | - | EES52153.1 | EES52037.1 | - |
| *Ca.* Manganitrophus noduliformans | **WP_168061792.1** | - | **WP_168061790.1** | **WP_168061789.1** | **WP_168059374.1** | WP_168059066 | **WP_168059970.1** | - |

**Table S3.** Results of redox potential calculations (scaled to experimental data) showing that mPQ has a redox potential ($E^0$, Q/$H_2$Q) similar to, but lower than, PQ. Redox potential is dependent on chain length; results for different chain lengths can be found in Supplementary Datafile S4. Further, calculations for 1,4-benzoquinone analogs show that addition of methyl or methoxy functional groups leads to progressively lower $E^0$. Consequently, $E^0$ values of UQ (R2, R3=methoxy, R6=methyl) and mPQ (R2, R3, R6=methyl) are lower than PQ (R2, R3=methyl). $E^0$ of PQ, mPQ, and UQ are higher than MK. The full dataset is available in Supplementary Datafile S4. RMSE, $E^0$ root mean square error.

| **Compound** | **Quinone functional groups** | **$E^0$, Q/$H_2$Q** | **RMSE** |
|---|---|---|---|
| $MK_{9:9}$ | - | 0.364 | 0.008 |
| $PQ_{9:9}$ | dimethyl | 0.551 | 0.008 |
| $mPQ_{9:9}$ | trimethyl | 0.517 | 0.008 |
| $UQ_{9:9}$ | dimethoxy, methyl | 0.480 | 0.008 |
| 2-methyl-1,4-benzoquinone | methyl | 0.651 | 0.008 |
| 2,3-dimethyl-1,4-benzoquinone | dimethyl | 0.574 | 0.008 |
| trimethyl-1,4-benzoquinone | trimethyl | 0.525 | 0.008 |
| tetramethyl-1,4-benzoquinone | tetramethyl | 0.481 | 0.008 |
| 2-methoxy-1,4-benzoquinone | methoxy | 0.627 | 0.008 |
| 2,6-dimethoxy-1,4-benzoquinone | dimethoxy | 0.522 | 0.008 |

**Table S4.** Sequence motif near the [2Fe-2S] cluster of the Rieske protein of Rieske/cytochrome *b* complexes in selected *Nitrospirota*. The 'SY' and 'SF' motifs are associated nearly exclusively with Rieske proteins interacting with HPQs in high-potential ETCs(88). The GY motif is characteristic for Rieske proteins interacting with LPQs. Multiple motifs are listed for species that contain multiple Rieske/cytochrome *b* complexes. Quinone inference is based on Supplementary Datafile S3. Asterisks denote cases in which no sequence motif could be identified: * Genes for the Rieske protein and cytochrome $b_6$ are not followed by a subunit IV with the conserved "PEWY" motif or a variant thereof; ** no subunit IV identified.

| Species | Sequence motif | Quinone | Source |
|---|---|---|---|
| *Leptospirillum ferrooxidans* | SY, SY | mPQ | Ref. (88) |
| *Leptospirillum ferrodiazotrophum* | SY, SY | mPQ | Ref. (88) |
| *Leptospirillum rubarum* | SY, SY | mPQ | Ref. (88) |
| *Nitrospira defluvii* | SY, SF | mPQ | Ref. (88) |
| *Nitrospira moscoviensis* NSP M-1 | SY, SF, GY | mPQ | This study |
| *Nitrospira inopinata* ENR4 | SY | mPQ | This study |
| *Nitrospira lenta* BS10 | SY | mPQ | This study |
| *Ca.* Nitrospira kreftii comreactor17 | SY | mPQ | This study |
| *Ca.* Nitrospira nitrificans COMA2 | SY | mPQ | This study |
| *Ca.* Nitrospira nitrosa COMA1 | SY | mPQ | This study |
| *Manganitrophus morganii* SB1 | GY, SF, SY, SF | mPQ | This study |
| *Manganitrophus noduliformans* Mn-1 | SY | mPQ | This study |
| *Nitrospirota* bacterium RIFCSPHIGHO2_02_FULL_42_12 | SY | mPQ | This study |
| *Nitrospirota* bacterium RIFCSPLOW2_12_42_9 | * | mPQ | This study |
| *Ca.* Manganitrophus sp. isolate GAC1' | SY, SF, GY | mPQ | This study |
| *Thermodesulfovibrio yellowstonii* | GY | MK | This study |
| *Thermodesulfovibrio islandicus* | GY | MK | This study |
| *Ca.* Magnetobacterium casensis MYR-1 | ** | MK | This study |
| *Dissulfurispira thermophila* | GY | MK | This study |
| Nitrospirae bacterium GWC2_57_13 | GY, GF, SF | MK | This study |
| Nitrospirae bacterium RBG_16_64_22 | SF | MK | This study |

**Table S5.** Quinone composition of the clades most closely related to *Cyanobacteriota*, *Nitrospirota*, and *Pseudomonadota* and their quinone biosynthesis pathways predicted from the presence of key biosynthetic genes (see Materials & Methods). Please note that taxonomic assignments differ between GTDB and NCBI taxonomies and "*Candidatus* Melainobacteriota" may variably include *Vampirovibrionophyceae* and "*Candidatus* Sericytochromatia". The full dataset is available in Supplementary Datafile S3. Asterisk: *Nitrospirota* were assigned into aerobic and anaerobic based on physiological evidence or presence of mPQ vs MK pathway genes (no pathway identified for 6/84 *Nitrospirota* genomes).

| | **Quinone in phenotype** | **Pathway predicted from genome** | **# of genomes with MK pathway (of total tested)** |
|---|---|---|---|
| **Closest relatives to Cyanobacteriota** | | | |
| *Cyanobacteriota* | PQ/PhQ | PQ/$MK_{men}$ | 73/74 |
| *Ca. S*ericytochromatia/Melainobacteriota | Not tested | $MK_{mqn}$ | 2/15 |
| Vampirovibrionophyceae/Melainobacteriota | Not tested | $MK_{mqn}$ | 5/6 |
| *Ca.* Margulisbacteria | Not tested | $MK_{mqn}$ | 2/18 |
| **Closest relatives to Nitrospirota** | | | |
| Aerobic *Nitrospirota** | mPQ | mPQ | 0/61 |
| Anaerobic *Nitrospirota** | MK | $MK_{mqn}$ | 18/18 |
| *Acidobacteriota* | MK | $MK_{mqn}$ | 6/7 |
| *Methylomirabilota* | MK | $MK_{mqn}$ | 5/6 |
| *Nitrospinota* | Not tested | $MK_{mqn}$ | 12/12 |
| **Closest relatives to Pseudomonadota** | | | |
| *Pseudomonadota* | UQ/MK | UQ/$MK_{men}$ | 22/130 |
| *Campylobacteriota* | MK | $MK_{mqn}$ | 10/10 |
| *Myxococcota* | MK | $MK_{mqn}$ | 8/11 |
| *Bdellovibrionota* | Not tested | $MK_{mqn}$ | 7/8 |
| *Desulfobacterota* | MK | $MK_{mqn}$ | 5/10 |

**Table S6.** Strains used in this study.

| | **Strain/mutant** | **Relevant genotype** | **Origin** |
|---|---|---|---|
| *Escherichia coli* | MG1655 | Wild type | Laboratory collection |
| | Δ*ubiAc* | MG1655 Δ*ubiA*::cat cured with pCP20 | Kazemzadeh et al.(89) |
| | Δ*ubiB* | as MG1655 but *ubiB*::cat, *cat* gene insertion at *Nru*I site (842pb) | Pelosi et al.(11) |
| | Δ*ubiCc* | Δ*ubiC* cured with pCP20 | Kazemzadeh et al.(89) |
| | Δ*ubiD* | as MG1655 but Δ*ubiD*::cat | Pelosi et al.(11) |
| | Δ*ubiX* | as MG1655 but Δ*ubiX*::kan | Pelosi et al.(11) |
| | Δ*ubiIF* | as MG1655 but Δ*ubiIc* Δ*ubiF*::Kan | Hajj-Chehade et al.(87) |
| | Δ*ubiIFc* | Δ*ubiIF* cured with pCP20 | This study |
| | Δ*ubiIFc* Δ*ubiE* | Δ*ubiIFc* +P1/Δ*ubiE*::Kan | This study |
| | Δ*ubiIHFc* | Δ*ubiIHF* cured with pCP20 | Arias-Cartin et al.(90) |
| *Anthocerotibacter panamensis-Pseudomonas sp. co-culture* | - | | Fay-Wey Li (Cornell, USA) (91) |
| *Gloeobacter violaceus* | - | | L.L. Jahnke, M.N. Parenteau (NASA Ames, USA) |
| *Leptospirillum ferrooxidans* | DSM 2705/C2-3 | | DSMZ |
| *Ca.* Manganitrophus noduliformans/*Ramlibacter lithotrophicus co-culture* | Mn1/RBP-1 | | J.R. Leadbetter (5) |
| *Ramlibacter lithotrophicus* | RBP-1 | | J.R. Leadbetter (5) |
| *Nitrospira defluvii* | A17 | | E. Spieck (92) |
| *Nitrospira inopinata* | ENR4 | | S. Lücker (93) |
| *Nitrospira kreftii* | comreactor17 | | S. Lücker (4) |
| *Nitrospira lenta* | BS10 | | E. Spieck (92) |
| *Nitrospira marina* | Nb-295 | | E. Spieck (94) |
| *Nitrospira moscoviensis* | M1 | | E. Spieck (1) |
| *Pseudomonas sp.* | - | | This study |
| *Thermodesulfovibrio islandicus* | DSM 12570 | | DSMZ |

**Table S7.** Primers for *ubiE* and *ubiF* used in this study.

| Primer | Sequence |
|---|---|
| UbiF5 | ATGCAGGGCGCATAGTGTAA |
| UbiF3 | CCGCGTCTTATCCGACCTAC |
| UbiE5 | CGGTCGGTTCCGGAGCAGCCG |
| UbiE3 | GCCGTTTTCAGCGCGGGTGAGC |
